# A modular, adaptable, and accessible implant kit for chronic electrophysiological recordings in rats

**DOI:** 10.1101/2025.02.27.640674

**Authors:** Raquel J. Ibáñez Alcalá, Andrea Y. Macias, Cory N. Heaton, Ricardo Sosa Jurado, Alexis A. Salcido, Neftali F. Reyes, Serina A. Batson, Luis D. Davila, Dirk W. Beck, Lara I. Rakocevic, Atanu Giri, Kenichiro Negishi, Sabrina M. Drammis, Ki A. Goosens, Travis M. Moschak, Alexander Friedman

**Affiliations:** Department of Biological Sciences, University of Texas at El Paso, El Paso, TX, USA; Computational Science Program, University of Texas at El Paso, El Paso, TX, USA; National Institute on Drug Abuse, Baltimore, MD, USA; Artificial Intelligence Laboratory, Department of Computer Science, Massachusetts Institute of Technology, Cambridge, MA, USA; Departments of Psychiatry and Pharmacological Sciences; Center for Translational Medicine and Pharmacology; Friedman Brain Institute; Icahn School of Medicine at Mount Sinai, New York, NY, USA

## Abstract

Electrophysiological implants enable exploration of the relationship between neuronal activity and behavior. These technologies evolve rapidly, with multiple iterations of recording systems developed and utilized. Chronic implants must address a litany of complications, including retention of high signal-to-noise ratio in probes and the ability to withstand excess force over the experimental period. To overcome these issues, we designed a chronic implant for rats. Our comprehensive protocol optimizes the entire implant process, from assembling and testing the probes (Neuropixels) to implantation. In addition to addressing the complications previously mentioned, our implant can vertically adjust probes with micron precision and is constructed using modular components, allowing it to be easily modified for various research contexts, electrophysiological recording systems, headstages, and probe types.

## Introduction

*In vivo* electrophysiological recording of neural activity in animal models is a critical tool for exploring and observing the activity of different brain regions during various behaviors^1,2^. The technology that enables these recordings has evolved drastically over time^3,4^. One of the first methods for recording brain activity was using a single microwire electrode in anesthetized animals^3,5^. Further work enabled single electrode neural recordings in awake, head-fixed animals^6^. Recording technology then advanced with the development of new probes (including electrode arrays^7,8^, tetrodes^9,10^, silicon probes, optical probes^11,12^, and others^3^) and associated implants (that were developed to leverage the different probes or increase the number of probes/channels that could be used simultaneously) to record neuronal activity while animals performed behaviors^13–19^. Despite these advances in implant development and probe technologies, there remain several areas where they could be improved; including making precise implantation easier, reducing mechanical displacement of the implant or probes post-implantation, enabling implants to be compatible with different recording systems, and striking a balance between weight and durability. A major division across implants is whether they are designed for acute^1,20,21^ (used for a day or two) or chronic^21,22^ (left implanted for weeks or months) recordings. Chronic implants must account for several additional factors like damage to the probe^16,23^, inflammation from the implantation process^24,25^, damage caused over time as the probe moves within the tissue^26,27^, scar tissue forming around the probes after surgery^26,28^, and the weight of the implant^13,16,29^. These issues have resulted in several variations in implant design and implantation protocols; for example, using more flexible materials when making probes^30,31^, altering the dimensions of the probe^32–34^, lowering probes slowly^35^, or using ultrasonic vibrations^36^ while lowering the probe to minimize the inflammation response and increase the clarity of the recordings.

As implantation technologies and techniques have improved, they also have been modified by prioritizing/adding/altering features of an implant to be better suited for a variety of use cases^37,38^. Some chronic implants have been designed to be very lightweight, thus they have a minimal or nonexistent on the subjects’ behavior though these implants are generally developed for mice with some implants providing modifications that allow for their use in rats^16,22,29^. There are other implants that are designed to be durable, particularly for rats, though these implants are heavier than alternate options^17,39^. Other implants may use a motorized mechanism for moving probes which adds weight to the implant and impedes the number of probes that can be used^15,19^. Certain implants add the capability of inserting probes at different angles or into a wider array for multiple recording sites^17,18^. Each of these implants can be great for different use-cases and trade-offs. We wanted to create an implant that is durable for extended use in rats (even those who have taken a stimulant like cocaine), able to vertically adjust probes with micron-level precision, minimize lateral movement of the probes, is tractable to advances in probe/implant technology, usable with different types of probes, and is compatible with multiple recording systems (we demonstrate the implant with both the OpenEphys ONIX and Neuropixel’s Data Acquisition systems), while maintaining a high signal quality. We developed our implant to be easy to assemble and simple to adopt; using only 3D-printed and commercially available parts. The implant is also comprised of modular components, allowing for easier modification and adaptation of the implant to suit different purposes. Along with the implant itself, we also have developed an accompanying implant kit that includes several tools for implant construction, testing, and a custom screwdriver design that enables high precision adjustment of probes post-implantation.

## Results

### Implant kit overview

We created a comprehensive implant kit designed for chronic *in vivo* electrophysiological recordings in rats. The implant can be constructed and assembled completely “in house” with higher-end consumer-grade 3D printers and widely available parts. The implant is simple to construct and can be built in two days, including 3D-printing, washing, curing, and adhesive curing when creating the individual components (**Fig. 1a**). The entire implant with 2 probes weighs ∼ 8.4 . g. When paired with our custom-designed screwdriver, probes can be vertically adjusted with micron precision (**Fig. 1b**). We also include a method for testing probes early in the assembly process (**Fig. 1c**). The implant was designed to be small and lightweight while being durable enough to withstand chronic use in rats (**Fig. 1d**). The implant can be built on a holding block, protecting the probes and making it easier to assemble (**Fig. 1e**). Our implant is compatible with different headstage platforms (**Fig. 1f**) and not all pieces need to be modified to allow the implant to be compatible with different probes. The shuttle shown is designed for use with Neuropixels 1.0 silicon shanks^40^, with a new design for Neuropixels 2.0^41^ provided as well. Overall, simplicity, adaptability, durability, accessibility, and precision are all important factors implemented in our implant design.

**Fig. 1.**
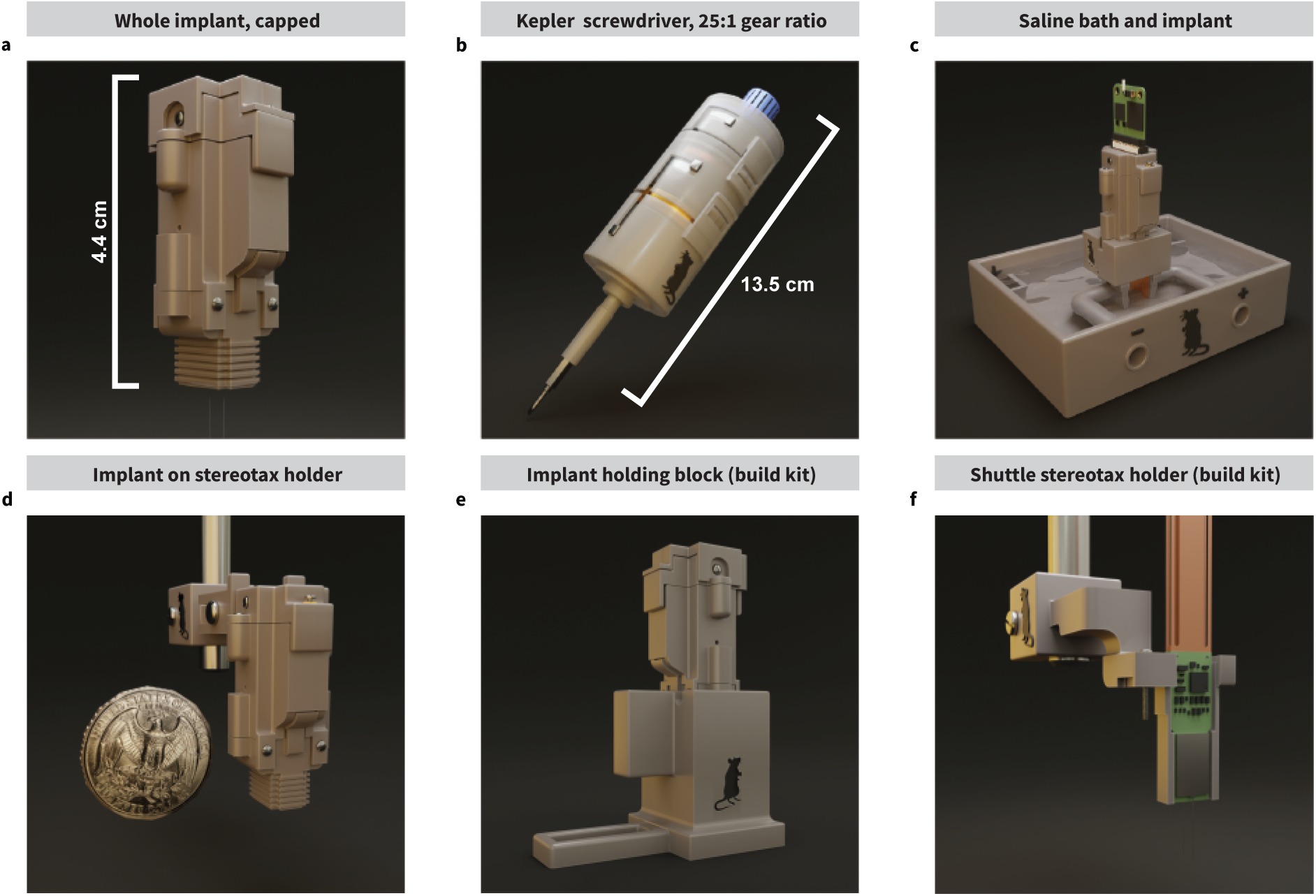
Kit overview. **a.** The implant is 3D-printed “in house” using a mid-high-end commercial 3D printer, is relatively easy to assemble, and features modularity to give the user the most power and control to adapt it to different needs. It also features a driving screw mechanism with a movable shuttle to adjust the vertical position of the probe(s) after surgery to reach a target brain location over several days. **b.** The Kepler screwdriver, a mechanical screwdriver with a gear system which, in conjunction with the implant, enables slow, micron-precision vertical adjustments to the probe’s position to not only keep track of location, but also protect the probe from damage as the probe shank traverses the brain tissue. Each turn at the input of the screwdriver results in a 25th of a turn at the output, which allows for a minimum of 0.012 mm adjustments with a 0.3 mm pitch screw. **c.** A custom testing environment for evaluating gain and noise characteristics of the probe and acquisition system. It consists of a receptacle to hold saline solution, stimulating electrodes, distal receptables to hold ground wires, and a detachable implant block which holds the assembled implant over the saline. **d.** The stereotaxic adapter, made to hold the implant in place during surgery. **e.** A large holding block as a part of the assembly kit to provide protection to the probe(s) while the implant is being assembled. If the probes are not retracted into the implant, the holding block protects the probe shanks in a tall channel to prevent them from being touched. **f.** The shuttle stereotactic adapter as a part of the assembly kit used to lower the fragile probes into the implant with the help of a stereotactic device. The shuttle is held in place with the same (or similar) drive screw used to adjust the probe’s vertical position on the implant.

### Implant components and reasoning behind design choices

To allow for our implant to be maximally flexible and generalizable, it is comprised of five modular components that can be modified without changing the other pieces, allowing the implant to be easily tailored to different research requirements (**Fig. 2a, Supplemental Fig. 1a**). The shuttle component holds the probes within the implant and consequently is the piece that can be altered to change the number, or types, of probes used in the implant (**Fig. 2b, Supplemental Fig. 1b**). Here, we used Neuropixels 1.0 probes but also provided prototype implant designs for Neuropixels 2.0 probes and optogenetic fibers for closed loop optogenetic and electrophysiological recording experiments. The shuttle piece also moves probes vertically via a driving mechanism using a threaded insert. This vertical movement is possible by using a fine screw (space between threads [*pitch*] ≤ 0.5 mm, we used a 0.3 mm thread pitch in our design to prevent “thread skipping”) in the driving mechanism which allows for finely controlled movement along the vertical axis. This fine vertical movement is important for reducing damage and thus maintaining a high signal-to-noise ratio through chronic recording experiments. In addition to the driving mechanism, the shuttle is also held in place by a “rail” molded into the inside of the implant body (**Fig. 2c**) and is protected with a molded grid pattern from the inside (**Supplemental Fig. 1c**). The rail continues into the skull interface (**Fig. 2d, Supplemental Fig. 1d**) to provide stabilization to the shuttle along the entire implant. This prevents the shuttle from moving laterally, which improves stability and reduces sheer forces on the probe shank, thus preventing breakage. The friction of the rail, along with the drive mechanism, prevents unwanted movement along the vertical axis which reduces drift. We designed a headstage interface for our implant that is compatible with multiple recording systems including the Neuropixels Data-Acquisition system^42^ and the Open Ephys ONIX system^43^; made possible by leaving the probes’ Zero Insertion Force (ZIF) connectors exposed at the top of the headstage interface so that any device can be connected through a compatible flat printed circuit connector (**Fig. 2e**). When not in use, the implant can be capped to protect the ribbons (**Fig. 2f**). To protect the probe from any pulling movement at the ZIF connectors and prevent the probes from moving in an unwanted manner, the extra length of the probe’s ribbon is folded into an “S” shape and tucked inside the implant to create some slack. The ZIF connectors are then threaded through two 0.5 mm slots at the bottom of the headstage interface, covering the inside of the implant so that the probes cannot be removed by pulling on the exposed ZIF connectors. The headstage interface piece also has a simple “biting” mechanism which holds the ZIF connectors in place with the two screws mounted onto the piece (**Supplemental Fig. 1e,f**). Using our system, we were able to record two rats for months at a time (one for 112 days and the other for 64 days). If an alternative system is desired, the interface piece can be altered to have an enclosure that is suitable for the recording system being used.

**Fig. 2.**
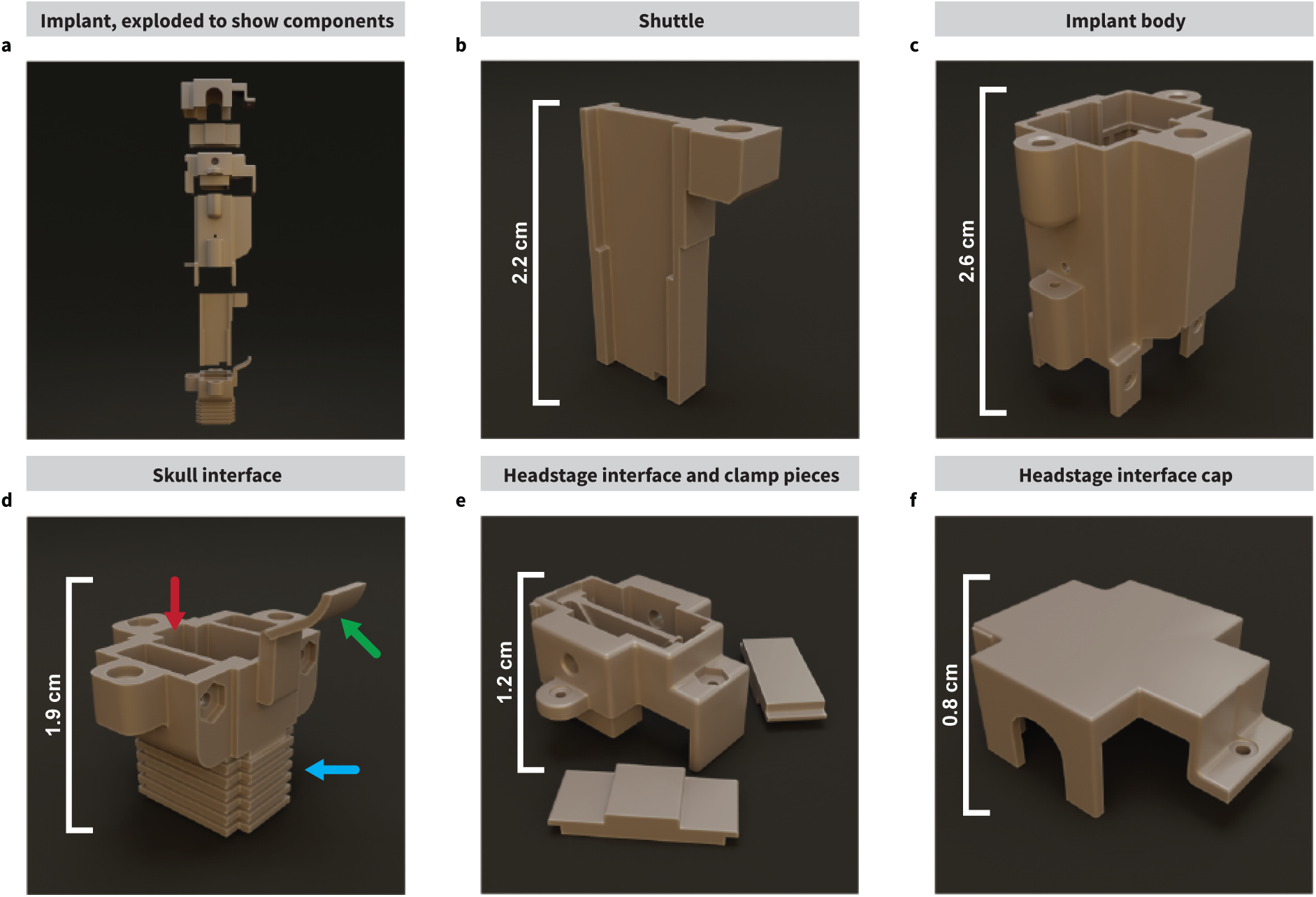
Implant components. **a.** Our modular implant is composed of 5 main parts. **b.** The shuttle can hold up to two probes and is moved on the vertical axis (Z) with a screw mechanism via a threaded insert on the side. **c.** The implant body holds the shuttle in place on an internal rail to prevent it from moving in any axis other than Z. The drive screw is inserted and turned from the top, going down the screw channel on the side. It is also reinforced with a grid pattern from the inside to protect the probes from damage originating from rat behaviors. Ground wires are threaded from the bottom and inserted into the implant body from the sides. **d.** The skull interface also features a rail (**red arrow**) to guide the shuttle vertically. It has two holes on the bottom through which the probe shanks come out. When assembled, the cover piece on the side (**green arrow**) seals the screw channel to protect the drive screw shaft and shuttle from contaminants. The bottom ridges (**blue arrow**) allow cement to adhere to the implant better. **e.** The headstage interface is headstage-agnostic to allow the implant to be used with most acquisition systems’ headstages. The probes’ ZIF connectors come out of the top of the piece and are held in place by a “biting” mechanism which includes two removable “teeth”. The mechanism is adjusted from the sides to hold the ZIFs in place and facilitate headstage connection. **f.** The headstage interface cap simply covers the top of the implant to protect the ZIFs from contaminants. It is held in place by pressure and a small screw.

Chronic recordings necessitate a durable shell to for the implant to protect the probes from animal behaviors. Rats often hit the implant against the walls of their home cages, increasing the likelihood that the probes could be damaged. Therefore, we used SLA 3D-printing instead of FDM. While FDM prints are inexpensive, they offer less strength than an SLA resin print due to structural imperfections. We used Formlabs Tough 2000 photopolymer resin with SLA printing due to its flexibility and durability.

### Custom screwdriver enables vertical movement of probes with micron precision

The number of rotations by the driving screw in the implant’s vertical movement mechanism can be used to precisely control the distance and speed at which the shuttle travels. By making the distance travelled per 360-degree rotation equal to the pitch of the drive screw, sub-dividing the rotations divides the distance proportionally (**see Methods ‘Calculating probe location on the vertical axis’**). For example, if the driving screw’s pitch is 0.3 mm, then a full turn would make the shuttle travel 0.3 mm along the shaft, half a turn would be 0.15 mm, and a quarter turn 0.075 mm. Such granularity slows the shuttle significantly, which prevents excessive stress on the probe shank as it moves through the tissue. This allows the position of the shank within the tissue to be estimated. However, it is difficult to know exactly how far and fast a small screw is rotated by hand. For this reason, we designed the *Kepler screwdriver* (**Fig. 3a**), a precision mechanical screwdriver with concentric cascading planetary gears (**Fig. 3b-e**), each with a gear ratio (GR) of 5:1 for a total of 25:1 meaning it takes 25 full turns at the input of the system to complete 1 rotation at the output (**see Methods ‘The *Kepler screwdriver’***). Thus, the distance travelled by the shuttle would be 0.012 mm per full turn of the *Kepler driver*. The *Kepler screwdriver* can be modified to change screwdriver bit compatibility (**Fig. 3f**), and limits how fast the screw can be turned to help protect the probes and has an optional mechanical counter attachment to limit human error when counting turns (**Fig. 3g**). The counter attachment tracks input rotations with a 10:1 planetary gear. This planetary gear rotates a disk with whole number etchings while fractions of a rotation are displayed at the knob (**Fig. 3h,i**). Importantly, the attachment’s GR is not counted towards the mechanism’s total GR and attaches at the top of the device, highlighting the modular design the implant itself also features. We provide instructions on how to 3D print and assemble the *Kepler screwdriver* in **Supplemental Note 1**.

**Fig. 3.**
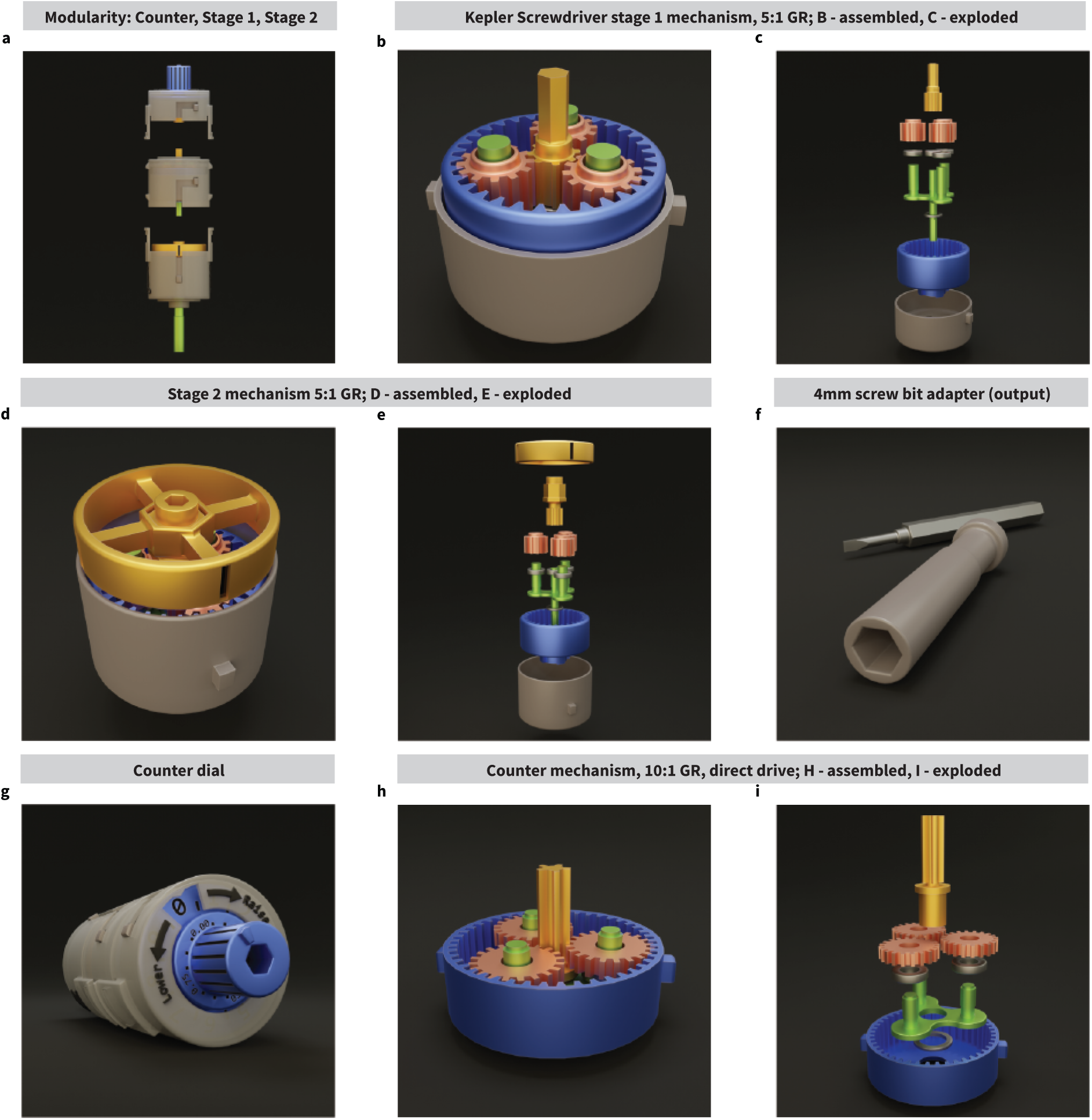
The Kepler screwdriver. **a.** The “Kepler screwdriver” or simply “Kepler driver” is a modular, 2-stage mechanical gear screwdriver which converts one rotation at the input into one 25th of a turn at the output. This is to help turn the implant’s drive screw slowly and precisely. For a drive screw of pitch = 0.3 mm, one turn of the Kepler driver would result in the shuttle component of the implant to move 0.012 mm along the shaft of the screw. **b-e.** The Kepler driver’s first (**b, c**) and second (**d, e**) gear stages are each a planetary gear system with a GR of 5:1. Because the first stage feeds into the second stage, the two stage’s GRs are multiplied together, resulting in a GR of 25:1. Their input is located at the central sun gear (yellow), and the output is on the carrier (green) which rotates within the stationary ring gear (blue), driven by three planetary gears (pink). Each gear mechanism is held in a case (gray) which holds the ring gear in place from the bottom. The main differences between the two stages are the sun gear, which has a hexagonal shaft on stage 1, and a hex socket at stage 2 (**c, e**). Stage 2 has an added ring indicator which simply follows the movement of the sun gear to show movement. **f.** The second stage carrier connects to the screwdriver bit adapter, which is originally made with a 4 mm hex socket for small bits; however, it can be easily re-sized or replaced if needed. **g.** To help prevent errors in turn counting, which is critical to estimate the shuttle’s travel distance along the drive screw, we have also designed a detachable counter module. The module’s gear system does not alter the turn rate of the input, rather it simply passes the turns directly down to the first gear stage of the screwdriver. The counter’s gear system output turns a disk which indicates the number of turns completed, with the counter dial indicating fractions of a turn. **h, i.** Similar to the other gear systems on the screwdriver, the counter features a planetary gear system with a GR of 10:1; thus, a whole rotation of the input is converted into a 10th of a rotation at the output. The input to the system is the sun gear, whereas the output is the carrier. However, unlike the other gear systems, the sun gear passes through the carrier to drive the input rotations directly downward onto the next gear system so that the rest of the mechanism’s turn rate is not affected by the counter module.

### Comprehensive build kit simplifies implant assembly, protects probes, and aids in implantation surgery

Electrode arrays tend to be very expensive, and the micron-thin nature of the probe shank makes them extremely fragile. If a shank is damaged, the entire probe becomes unusable. This is a challenge when loading the probes into the implant and throughout assembly. To minimize the risk of the probes breaking, we developed an assembly kit and protocol to make the process accessible to anyone (**Supplemental Note 2**). The assembly, after printing, washing, curing the 3D printed prices, and letting the adhesive dry can be completed in four hours. We developed an adapter so that a stereotax can be used to load probes onto the shuttle (**Fig. 4a**); this allows for probes to be maneuvered safely and lowered into place with the optional aid of a surgical microscope or a magnifying glass to monitor the shanks (**Supplemental Fig. 2a-c**). The shuttle and probes are then lowered into the skull interface which is held in place by our custom-designed holding block (**Fig. 4b,c**), thus protecting the probe shanks. The implant assembly can be completed with the two implant pieces on the block, keeping the shanks safe. When finished, the probes can be retracted into the implant, shielding them and preventing accidents. The implant can remain on the holding block until it is implanted. For surgery, we developed a stereotaxic adapter to hold the completed implant (**Fig. 4d,e, Supplemental Fig 2d**) which keeps the implant secure during implantation.

**Fig. 4.**
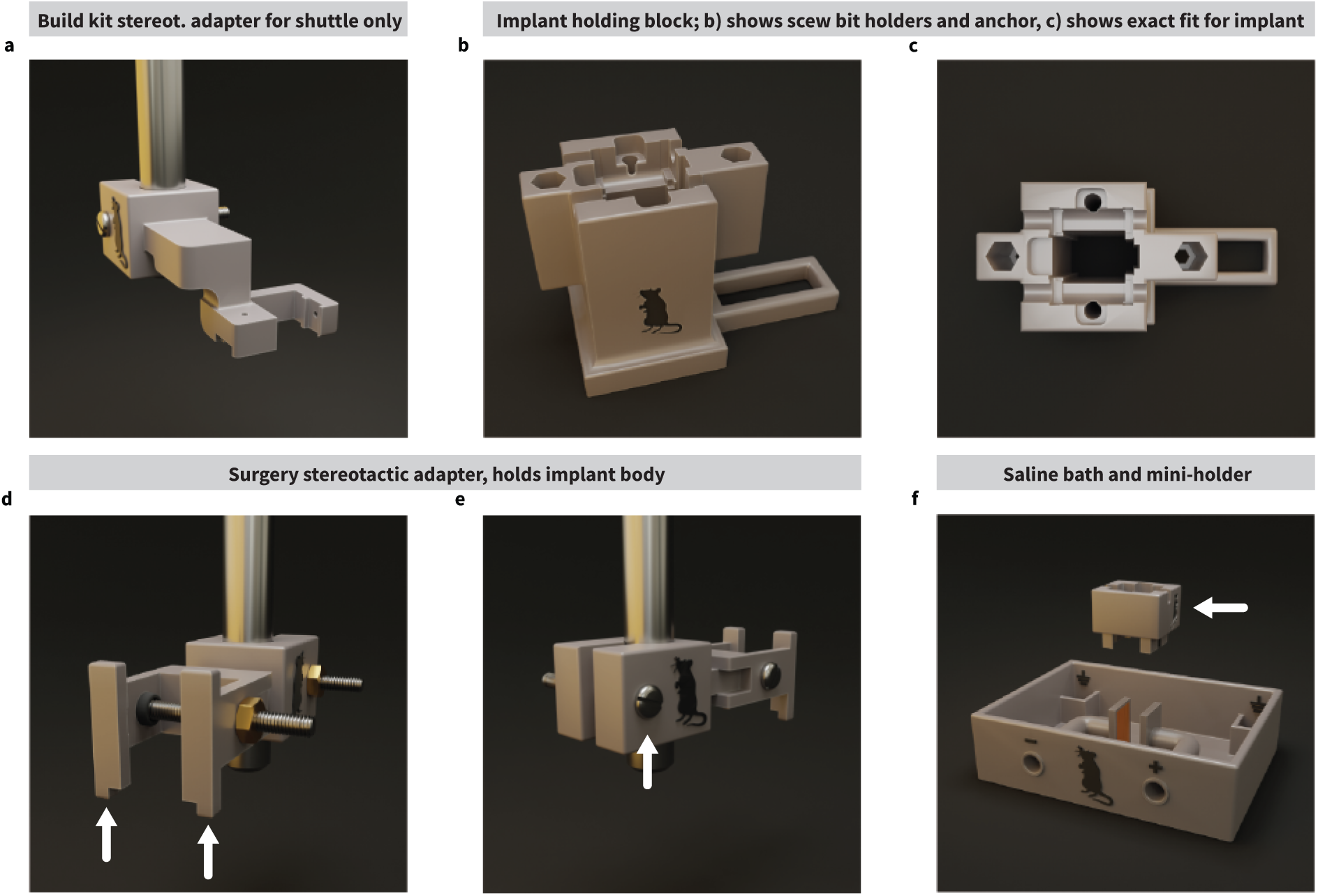
Assembly, surgery, and testing accessories. **a.** The shuttle stereotactic adapter is designed to hold one implant shuttle piece loaded with one or two probes. Its purpose is to leverage the three-coordinate system of a stereotaxic device to maneuver the shuttle and probes into position and slowly mount them onto the implant safely. The adapter block which attaches to a standard stereotaxic holder has a screw and nut to tighten the adapter around the holder. **b, c.** The implant holding block is designed to tightly hold the skull connector piece with the purpose of aiding in implant assembly. The holding block has an empty channel down the center tall enough to protect the probes’ shanks from damage when they have not been fully retracted. It also features two hex sockets which can hold a 4 mm screwdriver bit on each side, for storing alongside the implant in preparation for surgery. The tail also serves as an additional anchor point to screw the block in place. **d, e.** To hold a fully assembled implant steady during surgery, we created the implant stereotactic adapter. The adapter ‘arms’ (**d, arrows**) slide onto the implant and hold it by the headstage interface piece. The screw and nut can be tightened to ensure that the implant is held firmly in place. Similarly, the screw and nut on the adapter (**e, arrow**) can also be tightened to ensure that the adapter does not slide off the stereotactic device. **f.** To test probes before committing to implanting them, we also designed a saline bath. It consists of a receptacle to hold saline, two smaller receptacles for the probes’ ground wires/electrodes, and two electrode plates that flank the probe shank while it is in the saline to apply a voltage across them for noise and gain testing. The mini holding block (**arrow**) holds a fully assembled implant and is fully detachable from the setup. This allows the saline bath to be used with or without the implant.

Another feature of our implant kit is a tool for testing the probes before implantation. This can give useful insight into the electrical characteristics of the probes and the system they are connecting to while also ensuring the probes work after implant assembly. For Neuropixels probes it is recommended to test for gain and noise by dipping the shank into saline and initiating a recording as a trouble-shooting step alongside software self-tests. To make testing easy after the implant is completed, we designed a 3D-printed saline bath which features a tray that holds saline, two copper plates flanking the electrode shank, and small receptacles distal to the probes for the ground (**Fig. 4f**). To hold the implant above the saline, a miniaturized holding block was built which also allows for the adaptation of the bath to different implant shapes or to test the probes by themselves on a stereotaxic device **(Supplemental Fig. 2e,f)**. This saline bath allows for noise and gain testing on the probes before implantation (**see Methods ‘Probe testing’**).

### Implantation surgery

During implantation surgery, the implant is mounted onto a holding piece that can be connected to a stereotax to simplify aligning and lowering the implant to the desired brain target (**Fig. 5a,b, see Methods ‘Surgical and post-surgical procedures’**). To maximize the lifespan of implanted probes, several measures were taken. To start, the implant was affixed to the rat’s head in the most stable configuration possible to prevent damage from rat behaviors. To reduce this damage, we cleaned the skull surface of connective tissue, scored the skull, and added eight anchor screws (**Supplemental Fig. 3a-e**) around the implant, which were then covered in dental cement so that the implant had a strong foundation. These screws, in combination with a dental cement foundation, keep the implant in place during behavioral manipulations (**Fig. 5c,d**). After the probes are set within the tissue, the application of Vaseline around the base of the implant (**Supplemental Fig. 3h**) can prevent discharge and blood (which can potentially obstruct movement along the shuttle rails) from making their way into the body of the implant. At the end of the surgery, we sutured the skin at the anterior and posterior skin-implant interfaces and cauterized the remaining tissue around the implant with silver nitrate to prevent tissue from regenerating under the dental cement (**Fig. 5e, Supplemental Fig. 3i,j**). Post-surgical care is also important to prevent implant failure. To prevent infections, animals were given routine broad-spectrum antibiotic injections (Enrofloxacin). To reduce itching and to locally disinfect the surgical site, we routinely applied Hyaluronic acid and Chlorhexidine on the live tissue around the implantation site (**Supplemental Fig. 3k**). A combination of nystatin, neomycin, thiostrepton, and triamcinolone can also be used to treat the area. Finally, modifications to the animal’s home cage can also prevent damage to the implant; for example, a taller home cage with no point for the animal to pull on the implant can prevent the rat from tearing the implant off (**Fig. 5f**). We provide a full surgical protocol with information on the modified home cages in **Supplemental Note 3**.

**Fig. 5.**
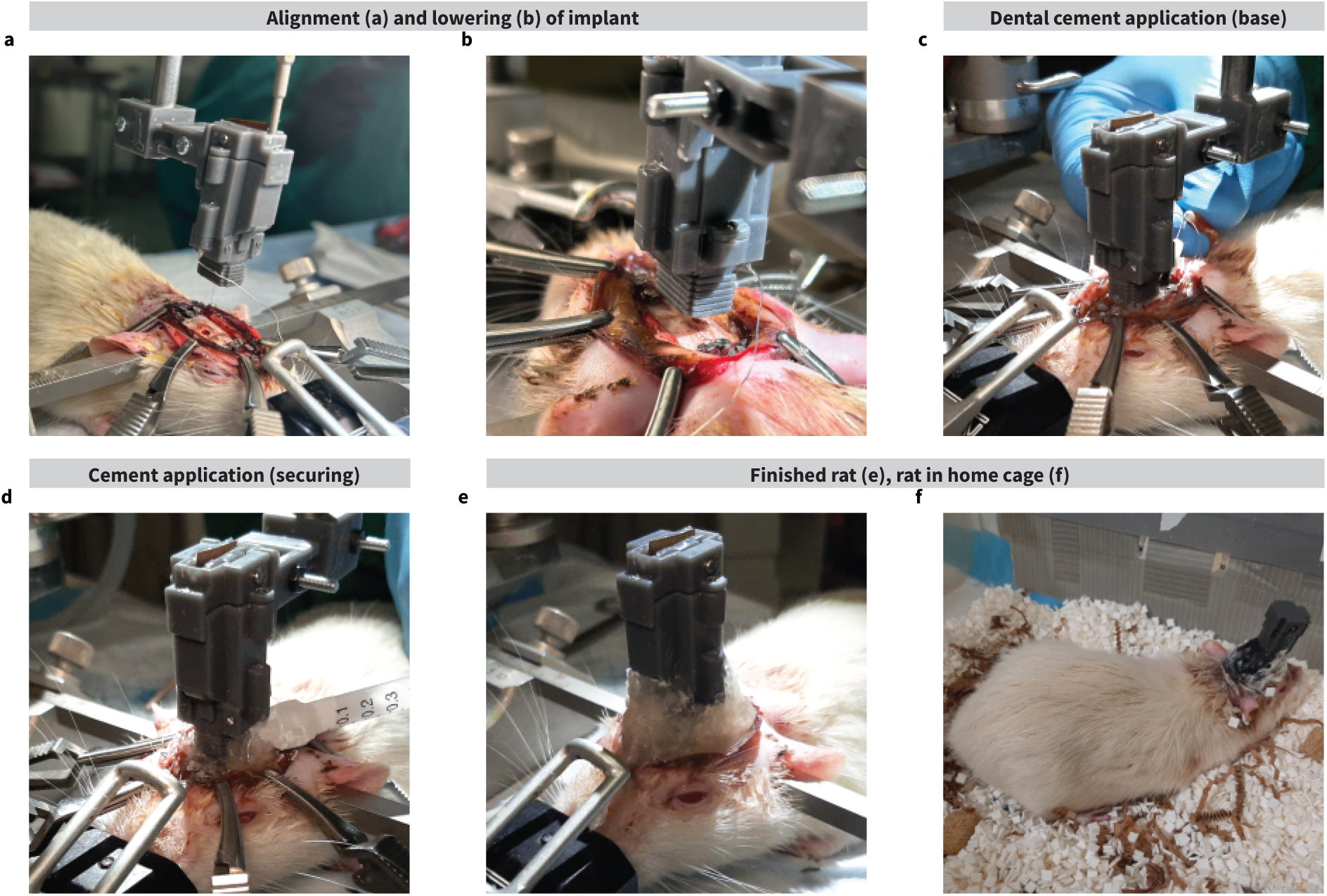
Surgical procedures. **a.** Implant alignment on the medial-lateral (ML) and anterior-posterior (AP) axes to implant the probes near the dorso-medial striatum. During this process, the probe is drawn just enough so that it is implanted at a known depth, taking initial target depth, skull thickness, and implant wall thickness into account. The amount of drive screw turns to reach this length is calculated using Equation 1 (see ‘Adjustments as a function of drive screw turns’). **b.** With the probe drawn at the appropriate length, the implant is lowered onto its final position on the dorsal-ventral axis (DV). Care is taken to do so slowly, watching the probe shanks for buckling with the help of a surgical microscope. The tip of the shank is ‘bounced’ on the brain tissue a few times to test for buckling; if buckling is observed, the implant is raised, and the tissue is re-hydrated with saline and dexamethasone. **c.** After the implant has been lowered onto the skull and the probe has been implanted, sterile Vaseline is applied to the bottom on the implant to prevent blood from entering the implant during the healing process. Metabond is also applied to the bone, screws, and base of the implant to create a foundation for the implant and strengthen adhesion of dental cement, as well as insulate the ground screws from the surrounding tissue. **d.** The dental cement is applied to the Metabond on the base of the implant and built up to cover the skull connector. This ensures a strong connection between the implant, cement, and bone. **e.** After the dental cement has cured, the remaining tissue around the implantation area is cauterized using silver nitrate. Finally, a post-surgical adjustment of the probe is done to slowly lower the probe further into the tissue using the Kepler driver (0.12 mm, or 10 turns with a screw pitch of 0.3 mm). The amount of turns to reach this distance is calculated using Equation 2 (**see ‘Adjustments as a function of drive screw turns’**). **f.** The rat is removed from the stereotactic device and post-surgical care is initiated. We applied a topical antibacterial antifungal corticosteroid cream and injected Enrofloxacin (antibiotic) and Meloxicam (NSAID) in a 10 ml Ringer solution immediately after (as well as three days after) surgery to reduce the chance of complications during recovery (**see Supplemental Note 3**). The rat is placed in a modified home cage for recovery. Figure depicts a rat weighing ∼ 500 g.

### Probe alignment, lowering to initial location, and vertical probe adjustment post-surgery

Lowering probes into brain tissue is a delicate process. One method for preventing probes from breaking is to lower them slowly while monitoring the shanks for buckling with a surgical microscope. We do this by lowering the implant slowly, raising it back up if needed, until we are sure that the shanks are inserted with minimal force. If the shanks buckle on the tissue during this procedure, it may be due to the tissue drying/hardening, which can be rectified by hydrating the tissue with 1 to 2 drops of 4 µg/mL dexamethasone. This complements the implant design where the entirety of the probe, shanks included, can be safely withdrawn inside the implant. During surgery, the probe shanks are only drawn after the surgical site is ready for the implant to be lowered onto the skull. The probes are at risk of breaking both immediately after surgery due to inflammation of the tissue, and at later time points because of scar tissue formation around the electrode shank; inflammation and scar tissue can also reduce recording quality over time. To minimize the dangers of moving a fragile object through scar tissue we leveraged the adjustable nature of the movable shuttle by making micro-adjustments using our *Kepler screwdriver* postsurgery (**Supplemental Fig. 3l**). With this approach, we slightly adjusted the position of the probe shank within the tissue in a slow and controlled manner to avoid breaking the probe shank and tracking the depth of the shank tip after each adjustment. To keep track of where the tips of the probe shanks are, we first must implant the probes at a known depth. To do this, we first measure the thickness of the skull with calipers, then draw the probes out of the implant to that distance plus the thickness of the skull interface’s bottom wall (0.5 mm). We next carefully lower the implant into place with the help of a stereotax while monitoring the probe shanks for buckling using a surgical microscope. Once the probes are implanted, they are further inserted into the brain (100 – 300 µm) post-surgery, and with our precise micro-adjustment protocol, we can track what depth the probes are located at after each adjustment. Further details about this process can be found in **Methods ‘Calculating probe location on the vertical axis’ and ‘Keeping track of probe depth during and after surgery’**. Using this method, we reached the dorsal striatum (-3.45 mm DV) within 3 weeks when making 100-300 µm adjustments 2-3 times per week. After the target was reached, only 50 µm adjustments were made every other week or when the signal degraded where spikes were undiscernible from local field potentials to maintain a high signal-to-noise ratio for at least three months after surgery (**see Methods ‘Post-surgical micro-adjustments’**).

### Implant grounding considerations

To ensure high quality signals are recorded, certain considerations must be accounted for during the implantation surgery to minimize the presence of artifacts. In electronic systems, a ground is the reference point from where voltage is measured. If the ground circuit is conducting a noise signal, that noise will be added onto the recorded signal and amplified by the recording equipment; this can overpower lower amplitude (roughly 10 – 100 µV) extracellular recordings or saturate the signal processing hardware, impeding signal analysis techniques such as spike sorting. In our surgical protocol we address this issue by grounding our probes as far away as possible from electrically active tissue and avoiding interconnecting probes to prevent ground loops. Because muscle tissue is electrically active (producing signals in the 1-100 mV range), it is important that any muscle artifacts are avoided by clearing the implantation area of muscle connective tissue during surgery. Proper grounding is critically important to avoid ground loops since they may introduce 50-60 Hz of electromagnetic interference artifacts; we provide more information in **‘Grounding considerations’**.

### Electrophysiology recordings from the implant

With our implant we successfully recorded two rats using the Neuropixels Data Acquisition System and the Open Ephys GUI. One was recorded over 64 days and the other over 112 days (**see Methods ‘Recording and Signal Processing’**). Using our implant, we were able to achieve clear recordings with a high signal-to-noise ratio (**Fig. 6a**). We would maintain the same probe depth if the recording quality was maintained, typically for five days, and then would lower the probes 60 µm which would once again give us high signal-to-noise ratio recordings at the tip. We also noticed minimal lateral movement of the shuttle after assembly prior to surgery, or drift during our electrophysiological recordings. From these recordings, we were able to extract various spike waveforms in MATLAB (**Fig. 6b,c, Supplemental Fig. 4a,b**) and spatially relate them to one another across a varying number of channels (**Fig. 6d, Supplemental Fig. 4c**), enabling further spike sorting.

**Fig. 6.**
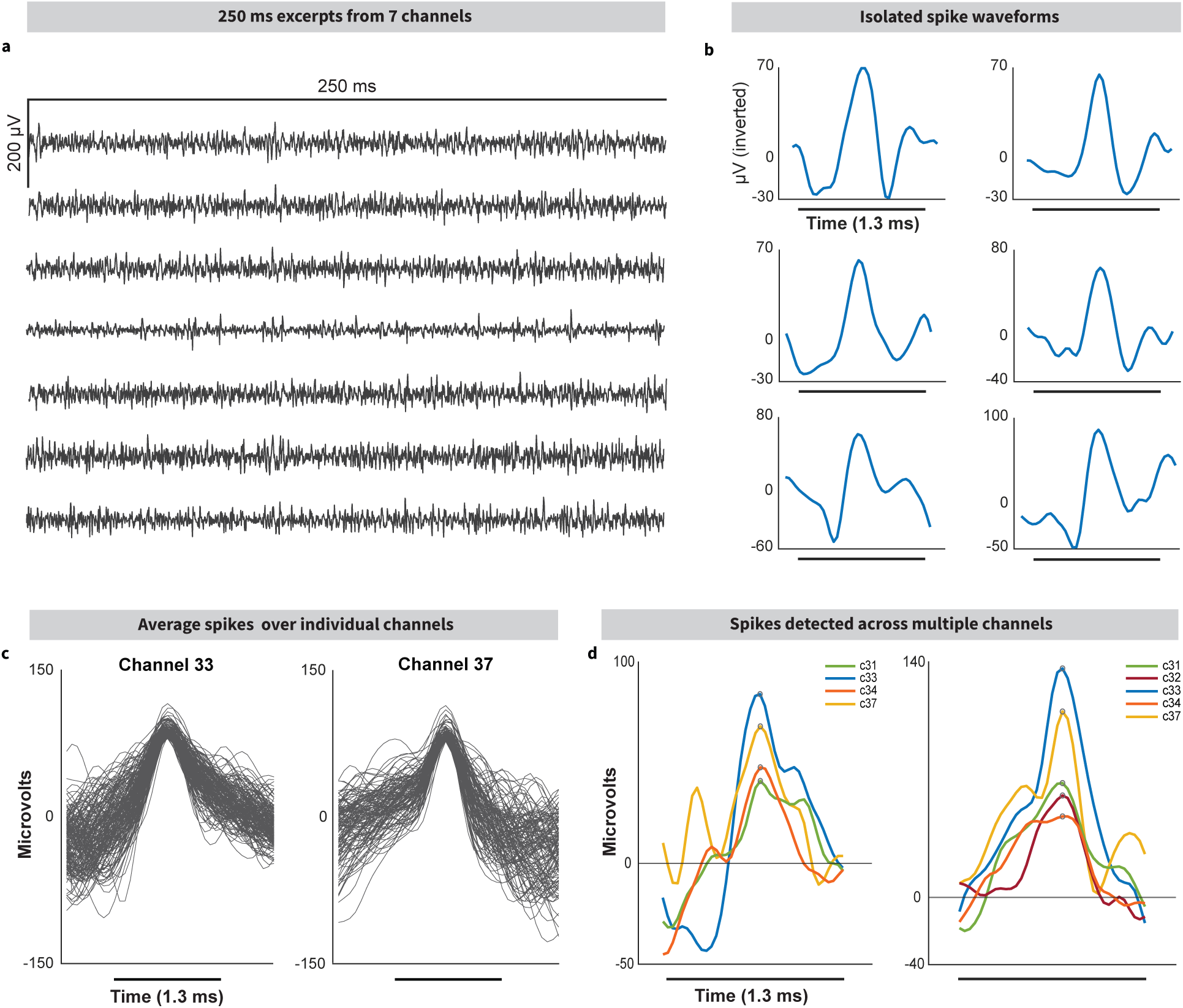
Stable recordings 75 days after implantation. **a.** Continuous recording of extracellular activity from a single Neuropixels 1.0 probe implanted at coordinates –1.2 mm AP, +2.5 mm ML (from Bregma) on a rat. The recording was made the day after the probe was driven -60 µm DV during a post-surgical probe adjustment 75 days after implantation, leaving the tip of the probe situated at -2.74 mm DV within the brain tissue. **b.** Isolated spike examples from (**a**). **c.** Average spike waveform over select channels. **d.** Spike waveforms present across neighboring channels. The general spike shape is conserved across a few physically proximal electrodes, but the spike amplitude is reduced the further from the ‘main’ electrode the spike is recorded. Spikes were aligned to the highest-value sample (circled) and overlaid to show that the general waveform shape is conserved across at least three channels corresponding to neighboring electrodes. This process can be expanded to enable spike sorting.

### Proof-of-concept modifications to the implant

To demonstrate the modifiable nature of our implant we have designed proof-of-concepts that may address common alternative use cases for our implant (**see Methods ‘Implant modifiability’**). Firstly, we designed a prototype for using Neuropixels 2.0 probes instead of the Neuropixels 1.0 probes (**Fig. 7a,b, Supplemental Fig. 5a-d**). This can be achieved by changing the shuttle piece to accommodate the new probe shape (**Supp Fig. 5a**). Secondly, we demonstrate how one may also add pin headers and connectors to the implant body, and electrodes to the skull interface to add electrical stimulation functionality (**Fig. 7c-e, Supplemental Fig. 5e**). Finally, changing the skull and headstage interface pieces to fit optical fiber cannulas can allow for the addition of optical fibers, which would enable closed loop optogenetic manipulation along with electrophysiological recording (**Fig. 7f, Supplemental Fig. 5f,g**). In our prototype, we include two optical fibers situated 500 µm above the electrode probe tip.

**Fig. 7.**
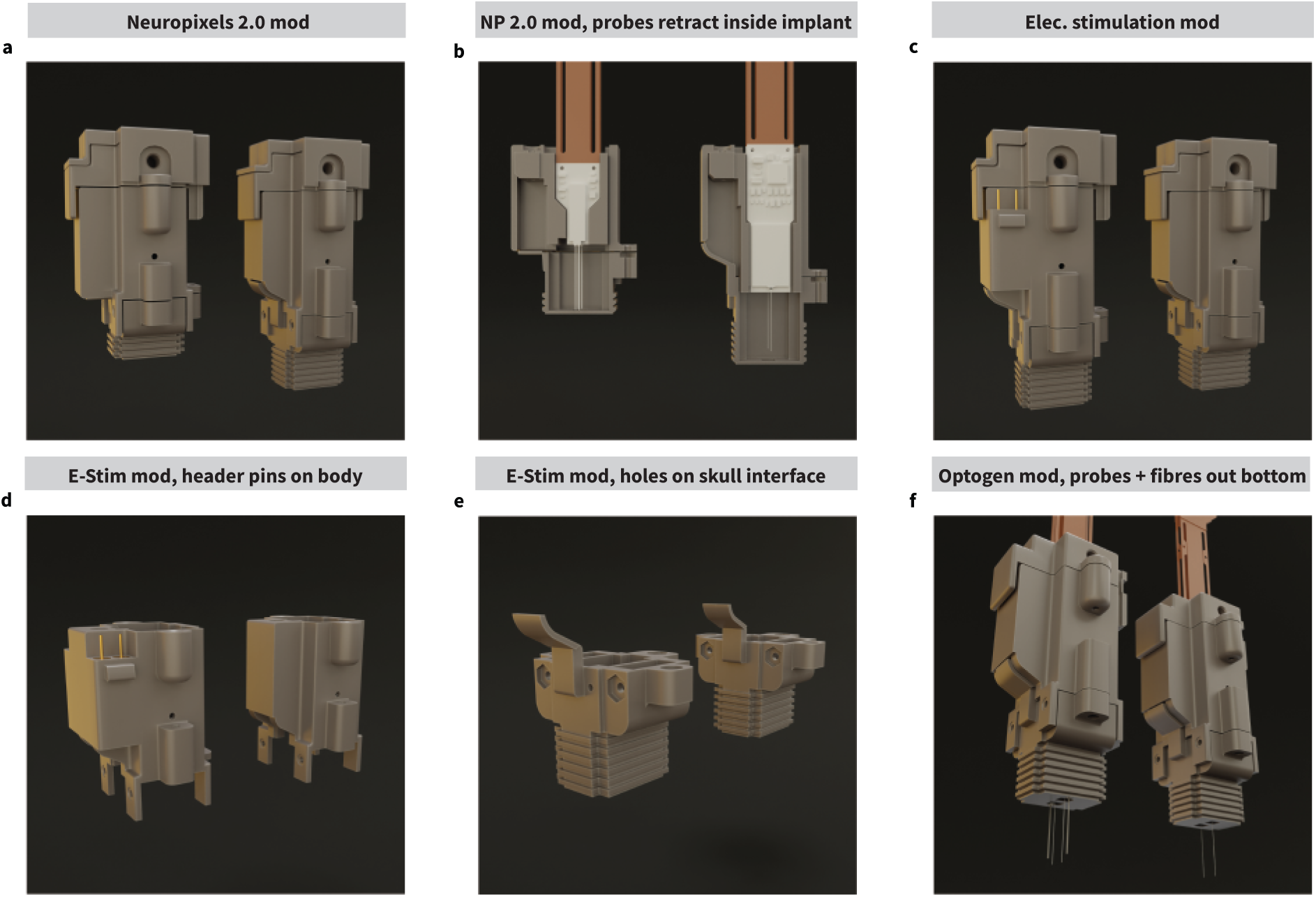
Modularity facilitates modifications. **a.** Side-by-side comparison between an adaptation of the implant for Neuropixels 2.0 (NP2.0) probes (left) and the original (right). In this modification, only necessary implant parts such as the shuttle (not pictured), implant body, and skull interface have been modified, leaving the headstage interface and cap untouched, highlighting the advantage of a modular design. The modifications lead to an 8 mm reduction in height and were done over the course of two days. **b.** A bisected view of both the NP2.0 (left) and original (right) implants showcasing the ability to retract the probes in both cases, offering the same protection to the probes. **c.** Side-by-side comparison between the electrical stimulation modification (left) and the original implant (right). This modification adds the ability to mount stimulation electrodes alongside the probes to expand the utility of the implant. **d-e.** Only the implant body (**d**) and skull interface (**e**) were modified for the electrical stimulation implant. A two-pin header receptacle was added to the implant body on either side of the drive screw rail. A wire would then be soldered to each header pin, one for the cathode (-) and one for the anode (+) of the stimulation electrodes. These wires would then travel down into the skull interface piece. Two holes on the side of the skull interface allow the wire to be fed into the skull interface, and the wires would be soldered onto stimulation electrodes situated at the bottom of the piece. **f.** The final proof-of-concept modification to the implant, the optogenetic modification. In this version static optical fibers are mounted alongside the probes, offset 1.25 mm to the side, and 0.5 mm above the tip of the probe. To make this modification, a fiber cannula receptacle was created on either side of the skull interface from the inside. These receptacles hold one cannula each, through which an optical fiber is threaded. Similarly, on the top of the implant, the headstage interface also holds a cannula on each side to allow optogenetic stimulation devices to be connected.

### Limitations and Potential Adaptations of Interest for Different Research Conditions

While constructing the implant, we did not account for probe removal and recovery following implant use because implants remain in use for months at a time and the multiple connection cycles increase the chance of damaging the Neuropixel’s probes’ ZIF connectors. Despite this, it is possible to recover the probes, though it requires gently cracking open the implant using a Dremel because the screws which hold the skull interface and implant body together often get covered by dental cement during surgery. Redesigning the implant by moving these screws elsewhere or further up the implant could allow for it to be opened with a screwdriver for easier probe recovery. Moreover, the design choice to not house the headstage(s) within the implant does put the fragile headstage electronics and ZIF connectors at risk of damage, however a headstage-agnostic mating method increases the implant’s cross-compatibility with different recording systems, for example the Neuropixels Data Acquistion system and the Open Ephys ONIX system which both feature headstages with different form factors. This targets a specific niche which aims to increase the implant’s usefulness no matter what recording equipment a neuroscience lab has access to. In the case where headstage protection is indispensable, our implant’s modularity enables the experimenter to design and replace the headstage interface piece with a housing piece.

The implant in its current iteration is heavier than some other chronic implant options^22,44^ and while the weight could be lowered by experimenting with different materials or using the implant with smaller probes, for rats, durability would remain a key factor to consider when creating a lighter version of the implant. If a smaller probe is used, then the overall size of the implant can be reduced. The overall implant weight and size of the bottom of the skull interface (measuring 8.3 x 11.6 mm) limits the utility of the implant to studies using adult rats, though this can be addressed by using smaller probes and modification of the implant. Although the implant in its current iteration was not intended for, nor was it tested on mice, we recognize that its weight (about 8.4 gr with one Neuropixels 1.0 probe) is a limiting factor to its use in mouse studies. However, an implant made for mice requires less strict durability requirements than rat implants do, thus implant’s wall thickness (0.5 mm) or build material could be revised for such an adaptation.

Modification to the shuttle piece, rail, and skull interface would also be necessary to target more lateral structures of the brain (larger than the 1.5 mm ML the implant is originally designed for), which may limit its use further. For targeting more medial structures, we believe that only the shuttle piece would need to be modified to reduce probe separation down to 2 mm (1 mm from the zero medial coordinate) by reducing the middle wall of the shuttle to 0.5 mm. The depth of implantation is limited by the length of the probe shank (10 mm) minus 0.5 mm to account for the thickness of the bottom wall of the skull connector, however this is usually sufficient for both rat and mouse brains. Finally, the current implant design can realistically only fit two probes (one on each side); adding more probes may require modifications to the overall implant size which would be dependent on the size and weight of the probes used.

## Discussion

We describe how to assemble, test, and surgically implant our chronic implant for electrophysiological recordings in rats. Apart from the probes, the implant, as well as everything to test and use the implant, can be assembled “in house” with consumer-grade 3D-printers and commercially available parts. The implant is constructed with modular parts, allowing each piece to be modified for various research use cases, including changing the shuttle to accommodate different probes or changing the headstage connector to be compatible with different recording platforms. Alongside the implant itself, we developed accessories that hold the implant on a stereotaxic device when adding the probes and for the completed implant during the implantation surgery. We also created a holding block so the implant could be constructed with minimal human contact with the probes, a saline bath for testing the probes and implant, and a custom-designed high gear-ratio screwdriver that enables probes to be adjusted vertically with micron precision. We also created implant-specific surgical protocols so every step of the process, from making the components to recording, is available within a single implant system.

Due to its modular design, future features can be easily added since each part can be modified separately. One example could be altering the implant-skull interface component to be individualized for each subject. In monkeys, this has been done by scanning the skull before printing the interfacing component^45,46^ and alternatively, it may be possible by using flexible materials that can adjust to different skull shapes. With a better seal, the headstage would be more stable and less cement would be needed, reducing the weight of the implant. Another idea used in non-human primate models is to allow the shuttle to move to different recording locations within the implant^47,48^, allowing researchers to look at a wider region of interest. The weight and durability of the implant could also be tailored to different use cases. For example, a miniaturized, less durable version of the implant could be used in mouse models or, in rats, a vertically shorter probe like the Neuropixels 2.0 may allow for a shorter implant. We designed this implant to be more durable at the cost of being heavier because we were using the implant with rats that were self-administering oxycodone and thus were more likely to damage the implant with excessive force. Another idea to explore could be adding more optogenetic fibers to the implant or finding a way to use the implant for cell-specific activity modulation. The implant could also be modified to add more probes or optical fibers with the upper constraint depending on the weight and size of the implant. One other interesting consideration is to modify the headstage to be more easily connected to the recording system. These future directions and developments aim to improve the utility of the implant and ultimately allow this implant platform to be suitable for various research contexts.

Overall, the principles underlying the entirety of the implant kit, including easy assembly, modularity, highly precise vertical adjustment of probes, adaptability to different recording systems, prioritization of accessibility, and the provision of a comprehensive pipeline for building, testing, and implantation are design considerations that could be useful when development future implants as electrophysiological recording technologies advance. The implant was also designed to have wide appeal for researchers who use rat models to explore neurological activity during behavioral task performance.

## Supporting information

Supplemental Methods; Supplemental Notes 1 through 4

---

**Supplemental Fig. 1.**
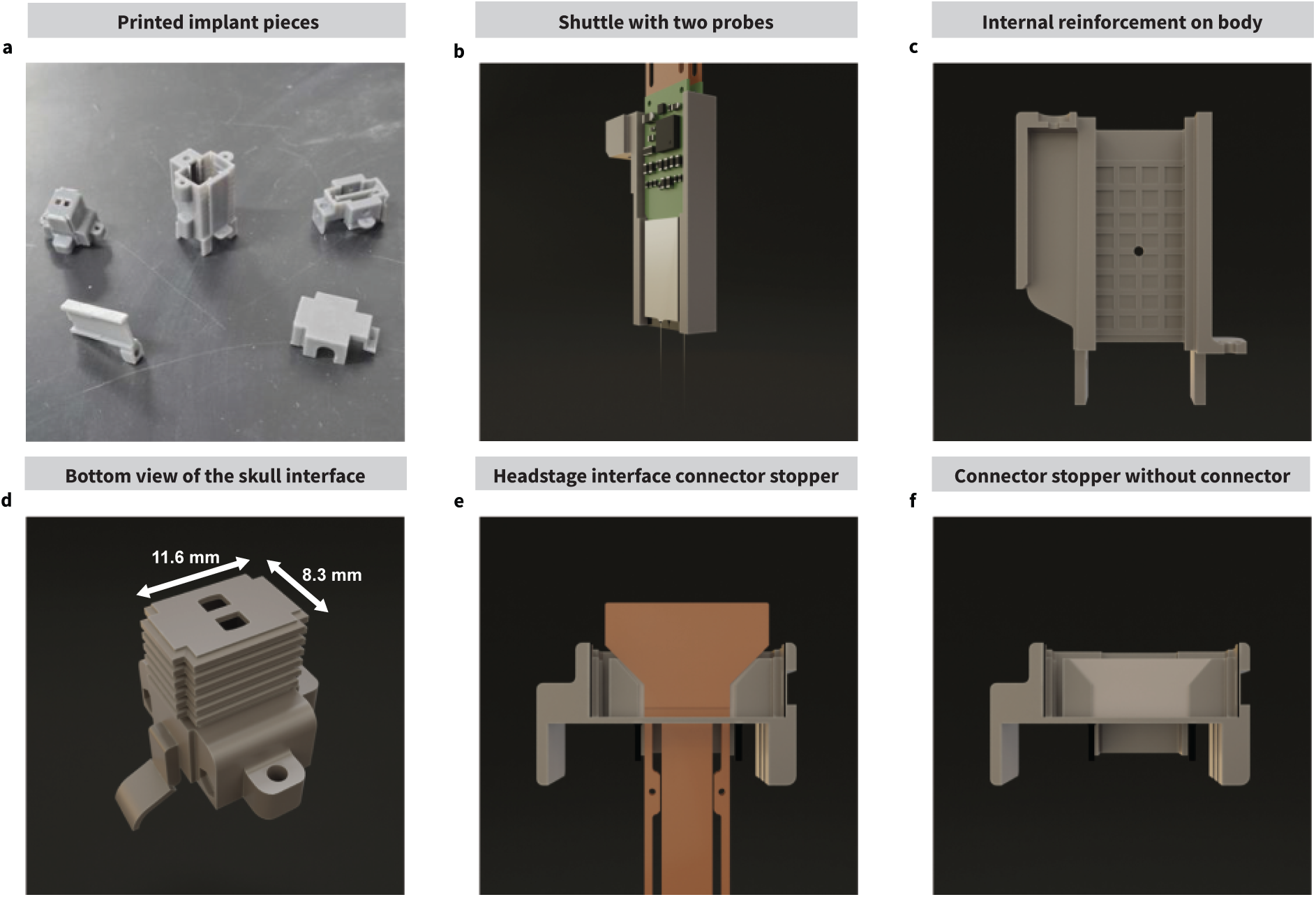
Implant components and design choices. **a.** Photograph showing the individual 3D printed components of the implant. **b.** The shuttle is designed with two “probe beds” on either side. Probes sit on the shuttle back-to-back and 3 mm apart. **c.** Internal reinforcement of the implant body. The “waffle pattern” prevents the walls of the implant body from being pushed inward or breaking. **d.** Bottom side of the skull interface showcasing the two holes where the probe shanks protrude out of the implant. The ribbed pattern on the skull interface helps increase the surface area which contacts the dental cement to strengthen the bond between it and the implant. **e, f.** Bisected view of the headstage interface to showcase the ribbon stopper. This helps keep the probe’s ribbon connector in place to facilitate connection of the headstage(s).

**Supplemental. Fig. 2.**
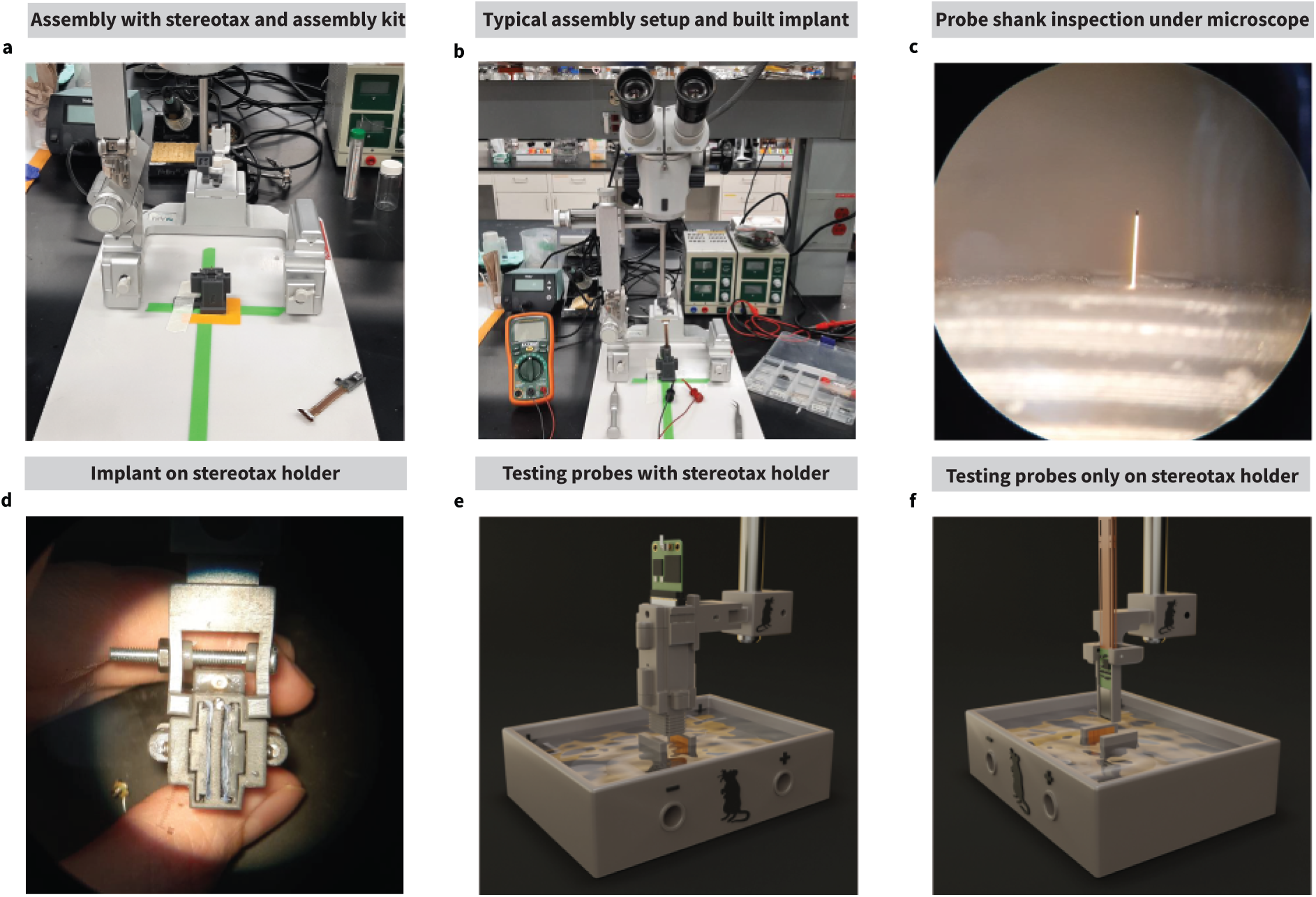
Assembly and testing setup. **a.** Use of a stereotaxic device, shuttle holder adapter, and holder block in the assembly process. The shuttle adapter is placed on the stereotaxic device and the shuttle is mounted onto it. The skull interface piece is placed on the holder block underneath, and the shuttle is aligned and lowered into the skull interface. **b.** Typical setup used to assemble the implant. A surgical microscope is used to monitor the probe shank(s) when lowering the shuttle into the skull interface piece. **c.** Inspecting the probe shank under the microscope after assembling the implant. **d.** Mounting the implant onto the stereotaxic holder implant adapter. **e, f.** Alternative probe testing setups. **e.** Instead of using the mini holder block, the implant is mounted onto the implant adapter and lowered into saline in the saline bath. **f.** The probes are tested on the shuttle without the implant using the stereotaxic holder shuttle adapter.

**Supplemental. Fig. 3.**
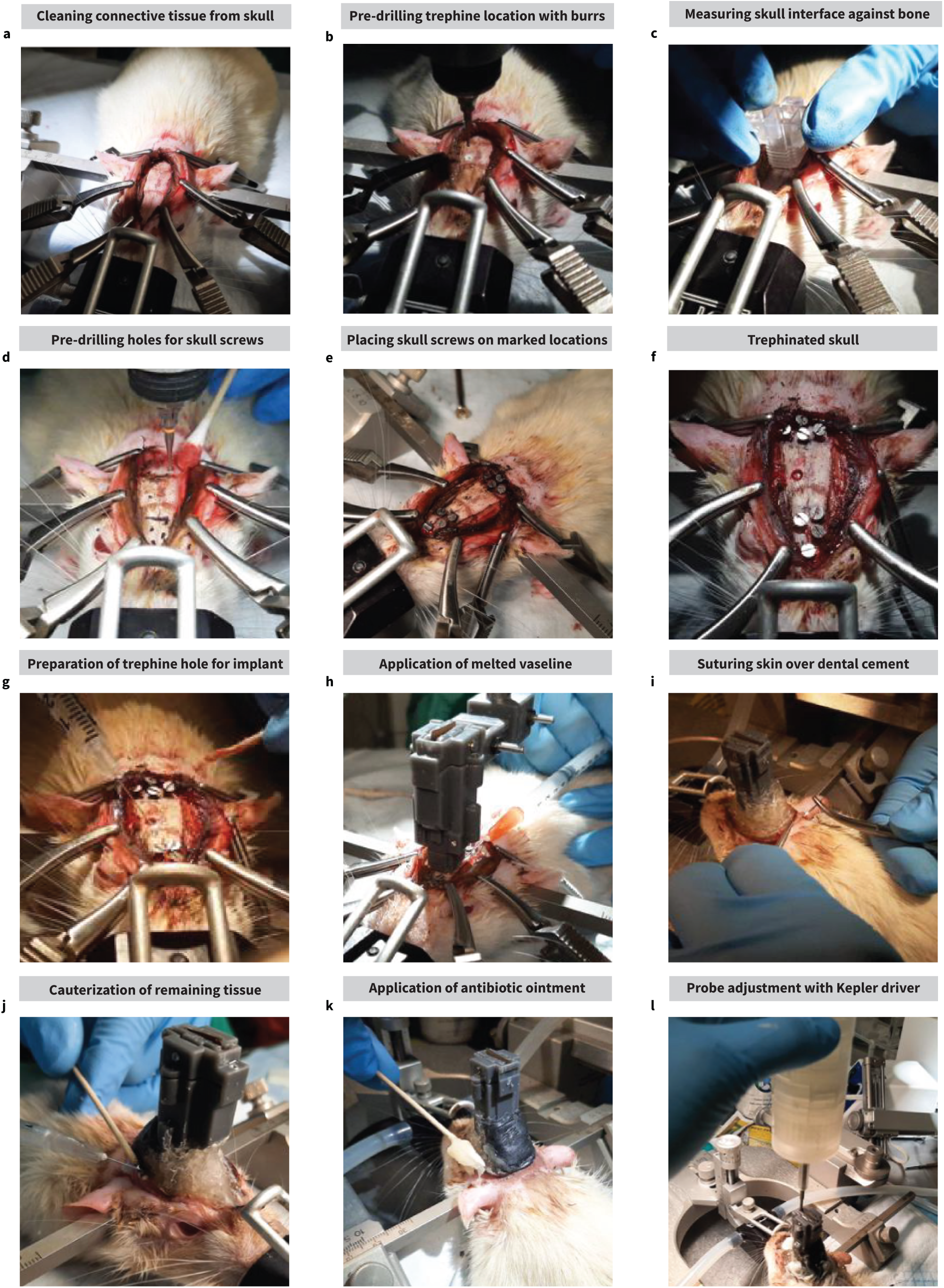
Surgical procedures (cont.). **a.** Clearing the implantation site of connective tissue. **b.** Marking the trephination site with a burrs drill bit. **c.** Measuring and marking the edges of the skull interface piece to plan for anchor and ground skull screws. **d.** Pre-drilling screw sites with a burrs drill. **e.** Final placement of all skull screws. **f.** Trephined skull. **g.** Cleaning and hydrating the trephine hole with 4 µg/mL of Dexamethasone. **h.** Application of melted vaseline to the edges of the implant after lowering it and inserting the probe shanks into the tissue. This helps prevent blood and discharge from seeping into the implant and prevents the shuttle from moving. **i.** Suturing skin over the dental cement to prevent it from scarring underneath and lifting the implant over time. **j.** Cauterizing the remaining tissue around the implant to prevent it from scarring under the implant. **k.** Applying antifungal, corticosteroid, and antiseptic ointment on the exposed tissue to aid the healing process. **l.** Post-op probe adjustment. A large 0.15 - 0.3 mm adjustment is made immediately after surgery.

**Supplemental. Fig. 4.**
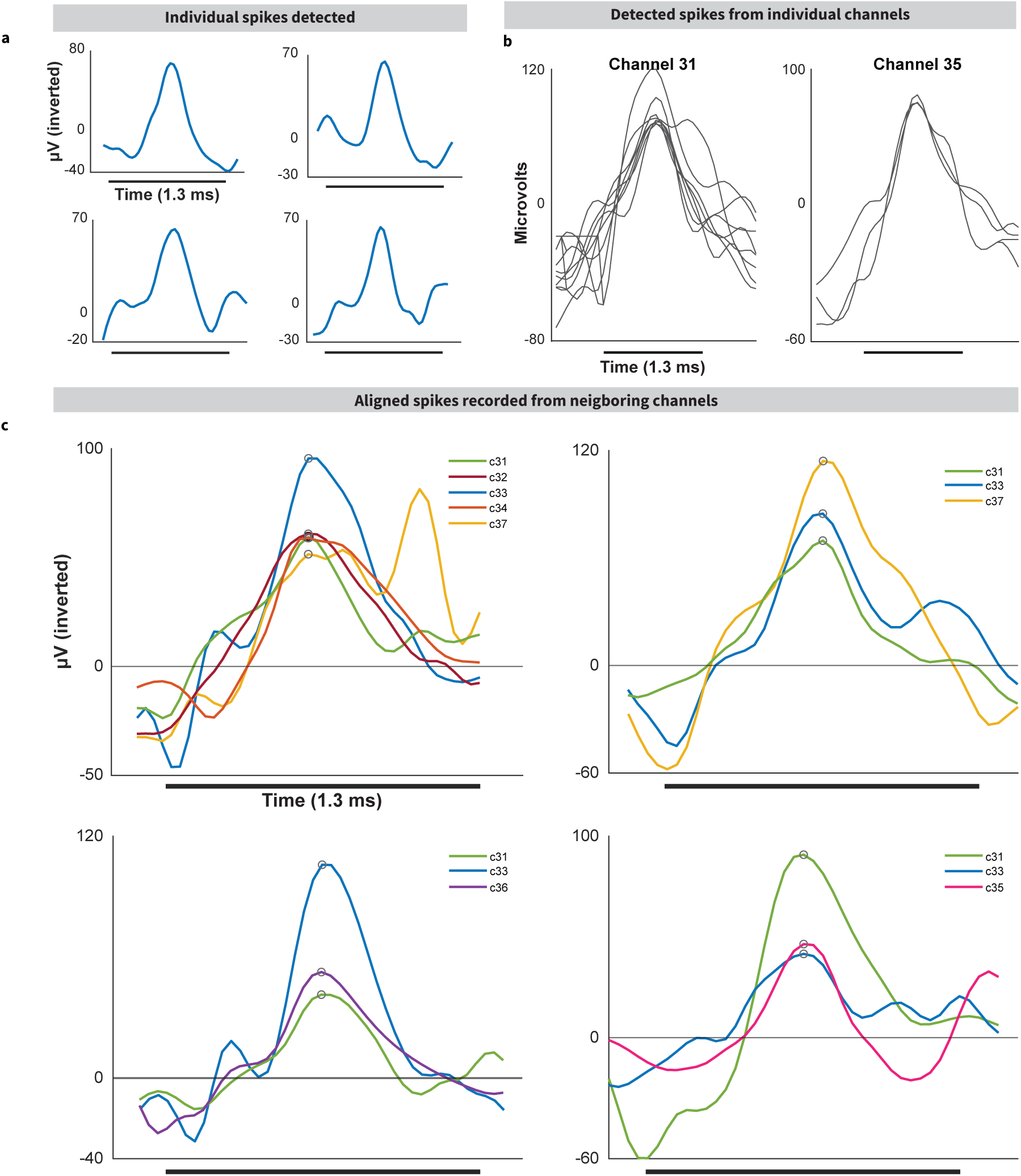
Electrophysiological recordings (cont.). **a.** Isolated spike waveforms detected from individual channels. **b.** Spikes which were detected across individual channels, overlaid and aligned to show the average waveform. **c.** Spikes detected across various neighboring channels. Spikes were aligned to the highest-value sample (circled) and overlaid to show that the general waveform shape is conserved across at least three channels corresponding to neighboring electrodes. This process can be expanded to enable spike sorting.

**Supplemental. Fig. 5.**
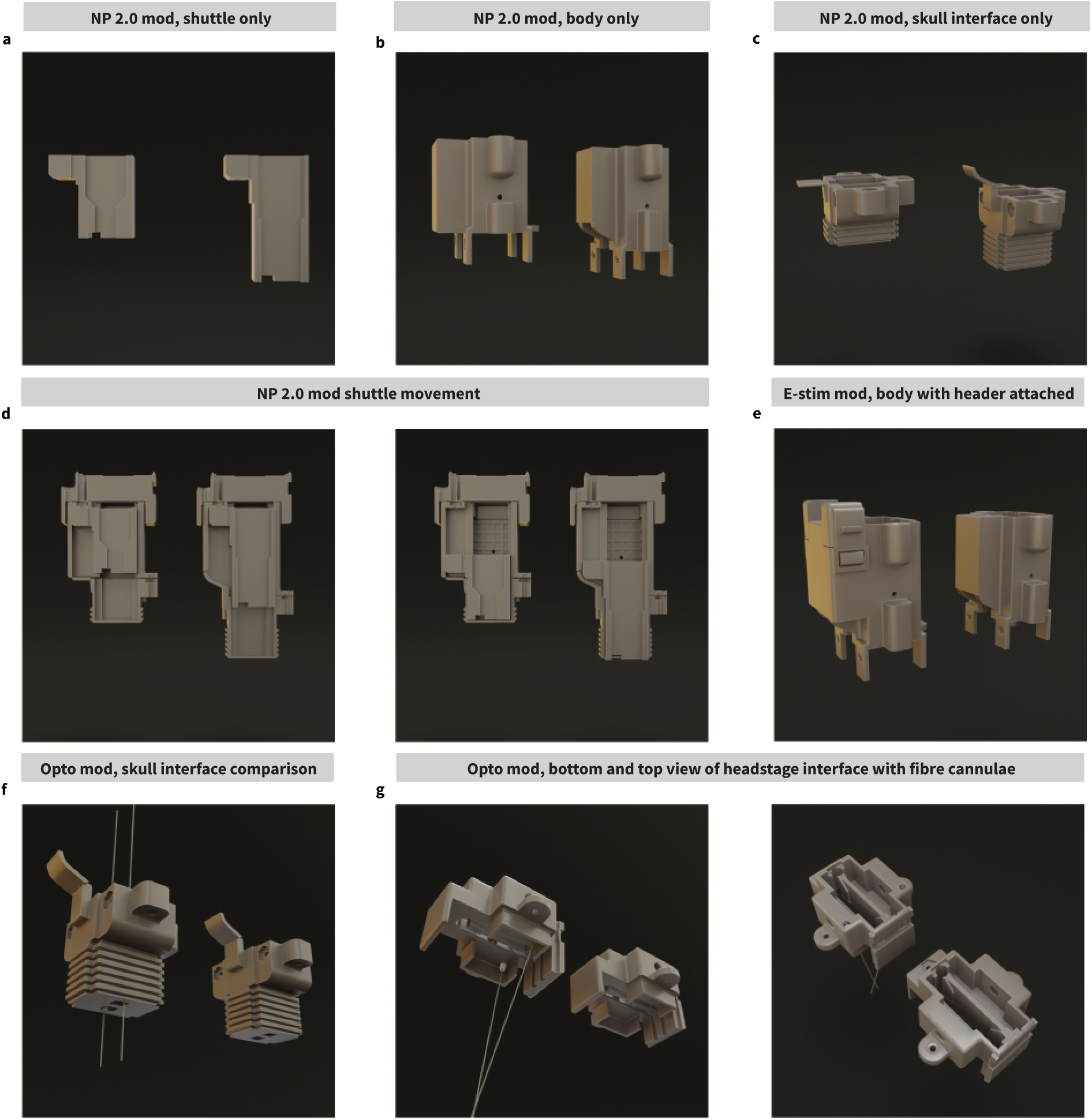
Proof-of-concept modifications, extended. a-c. Side-by-side comparisons of each of the modified pieces that make up the Neuropixels 2.0 (NP2.0) modified implant; modified pieces are on the left, and the original pieces are on the right. The shuttle (**a**) was shortened, and the shuttle bed was altered to accommodate the shorter NP2.0 probe. The implant body (**b**) was also shortened slightly, and the drive screw rail was elongated to ensure that the screw mechanism and shuttle are covered. The skull interface (**c**) was also shortened, and the drive screw rail cover was modified to fit the new shape seen on the implant body. **d.** Side-by-side bisected view of the NP2.0 and original implants showcasing the shuttle’s ability to retract into the implant (left panel) and be lowered fully (pictured right). e. With the addition of pin headers to implant body for the electrical stimulation implant modification, we have also designed a header connector (left) with a latch that secures the connector in place and may resist some pulling. **f.** Bottom view of the optogenetic mod skull interface. The cannulae are inserted from the inside of the skull interface piece and hold an optical fiber 1.25 mm away from the probe laterally, and 0.5 mm away from the probe tip vertically. **g.** Bottom (pictured left) and top (pictured right) views of the headstage interface piece made for the optogenetic mod. Cannuale are inserted into the piece to direct the optical fiber through one side of the implant body to avoid excessive bending and possible breakage. On the top part of the piece, the recessed cannulae create a connector receptacle for optogenetic equipment.

