## Supplemental Methods; Supplemental Notes 1 through 4 for "A modular, adaptable, and accessible implant kit for chronic electrophysiological recordings in rats"

#### Calculating probe location on the vertical axis

***Adjustments as a function of drive screw turns.*** By dividing the rotations of the drive screw in the implant's screw mechanism, one can control the distance that the shuttle travels along the shaft of the screw. If the amount of distance travelled per whole rotation is equal to the pitch of the drive screw, subdividing the rotations would divide the distance proportionally. Such granularity also slows the shuttle down significantly, which can prevent excessive stress on the probe shank as it moves through tissue. We can model the distance the shuttle would travel along the drive screw shaft as a function of turns at the drive screw.

$\mathrm{D} = \mathrm{P}\,\mathrm{T}_{\mathrm{Dr}}$ *(Equation 1)*

Where:

- $D$ is the distance in millimeters,
- $P$ is the drive screw pitch,
- $T_{Dr}$ is the direct turns of the drive screw

For example, if one would wish to make the shuttle and probes travel 0.06 mm along the shaft of the drive screw by hand, we can figure out how many turns it would take to achieve this by solving for $T_{Dr}$ if we know that the drive screw pitch is (0.3 mm):

$$T_{Dr} = \frac{D}{P} = \left(\frac{0.06}{0.3}\right) = 0.2$$

Meaning that it takes 0.2 turns of the drive screw to make the shuttle travel 0.06 mm along the drive screw. This is not achieved easily when done by hand, in most cases.
To achieve these precise movements, we developed the "Kepler screwdriver" (or simply *"Kepler driver"* or *"Kepler"*), a mechanical screwdriver designed with cascading planetary gears, each with a gear ratio (GR) of 5:1, resulting in a total GR of 25:1 (**see section *"Gear ratio calculations"***). This means that it takes 25 whole turns at the input of the system to complete 1 rotation at the output. With this GR, we can figure out the distance traversed by the shuttle along the shaft of the drive screw as a function of turns at the input of Kepler ($T_K$) instead of turns at the drive screw directly:

$D = P\frac{T_K}{GR}$ (*Equation 2)*

Where:

- D is the distance in millimeters,
- P is the drive screw pitch,
- $T_K$ is the turns at the input of Kepler,

- and GR is the gear ratio of Kepler.

Expanding upon the previous example, if we wish for the shuttle to travel the same 0.06 mm, we can solve for $T_K$ if we know that the gear ratio of the Kepler driver is 25:1 (25/1) and the drive screw pitch is 0.3 mm:

$$T_K = \frac{D}{P} GR = \left(\frac{0.06}{0.3}\right)\left(\frac{25}{1}\right) = 5$$

This tells us that we can make the probe and shuttle travel 0.06 mm within the implant by turning Kepler only 5 times, which is much easier to do than the 0.2 turns of the drive screw that we calculated above.

***Probe location during and after surgery.*** We can estimate where the tip of the probe is while the probes are inside the brain tissue through a series of calculations that can be made during and after surgery. During surgery, we first figure out the distance $L_0$ the shanks must be drawn to reach a target implantation depth to have a known starting point. This is a function of the target initial implantation depth ($DV_0$), skull thickness ($A$), and the implant's thickness and the bottom-most wall ($B$):

$$L_0 = |DV_0| + A + B \qquad \textit{(Equation 3)}$$

Where:

- $L_0$ is how far the shank must be drawn to achieve an initial implantation depth of $DV_0$,
- $|DV_0|$ is the absolute value of the target initial implantation depth,
- $A$ is the skull thickness, which is measured directly using callipers,
- And $B$ is the thickness of the implant wall at the bottom

For example, to implant at a depth of $DV_0$ = -1.4 mm on a rat with a skull thickness of $A$ = 0.7 mm, and assuming the implant has a wall thickness of $B$ = 0.5 mm, the shank must be drawn $L_0 = |-1.4| + 0.7 + 0.5 = 2.6$ *mm.*

To then figure out how many *direct* drive screw turns it takes to draw the shank to this length, we must plug $L_0$ from ***Equation 3*** into ***Equation 1*** as D and solve for $T_{Dr}$.

$$T_{Dr} = \frac{D}{P} = \frac{2.6}{0.3} = 8.67\ turns$$

We use a similar concept to know where the tip of the shank might be after each micro-adjustment, but instead of using ***Equation 1*** to calculate the direct drive screw turns, we use ***Equation 2*** to calculate the turns using the Kepler screwdriver since that gives us the most precise adjustment.

For example, if we want to make an adjustment of $D$ = –0.06 mm into the brain tissue, we can plug the absolute value of $D$ into ***Equation 2*** and solve for $T_K$. We know that the drive screw pitch $P$ = 0.3 and that the gear ratio of Kepler is $GR$ = 25/1, thus:

$$T_K = \frac{|D|}{P} GR = \frac{0.06}{0.3} 25 = 5\ turns$$

Finally, the approximate depth *DV* at which the tip of the probe shank is located within the brain after surgery is the initial implantation depth ($DV_0$), plus the sum of each micro-adjustment distance ($D_i$ ) made post-surgery.

$DV = DV_0 + \sum_n^1 D_i$ *(Equation 4)*

In this way, we can keep track of the shank depth several days after implantation and know how far we need to drive the probes into the brain to reach a target recording area. This information can then be used to calculate where a specific row of electrodes on the electrode array shank is located based on the electrode pad size and spacing between pads. We have created an Excel sheet when one may log each micro-adjustment to keep track of where the tip of the shank is over time and have made it available on GitHub (https://github.com/rjibanezalcala/EXPLORE/tree/main/SURGERY_KIT).

### The Kepler screwdriver

***Overview.*** The Kepler screwdriver, or simply "*Kepler driver" or "Kepler"* (**Fig. 1b**) is a modular (**Fig. 3a**), specialised screwdriver with a two-stage planetary gear system (**Fig. 3b-e**) encased within. It is compatible with any screwdriver bit with a 4 mm hexagonal shaft (**Fig. 3f**). The gear mechanism divides the speed and amount of turns at the input knob by 25 thanks to its gear ratio (**see "Gear ratio calculations"**). This is to divide the turns of the implant's drive screw to increase precision of any adjustments made to the vertical position of the probes, as well as minimise the possibility of human error when tuning the screw (**see "Preventing miscounting of turns"**). We provide a full build guide for the Kepler driver in **supplemental note 2**. All files related to the Kepler driver can be found in our GitHub repository (https://github.com/rjibanezalcala/EXPLORE/tree/main/KEPLER).

***Gear ratio calculations.*** In a gear-driven system, the proportion of output rotations to input rotations is called *turn ratio*, determined by the *gear ratio* (*GR*) of the system; herein, we will simply refer to the GR for simplicity. In the simplest spur gear arrangement, the GR of the system is determined by the number of teeth on the driver gear (input) and the teeth on the follower gear (output). The number of teeth on a gear also determines the diameter of the gear itself because since the two gears are expected to fit into each other, both the size of the teeth and the space between teeth must allow for the teeth on both gears to interlock. Therefore, if a driver gear has less teeth than the follower gear it will also be smaller and will complete several rotations before the follower manages to complete one, if the output of the system is placed on the follower gear, the input rotational movement at the driver gear will be slowed down at the output. However, a spur gear arrangement with large GR requires gears with many teeth and can thus become cumbersomely large and difficult to produce. Because the *Kepler driver* was meant to be portable and possibly used during rodent surgeries, we were constrained on the size of the device.

To make the mechanism compact, we decided on implementing a concentric, cascading planetary gear arrangement. This minimises the amount of space that the gears take and facilitates 3D printing of the entire screwdriver using a commercially available SLA printer (resin printer) to cut down on the cost of material. The whole mechanism consists of two planetary gear assemblies where the output of one is transferred to the input of the other. Cascading planetary gears multiply the GR of each stage, resulting in a much higher GR in a compact form factor. It also avoids plane movement transferring or offsetting the output from the input drive; movement is transferred vertically directly down the centre of the system to the output. A planetary gear is composed of an outer ring gear within which a carrier rests, and on the carrier are several planetary gears and a sun gear in the center.

In our design, the input to the mechanism is the sun gear and the output is the carrier, while the ring gear is held in place. For a setup like this, the inverse of the GR is given by the equation:

$$\frac{1}{GR} = \frac{S}{(R+S)} \qquad \textit{(Equation 5)}$$

We sought to achieve a GR of 25:1 because this would give us the smallest distance per turn that was useful for our purposes. This would then mean that 25 rotations at the input equals one rotation at the output, which when using a drive screw mechanism with a pitch (spacing between threads) of 0.3 mm, the smallest distance an object would travel along the shaft of the screw would be 0.012 mm (0.3 mm / 25 = 0.012 mm, **see Supplemental Note 4** for materials). To achieve this GR, we chose the following gear specifications:

| | **Teeth** | **Diameter (mm)** |
|---|---|---|
| **Planet gears (x3)** | 12 | 10.87* |
| **Sun gear** | 8 | 8 |
| **Ring gear** | 32 | 33 OD, 29.1* ID |

***Supplemental table 1 |*** *Number of teeth and diameter for each gear that comprises each of the two planetary gear assemblies of the Kepler driver. Diameters marked with a '*' were adjusted to fit our 3D printer's tolerance, original dimensions are 1)* ***planet gear:*** *12 mm, and 2)* ***ring gear:*** *ID 32 mm. 'ID' is shorthand for 'internal diameter' and 'OD' is shorthand for 'outer diameter'. OD value is not important for planetary gear function but is important for defining the overall diameter of the Kepler driver.*

If we plug in our values for the gear teeth, we get the gear ratio for each stage of the screwdriver's mechanism.

$$\frac{1}{GR} = \frac{S}{(R+S)} = \frac{8}{(32+8)} = \frac{1}{5}$$

By cascading the two planetary gears, the GR of each stage is multiplied, resulting in a gear ratio of…

$$GR = \frac{1}{5} * \frac{1}{5} = \frac{1}{25}$$

Although implementing cascading planetary gears does increase the complexity of the device and the number of components to print, everything can still be printed over several 3D printing jobs with the same 1 litre cartridge of resin. We have also provided an assembly guide and all the STL files for the screwdriver to guide users through the process of building the device (**see Supplemental Note 1**).

***Preventing miscounting of turns.*** Making fractions of a turn easily visualizable prevents miscounting, especially after long surgeries. We designed an optional mechanical counter that can be attached to Kepler (**Fig. 3g**). The counter simply transfers the input rotations to a 10:1 planetary gear (sun gear: 5 teeth; planet gear: 20 teeth; ring gear: 45 teeth) where the carrier is the output and the ring gear is fixed (**Fig. 3h,i**). A disk which explicitly indicates whole, and half rotations is attached to the carrier, and thanks to the gear system, will only rotate a 10th of a turn per full rotation of the input knob. The rotational movement of the whole counter mechanism is directly transferred to the input of the first 5:1 gear stage of the screwdriver, so the input to the screwdriver goes unchanged. The knob for the counter has 0.25, 0.5, 0.75 indicators as a reference for the fractions of a rotation. The counter attaches to the device at the top via a modified version of the original enclosure which can be easily replaced. This thus continues the theme of modular design which the implant itself features.

### Probe testing

***The saline bath.*** To test probes that have already been loaded into an implant, we designed a saline bath (**Fig. 1c**) which can be 3D printed and is adapted to our implant's form factor. The bath features a tray to hold saline in, two copper plates which serve as stimulating electrodes which flank the probes on both sides connected to standard banana connectors, and small receptacles distal to the probes where the ground electrode(s) can be placed during testing. To hold the implant, we included a miniaturized holding block specific for the saline bath where the implant can rest while still allowing the shanks to be lowered into the saline. This block is separate from the bath (**Fig. 4f**) to allow for adaptation to different implant shapes. When placed onto the bath, the block holds the implant securely above the saline. This setup allows for noise and gain testing on the probes as a troubleshooting step, or as a peace-of-mind step to take before implantation onto a live rat. Because the holding block is not built into the bath, one may remove it and test the probes inside the implant or by themselves without the implant under a stereotaxic device (**Supp. Fig. 2e, f**). This can be achieved by holding onto the shuttle and probes using our shuttle holder (**Supp. Fig. 2f**) or directly holding the probes using the manufacturer's dovetail holder (https://www.neuropixels.org/probes-np1-0) if the probes have a metal cap. Saline bath assembly can be found in **Supplemental Note 2**. The saline bath and stereotaxic holders can be found on GitHub (https://www.github.com/rjibanezalcala/EXPLORE).

***Gain tests.*** Testing of the probes was done using our custom saline bath (**Fig. 1c**) following the instructions in the Neuropixels 1.0 probe user manual *before* a probe was implanted. To test gain, a built implant with one fully retracted mounted probe was placed on the mini holder block (**Fig. 4f**) with external reference and ground connected to an Iso-flex's electrical stimulation isolator's (see *"Materials list"* section *"Other hardware"*) ground terminal. The block was then slotted onto the saline back above two stimulating electrodes. The bath was filled with sterile saline (0.9 g NaCl per 100 mL PBS) and the two banana connectors (one red for the cathode and one black for the anode) were plugged into the stimulator. The probe's Zero Insertion Force (ZIF) connector was mated with one headstage's Flat Printed Circuit (FPC) connector and the headstage was plugged into a Neuropixels card on a PXIe-1082 Chassis. The stimulator was set to deliver a current of 260 µA and was connected to a Master-9 (see materials list section *"Other hardware"*) with a pulse train routine of 100, 100 µs square pulses, spaced 250 ms apart for triggering; this setup was left on standby until needed. After an acquisition was started through software (SpikeGLX, Open Ephys GUI, or Bonsai), the probe was lowered into the PBS by turning the drive screw clockwise until at least the first 3 mm of the probe shank were submerged; a change in the recorded signal from large to low-amplitude noise confirmed that the recording electrodes had been submerged. Nominal gain was set to 1000 on both AP and LFP channels via software. The Master-9 was then triggered to start the pulse train routine and spiking patterns matching in frequency to the pulse train were observed on the acquisition window with little background noise. The stimulation amperage was adjusted to avoid saturating the recording equipment if the positive and negative peaks of the signal were clipped. After measuring the recorded signal's amplitude, we compared the recording to the true value set on the stimulator and calculated the gain by dividing the measured amplitude over the true amplitude. As indicated in the user manual, the resulting gain should not deviate more than 10% of the value set on the electrodes.

***Noise tests.*** Noise tests were not performed because we did not have a faraday cage on hand, which would prevent us from appreciating the true noise characteristics of the system and bias our noise tests. Nevertheless, we deemed noise levels acceptable if we were able to observe the stimulation signal with the external reference placed in one of the ground receptacles located distal to the stimulating electrodes on the saline bath, or after changing the electrical reference of the probes from external to tip.

### Surgical and post-surgical procedures

***Tracking probe depth during and after surgery.*** To know the vertical location of the probes (*DV*), we first inserted the probe shanks at a known depth ($DV_0$). To do this, we measured the skull thickness by directly measuring the bone extracted from the trephine hole using calipers. Although measuring with calipers is preferred to prevent damage to the probes, this distance may also be measured with a stereotaxic device by subtracting the DV at which the probe tips touch the bone at the edge of the trephine hole, from the DV at which the probes touch the brain tissue. The thickness of the bottom wall of the skull interface (typically 0.5 mm) is then added to the skull thickness (**see 'Calculating probe location on the vertical axis', Equation 3**), and the probes are drawn out to this total distance by turning the drive screw counterclockwise the appropriate amount of times (**determined by Equation 1**). This length of the probe shank is then implanted into the brain. Any micro-adjustment (**see 'Post-surgical micro-adjustments'**) made after surgery ($DV_{1\text{-}n}$) is also tracked and cumulatively added to the initial implantation depth (**Equation 4**). For more information, see **Supplemental Note 3**. We have created an Excel sheet when one may log each micro-adjustment to keep track of where the tip of the shank is over time and have made it available on GitHub (https://github.com/rjibanezalcala/EXPLORE/tree/main/SURGERY_KIT).

***Implantation surgery and post-surgical care.*** To ensure that the implanted probes are useful for as long as possible, several considerations must be considered. To start, the implant must be affixed to the rat's head in the most stable way possible to prevent it from being torn off by the rat. To do this, we scored the skull and added various anchor screws around the implant (**Fig. 5a, b**), then covered them in dental cement so that the whole implant had a strong foundation (**Fig. 5c**). The screws must be driven deep enough into the bone that they are secure, but not deep enough to cause damage to the brain. This in combination with a large foundation of dental cement covering the entirety of the exposed bone keeps the implant in place after surgery (**Fig. 5d, e**). After the probes have been introduced into the tissue, applying Vaseline around the "foot" of the implant where it contacts the skull can prevent discharge and blood from making its way into the body of the implant to prevent buildup from forming along the shuttle rails and essentially gluing the shuttle in place hindering further vertical adjustment of the probes. At the end of the surgery, we cauterized the remaining tissue around the implant with silver nitrate to prevent the tissue from regenerating under the dental cement and lifting the entire implant from its anchor. Post surgical care is also important in not only keeping the animal comfortable and healthy but also prevents implant failure. To prevent infections, provide pain relief and hydrate, animals were given routine broad-spectrum antibiotic injections (Enrofloxacin) and an injected NSAID (Meloxicam) in 10 ml of Ringer. Then, to reduce itching and locally disinfecting the surgical site, we routinely applied Chlorhexidine and a triple antibiotic on the live tissue around the implantation site. A combination of nystatin (antifungal), neomycin (antibiotic), thiostrepton (antimicrobial), and triamcinolone (steroid) was also used to treat this area. Hyaluronic acid can later be applied to keep the area moisturized after the tissue has scarred (**see Supplemental note 3**).

***Home cage modification.*** Modifications to the animal's home cage can help extend the lifespan of the implant by creating an environment that is comfortable to the rat with a large object attached to its head. A cage which is twice as tall as a standard rat home cage will allow the animal to move and rear freely. A taller cage necessitates giving access to water from the side of the cage instead of from the top. Additionally, reducing the length of the waterspout that protrudes into the cage helps eliminate points where the animal's head can get stuck on which could lead to pulling on the implant until it rips off. The waterspout should not protrude more than 20 mm into the cage and should be situated close to the bottom of the cage or high enough to prevent the animal from using it as leverage. For more information, see **Supplemental Note 3**.

### Post-surgical micro-adjustments

Immediately after surgery, it is likely that there will be a very aggressive inflammation response in the tissue. Over time, scarring may occur around the electrode shank which may put the probes at risk of breaking if the probes are lowered too quickly after surgery. Scar tissue forming around the electrodes may also degrade the recorded signal over time. To combat the dangers of moving a fragile object through scarring, changing tissue and the natural degradation of the signal due to brain tissue immune response, we leveraged the adjustable nature of the movable shuttle by making a series of post-surgical micro-adjustments using our Kepler screwdriver. With this approach, we were able to slightly adjust the position of the probe shank within the brain tissue in a slow and controlled manner to avoid breaking the shank and keeping track of where the tip of the shank is after each adjustment (**see *'Calculating probe location on the vertical axis'***). We were able to reach our target at the Dorsal Striatum (–3.45 mm DV) within 3 weeks (after an initial implantation depth of –1.5 mm) by making 100 – 300 µm adjustments 2 – 3 times per week. Afterward, only 50 µm adjustments were made every other week or as soon as the recorded signal had degraded to a degree where spikes were undiscernible from the local field potentials (LFP) to maintain a relatively good SNR for at least 3 months after surgery.

#### Grounding considerations

***Ground needs to be stable and quiet.*** Typically, electrical ground is needed to aid in the denoising of electrophysiological signals during the pre-amplification stage of the signal processing pipeline. Because voltages at the input of an amplifier are measured with respect to ground, ground must be as free of noise as possible to avoid unintentionally introducing artifact signals into the system, as this artifact will be present at the output of the amplifier. Additionally, grounding serves as a low-impedance path for current to flow and acts as shielding against electromagnetic interference (EMI). For this reason, a quiet and stable grounding circuit must be present in any electrophysiological recording equipment, as the very low amplitude signals (roughly 10 – 100 µV) produced by neurons are extremely susceptible to large-amplitude noise and artifacts which may range from 1 mV to several volts, depending on the source. If no grounding is present at all, the entire recorded signal will at least be corrupted by 60 Hz line noise, and any other EMI present in the recording environment (for example, from the host computer) which impedes spike sorting later down the pipeline.
To ground the probes within the implant, we included a simple grounding circuit within the implant, which extends out to a small copper plate through which a screw is placed (**see Supplemental Note 2**). This not only connects the probes and screw together but also makes it so that the surgeon may place the ground on whichever screw on the skull. We chose copper for this plate because unlike stainless steel, which is what screws tend to be made of, it is a low-impedance metal and takes solder well, which is important for affixing the ground wire. Any other similarly conductive material would be appropriate (such as a tin-copper alloy) if it solders well. For the wire, a decently conductive material with low impedance such as Silver should be chosen to allow current to pass onto the ground, however, it is important to note that the total impedance of the ground circuit will be equal to the summation of resistances of each element of the grounding circuit, provided that only resistive elements such as the wire, plate, and screw exist in the circuit and no other capacitive or inductive elements are introduced. Maintaining a clean, simple, low-impedance ground will aid in keeping recorded signals as free of artefacts as possible.

***Avoiding muscle tissue on ground circuit.*** Because muscle tissue is electrically active, it is common for it to be a prominent source of artifact in electrophysiological recordings when it meets any part of an electrical circuit. For electrode arrays, the reference, or particularly, the grounding circuits are the most susceptible as they tend to be outside of the tissue they are recording from and may contact electrically active tissue, particularly muscle. Voltage readings from muscle activity result in signals that are magnitudes larger in amplitude than extracellular neuronal signals and may contain similar frequency components. This means that a recording from brain tissue can easily be obscured by electromyographic signal patterns both in the voltage and frequency domains, which can complicate filtering, hinder spike detection, and prevent accurate spike sorting later down the pipeline.

There are a variety of muscles located on the rat skull. Of special importance, however, is the temporalis muscle, due to its location. Most procedures for brain electrophysiology implants, including ours, require reflecting the temporalis muscle at the dorsal crest of the rat skull to access the bone and perform a craniotomy to implant the probe shanks into. This large muscle extends beyond this area however, and is involved in jaw closing (https://peerj.com/articles/448/), which means it will be very active throughout any recording session as a rat bruxes, grooms, or eats. Avoiding electrical contact with this muscle is crucial for clean recordings that are useful for spike sorting algorithms.

To avoid contacting muscle with any component of the grounding circuit on the probes, we create a large incision on the rat's head which extends from between the eyes to just behind the ears. We then reflect the skin and muscle from the implantation site and scrape the connective tissue off of the skull to make enough space for grounding screws and ensure they are not contacting any surrounding tissue. We also coat the screws in dental cement to shield them from the tissue. This prevents the ground screws from contacting any surrounding muscle and provides a sufficiently "quiet" ground for the probes, which prevents (or at least partially prevents) electromyographic artifacts from contaminating our recordings.

***Avoiding ground loops.*** Having multiple recording devices (probes) on the same rat may present a challenge with grounding. Avoiding ground loops is extremely important so that each probe has a quality reference point from which to measure voltage from, which in turn prevents the recorded signal from being corrupted by EMI. Ground loops occur when 2 or more devices are interconnected through their ground circuits. The resulting ground circuit creates a physical closed loop, which acts as a single-turn secondary winding of a transformer with the primary winding being the summation of all the magnetic fields produced by neighboring devices. When oscillating (50 – 60 Hz) ambient magnetic fields pass through the ground loop, an oscillating current is induced and conducted through the grounding wire, into the recording device, and consequently introducing EMI into the recorded signal. Because of the low resistance of the wire, even a small ambient magnetic field can induce a current in the ground loop which can significantly interfere with the low-amplitude recordings. In audio equipment, the induced EMI can present itself as a constant low-frequency hum.

To prevent ground loops when two probes are present on the implant, it's important to give each probe its own separate ground circuit. One can achieve this by simply avoiding interconnection of each probe's ground screw during surgery. If more than one screw is included in a probe's ground circuit, the same applies; that probe's ground circuit must not be connected to the other probe's ground. In a single probe setup, limit the number of ground screws in the circuit. Each probe, whether in a single or double probe configuration, should have at minimum one ground screw placed in an area distal to the recording site, and away from any tissue other than bone. Accidental ground loops, where the same wire comes into contact with itself, are also prevented by using wire that has been coated with an electrically insulating material such as Perfluoroalkoxy (PFA). Additionally, as mentioned in a previous section, minimize the surface area that the ground circuit covers as long stretches of (un-looped) wire can also act as an antenna and pick up EMI. The ubiquitous nature of ambient magnetic fields makes avoiding ground loops an extremely important step in denoising a recorded signal; although EMI can be filtered away, it is always best to avoid picking it up altogether to get the cleanest, most artifact-free signal possible.

#### Recording and signal processing

***Recording environment and procedure.*** 50-minute recording sessions were done on two implanted rats separately over 64 days for one, and over 112 days for the other. Rats were anaesthetized prior to recording using vaporized Isoflurane in order to connect the headstage safely, then individually recorded freely moving in a dark room from within their home cages with the lid removed. Rats were supervised but with minimal interaction between the rat and the experimenter. The data acquisition equipment used was a PXIe-1082 chassis with a Neuropixels PCIe card recording from one Neuropixels headstage and Neuropixels 1.0 probe on each rat.

***Signal processing.*** Recorded signals were pre-processed and visualized in MATLAB using a combination of custom code and plugins (**Fig. 6a**). The signal processing chain consisted of four ordered processing stages: 1) artifact identification and removal using visual inspection and thresholding (z-score = 10 threshold; defined as a number of standard deviations from the entire background signal on the processed channel), 2) common local field potential (LFP) averaging (referencing 7 channels above and/or below the processed channel), 3) band-pass filtering (with 2$^{nd}$ order Butterworth filter, filter pass-band = 300 : 5000 Hz), and 4) spike extraction (using a voltage threshold = 80 µV). Source code for this signal processing chain can be found on GitHub (https://github.com/lddavila/cluster_neuronspikes/tree/main/Porting%20Open%20Ephys).

***Spike detection and alignment.*** Spike detection and alignment was done offline in MATLAB over one channel at a time (https://github.com/lddavila/cluster_neuronspikes/tree/main/Porting%20Open%20Ephys). Spikes were detected with a threshold of -60 µV, inverted, then cut to 1.3 millisecond clips with maximum value of the spike as the median data point (**Fig. 6b, Supp. Fig. 4a**). Detected spikes were aligned afterward by plotting each spike centered at the maximum value of each (**Fig. 6c, Supp. Fig. 4b**). To relate the detected spikes to activity on neighboring channels, the timestamp of the highest-amplitude sample from each detected spike was saved and analyzed across the nearest 4 channels above and below (corresponding to neighboring electrodes). The same spike detection algorithm was run over these channels to find similar spiking activity in those channels (**Fig. 6d, Supp. Fig. 4c**).

#### Implant modifiability

***Modularity facilitates modifiability.*** We have chosen a modular implant design so that any adaptations to the implant can be made in as little time as possible by isolating only key parts of the implant instead of having to always make modifications to the general structure. For example, although our ‘base’ implant is designed around the Neuropixels 1.0 probe, the implant can be adapted to a 2.0 probe in under a day by modifying only the shuttle. Assuming the general dimensions of the shuttle are left unchanged, the modified shuttle can still fit in the implant body and be driven by the same drive screw mechanism. If desired, the implant can be shortened to accommodate the smaller 2.0 probes in an additional day or two, only adapting two additional implant pieces (see ***'Neuropixels 2.0 adaptation'***). In a similar fashion, individual implant parts can be modified to add stimulation electrodes and wires (see ***'Electrical stimulation modifications'***), or optical fibre cannulae to include optogenetic stimulation capabilities on-board the implant (see ***'Optogenetic modifications'***). Modified versions of the implant can be found in our GitHub repository (https://github.com/rjibanezalcala/EXPLORE/tree/main/IMPLANTS).

***Mesh-oriented implant editing.*** The open-source, 3D software *Blender* (version 3.4.0) was chosen over CAD software for the development of this project to provide an alternative method for modifying the base implant and in this way allowing for more flexibility. Blender's mesh-oriented 3D modelling makes modifications easy by enabling the user to modify individual sections on a mesh (*vertices, edges, or faces),* or by using non-destructive mesh editing through *Modifiers.* For example, if one wishes to alter the dimensions of the implant, the mesh can be scaled, or groups of vertices on the mesh can be moved around on the X, Y, or Z axes (see ***'Neuropixels 2.0 adaptation'***). On the other hand, with an *Additive Boolean modifier*, an attachment can be 3D modelled and then attached to the base implant mesh (see ***'Electrical stimulation modifications'***), or with a *Subtractive Boolean modifier,* holes and channels can be made on the implant to embed or made room for wires or optical fibre (see ***'Optogenetic modifications'***). These modifications can also be made in CAD software such as AutoCAD, SolidWorks, or the open-source FreeCAD by modifying the STL files directly, but the inclusion of Blender as a 3D model-editing platform extends the modifiability of our implant to one additional platform and give the most power to the end-user.

***Neuropixels 2.0 adaptation.*** Our implant can be easily adapted to the newer Neuropixels 2.0 probes by making quick and small modifications. As mentioned before, the shuttle is the main implant piece that needs modification to fit the 2.0 probe snugly on the implant. However, in this paper we have made modifications to the whole implant to provide a ready-made implant for the newer generation probes (**Fig. *7a***, modified implant pictured left). The first modification made was to the shuttle, where it was shortened, and the *probe bed* was modified to fit the new probe's form factor (**Supp. Fig. 5a**, pictured left). The next modification was on the *implant body.* The body was shortened slightly, and the drive screw rail was elongated to ensure that the shuttle is fully protected regardless of where it is located on the implant (**Supp. Fig. 5b**, pictured left). Finally, the skull interface was also shortened, and the *rail cover* was modified to fit the new screw rail on the body (**Supp. Fig. 5c**, pictured left). In both the original and modified designs, the probe and shank are protected fully when the shuttle is at its highest position (**Fig. 7b, Supp. Fig. 5d**). The result of this modification yielded a smaller implant, measuring approximately 3.6 cm in height compared to the 4.4 cm of the original implant (including the headstage interface and cap pieces). These three modifications were made over the course of two days and illustrate how the implant can be quickly modified to different probe form factors instead of needing to design an entirely new implant around a different probe which could take several weeks.

***Electrical stimulation modification.*** We have provided an example modification to the implant that includes a connector for adding electrical stimulation capabilities to the implant (**Fig. 7c**, pictured left). In this modification, the *implant body* is fit with a header pin receptacle (**Fig. 7d**, pictured left) and stimulation wire connectors (**Supp. Fig. 5e**). and the skull interface is modified so that 0.2 mm wire can stick out of the bottom of the implant (**Fig. 7e**). The setup allows for the header pins and wires to be attached from the top of the implant, then the wires are fed into the skull interface via the side holes and finally soldered to stimulation electrodes affixed to the bottom. In this example, only the implant body and skull interface are changed while every other part of the implant fits in place without any modifications.

***Optogenetic modification.*** The last modification to our implant that we made is one that allows for optical fibres and cannulae to be added to the build for experiments that include optogenetic stimulation. In this design, optical fibres come out of the bottom of the skull interface 1.25 mm away from each probe shank, 0.5 mm above the tip of the shank (**Fig. 7f**, pictured left). This is achieved through the addition of receptacles where cannulae (1.1 mm diameter cannula for 0.1 mm diameter optical fibre) are affixed (**Supp. Fig. 5f**). While the implant body and shuttle remain unchanged, the headstage interface is modified so that an additional two cannulae can be affixed on the top to create a receptacle to connect optical stimulators (**Supp. Fig. 5g**). The top receptacles are offset and slightly tilted to move the optical fibre out of the way of the probe ribbon when it is folded inside the implant body.

### Supplemental Notes

#### Supplemental Note 1: “Kepler screwdriver” building procedures

Assembly of the Kepler screwdriver, or simply *Kepler driver* does not require extra steps to post-process the 3D printed pieces such as drilling or sanding. It also does not require any fastenings to put together. However, we do recommend lubricating the planetary gears before putting the device together completely. We provide instructions on this within this assembly guide. Because this device is 3D printed, one may expect small imperfections on the components brought up by the printing process to cause minor issues in the device’s functionality. This can be resolved with repeated use of the device until movement of the components becomes smoother over time.

##### Printing the Kepler driver components

It is recommended that every component is printed using an SLA printer (photopolymer resin printer) at a high resolution (for example 0.005 mm layer height) to ensure that component dimensions are as close to the designed dimensions as possible. FDM printers may not be able to offer this resolution and, because of the nature of the printing technique, may introduce imperfections that might interfere with the device’s operation.

We were able to successfully print and assemble several Kepler drivers and replacement parts using a Formlabs Form 3 and Clear V4 resin (**see supplemental note 4 ‘Materials and sourcing’ under section ‘3D printer and accessories’**). A different resin which offers good resolution may be used instead if desired, however this was not tested by us. Keep in mind that Clear V4 resin is relatively brittle and may break if the device is dropped but will withstand regular day-to-day use under normal conditions. Finally, because Clear V4 resin has the tendency to deform during post-curing when prints are not very dense, we recommend not doing a post-cure; instead, simply washing the screwdriver components and letting them cure at room temperature over time (may take up to 7 days) will ensure that the shape of the components is preserved.

All Kepler driver components are available as STL files in our GitHub repository (https://github.com/rjibanezalcala/EXPLORE). Pre-made Preform (**see supplemental note 4 ‘Materials and sourcing’ under section ‘Software’**) projects are also provided in the same repository for downloading and printing directly. The components list is shown below (**Table S1.1, Fig. S1.1**), which indicates what components are needed and how many of them to print. Note that the addition of the counter module is optional, however we recommend adding it as it counts the number of turns made at the input which can reduce user error. If the counter module is added, make sure you make the replacements indicated in **Table S1.1**. This list can also be found in **Supplemental Note 4**.

| | | |
|---|---|---|
| **“Kepler” screwdriver** | Bottom casing | 1 |
| | Bottom casing shell | 1 |

| | | |
|---|---|---|
| **printable components** | Knob (small or large)* | 1 |
| | Bit adapter | 1 |
| | Middle indicator | 1 |
| | Top casing | 1 |
| | Top casing shell** | 1 |
| | Carrier | 2 |
| | Carrier spacer | 2 |
| | Planet gear | 6 |
| | Planet gear spacer | 6 |
| | Sun gear shaft*** | 1 |
| | Sun gear socket | 1 |
| | Sun spacer | 2 |
| | Ring gear | 2 |
| | Counter ring gear (optional) | 1 |
| | Counter carrier (optional) | 1 |
| | Counter carrier spacer (opt.) | 1 |
| | Counter planet gear (opt.) | 3 |
| | Counter planet gear spacer (opt.) | 3 |
| | Counter sun gear (opt.) | 1 |
| | Counter sun gear spacer (opt.) | 1 |
| | Counter knob (opt.)*<br>*Replaces regular knob* | 1 |
| | Sun gear shaft adapter for counter (opt.)***<br>*Replaces one regular sun gear* | 1 |
| | Counter face (opt.) | 1 |

| | | |
|---|---|---|
| | Counter shell (opt.) | 1 |
| | Top casing shell for counter module (opt.)**<br>*Replaces regular top shell* | 1 |

**Table S1.1 | *Kepler driver components list.*** This table shows all the 3D printable components necessary to assemble the Kepler driver. Counter components (marked as optional) are not required to use the Kepler driver; however, the counter module does provide some readability to the device and is recommended. Items marked with ‘*’ indicate that one may replace the other if the counter module is to be used. Thus, if the counter module will be added, print the *top casing shell for counter module* instead of the regular *top casing shell* and the *sun gear shaft adapter for counter* instead of the regular *sun gear shaft.* All listed components can be found in our GitHub repository at https://github.com/rjibanezalcala/EXPLORE/tree/main/KEPLER.

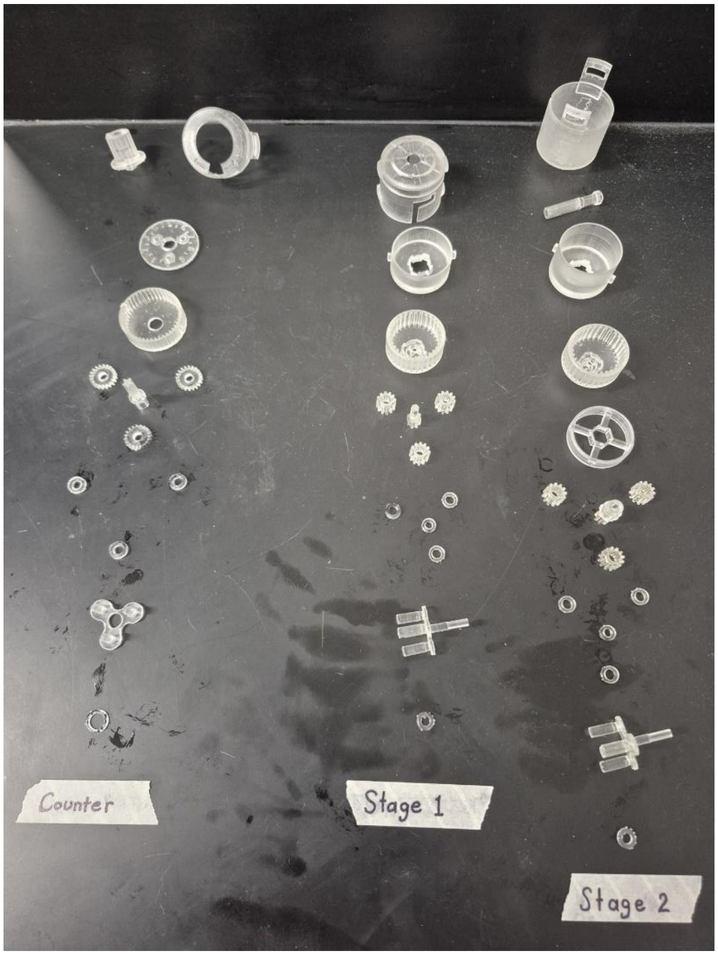

**Figure S1.1 | Kepler components.** The photograph shows all the Kepler driver components classified by what subsystem of the device they belong to. The top components encompass the outer part of the device (shell) and the inner components are toward the bottom.

**In summary:**

a. Print Kepler driver components using an SLA printer with high-resolution compatible resin (this type of resin tends to be runnier) at a resolution of at least 0.005 mm layer height.
b. Do not post-cure the Kepler driver components, instead simply wash them and let them cure at room temperature for a few days. If post-curing is still desired, we recommend limiting the cure time.
c. All 3D printable components can be found alongside ready-to-print Preform projects at https://github.com/rjibanezalcala/EXPLORE/tree/main/KEPLER.
d. The components list and quantities are listed above in **Table S1.1** or also under '"Kepler" screwdriver printable components' in **supplemental note 4**.

#### Assembling the planetary gear systems

**Relevant components (screwdriver):**

- Sun gear (shaft, and socket), 1 each.
  - Replace sun gear (shaft) with sun gear shaft adapter for counter if printing a counter module.
- Sun gear spacer (regular), total of 2.
- Planet gears (regular), total of 6.
- Planet gear spacers (regular), total of 6.
- Ring gear (regular), total of 2.
- Gear carrier (regular), total of 2.
- Gear carrier spacer (regular), total of 2.
- Middle indicator, total of 1.

**The main planetary gear systems**

The instructions below assume only the components listed under "Relevant components (screwdriver)" (**Fig. S1.2a**).

1. Slide a sun gear spacer into the middle peg on each carrier. Then, slide one planet gear spacer into the three surrounding pegs (**Fig. S1.2b**).
2. Place a sun gear (shaft and socket) in the centre peg on each carrier (**Fig. S1.2c**), and one planet gear into each of the surrounding pegs (**Fig. S1.2d**). Wiggle the sun gear a bit if this gives your trouble.
3. Slide a carrier spacer on the bottom side of the carrier (on the longer shaft side (**Fig. S1.2e**), then slide the whole assembly into a ring gear (**Fig. S1.2f**). Repeat for the other carrier and ring gear.
4. Test the planetary gear systems by turning the sun gear in both directions. Lubricate the gears as needed.
5. Make sure to take note which one has the sun gear shaft, and which one has the sun gear socket and middle indicator, the former will be the stage 1 gear system, and the latter will be stage 2.
6. On the stage 2 gear system, slide the middle indicator piece onto the sun gear socket so that it sits on top of the whole gear system (**Fig. S1.2g**).
7. Set both gear systems aside for now (**Fig. S1.2h**).

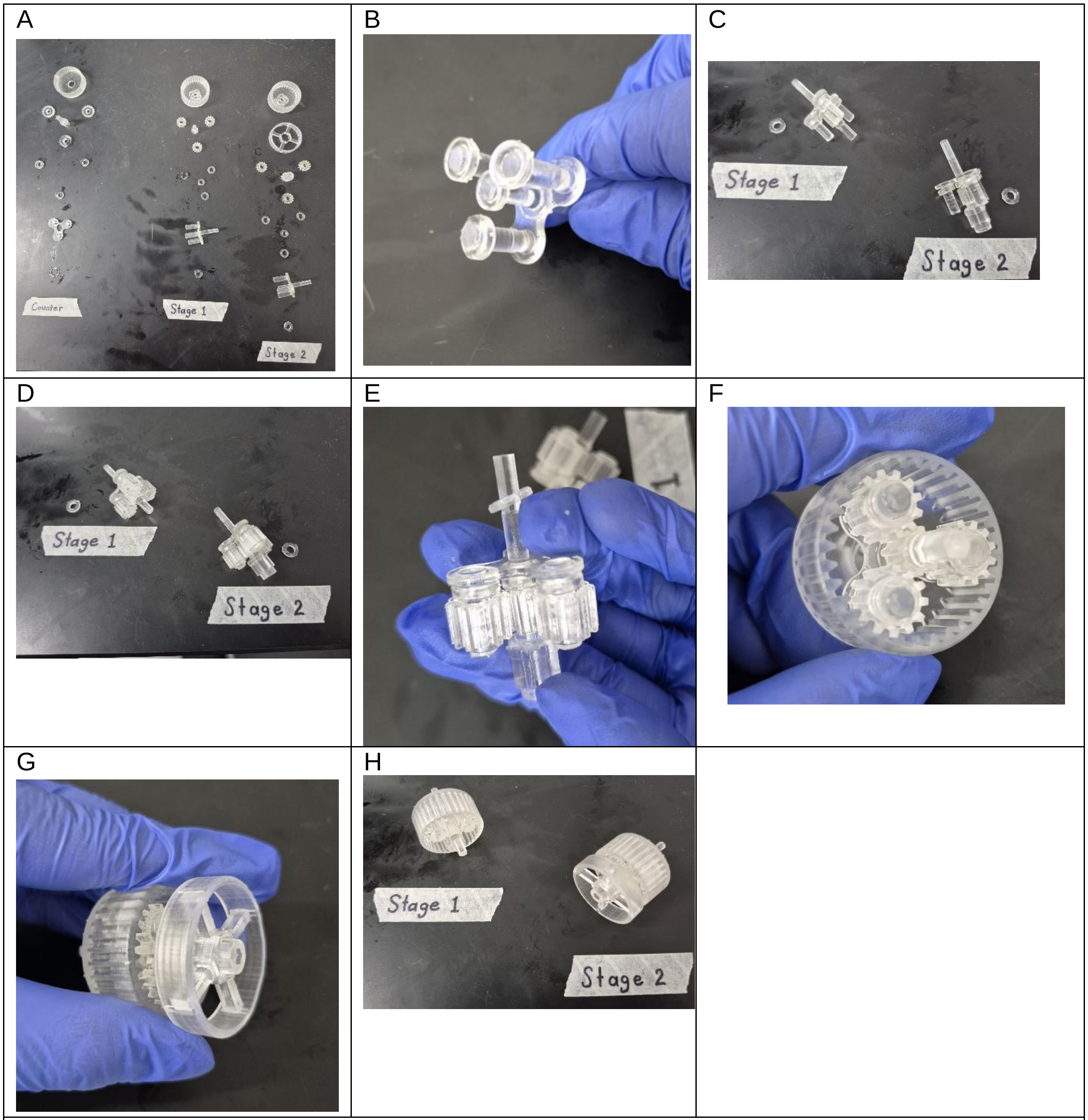

**Figure S1.2 | Stage 1 and 2 gear systems assembly. a.** Gear system components, separated by the subsystem they belong to. **b.** Placing the sun (middle) and planet gear spacers on the carrier. **c.** Placing the sun gears on the carriers. Stage 1 requires the sun gear shaft and Stage two the sun gear socket. Carrier spacer pictures off to the side of each carrier. **d.** Placement of the planet gears around the sun gear on the carrier. **e.** Placement of the carrier spacer on the bottom side of the carrier. **f.** Placement of the carrier assembly within the ring gear, completing Stage 1. **g.** Placement of the middle indicator on top of Stage 2's sun gear. **h.** Comparison between Stage 1 and Stage 2 gear systems.

#### The counter module

**Relevant components:**

- Sun gear shaft adapter (counter)

- Sun gear spacers (counter)
- Planet gears (counter)
- Planet gear spacers (counter)
- Ring gear (counter)
- Counter module planetary gear system
- Counter face
- Counter shell
- Counter knob

**The counter planetary gear system**

Follow steps 1 through 5 from the main planetary gear systems to construct the counter's planetary gear system (**see 'Assembling the planetary gear systems', Fig. S1.3a-d**).

**Assembly**

1. Place the counter face piece face up on top of the carrier pegs. Make sure that the pegs slot into place on the bottom of the counter face (**Fig. S1.3e**).
2. Hold the gear assembly by the side pegs on the ring gear and insert the whole assembly face-up into the counter shell, by aligning the side pegs to the small notches on the shell.
3. Push the gear assembly all the way up inside the shell, then twist it to follow the notches until everything clicks into place (**Fig. S1.3f**). The gear assembly should not rotate within the shell if it has been inserted correctly.
4. Insert the knob into place on top of the counter module assembly and rotate the gear system until the '0' marking aligns with the small arrow on top of the "number window", then re-insert the knob so that the '0.00' marking is on top (**Fig. S1.3g, h**).
5. Firmly push down on the knob so that somewhat stays in place. The knob may fall out as there is no fastening which holds it in place.
6. Set the counter module aside (**Fig. S1.3i**).

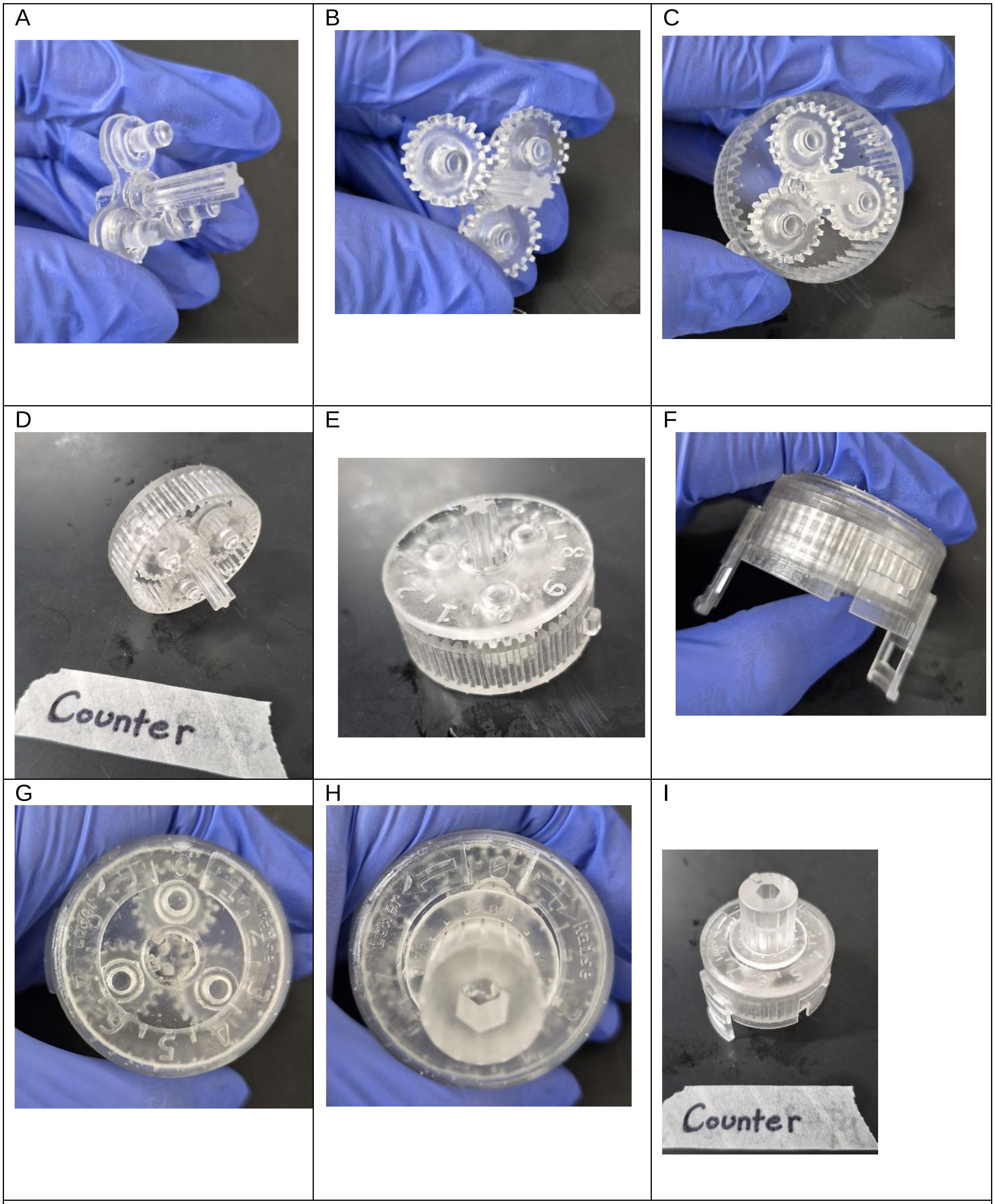

**Figure S1.3 | Counter module subsystem assembly. a.** Placement of planet spacers, sun spacer, and sun gear on counter carrier. **b.** Placement of counter planet gears on the carrier. **c.** Placement of carrier assembly inside of the counter's ring gear. **d.** Finished counter module gear system. **e.** Placement of the counter face on top of the carrier pegs. **f.** Sliding the counter gear system and face into its shell by aligning the pegs on the ring gear to the notches on the shell. **g, h.** Setting the counter system to zero (**g**) before placing the knob on top, also centred at 0 (**h**). **i.** Finished counter module.

#### The screwdriver body

**Relevant materials:**

- Knob (small or large), x1 (if not using the counter module)
- Top casing shell, x1 (if not using the counter module)
    - If using counter module, replace with the top casing shell for counter module
- Top casing, x1
- Bottom casing shell, x1
- Bottom casing, x1
- Bit adapter

**Assembly instructions:**

1. Insert the stage 1 planetary gear system (see ‘assembling the planetary gear systems’) into the top casing piece, and the stage 2 system into the bottom casing (**Fig. 1.4a, b**). Make sure you align the bottom of the ring gears to the patterns on both the casings so that the whole assembly slides fully into place.
2. Slide the bit adapter into the bottom casing shell from the inside (**Fig. 1.4c**). The bit adapter should not fall out when the shell is held right-side-up (the rat logo on the side should be right-side-up, **Fig. 1.4d**).
3. Slide the stage 1 gear system and bottom casing down the bottom casing shell so that the side pegs on the bottom casing go through the shell notches, twisting until everything slides into place (**Fig. 1.4e**). Set it aside.
4. Slide the stage 2 gear system and top casing in the same way (**Fig. 1.4f**).
5. Align the bottom shaft on the stage one gear system into the sun gear socket on the stage 2 system, then slide everything into place. Make sure that the side latches on sides of the two shells latch together (**Fig. 1.4g**).
    a. If a counter module is used, repeat this with the top latches on the top casing shell (**Fig. 1.4h**).
    b. If a counter module is not used, inset the regular knob on the top of the screwdriver.
6. Give the completed screwdriver a few turns to test if everything turns as it should.
7. Slide a 4 mm screwdriver bit into the bit adapter to use the Kepler driver (**Fig. 1.4i**).

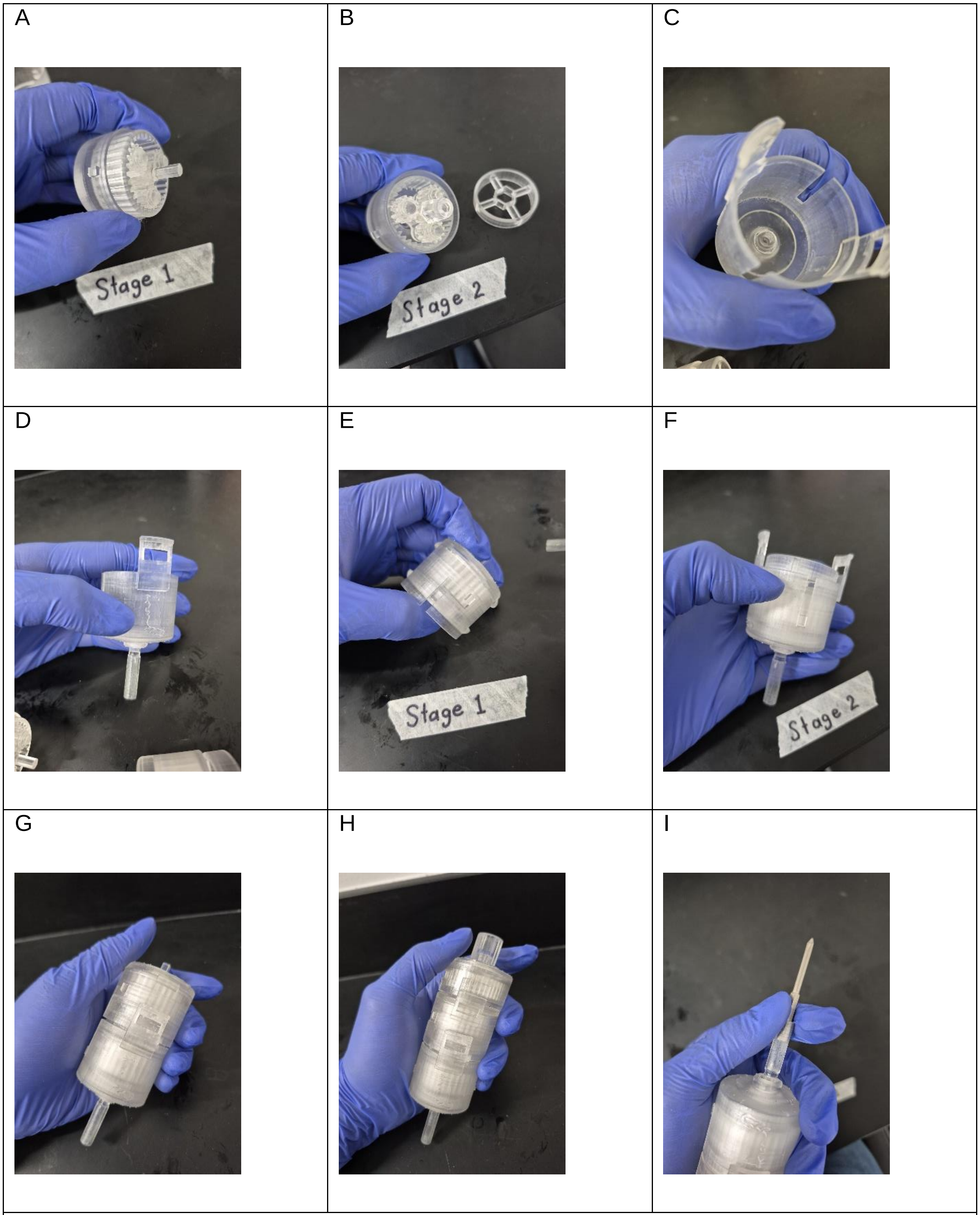

**Figure S1.4 | Assembling the Kepler driver. a, b.** Inserting the Stage 1 gear system into the top casing (**a**) and the Stage 2 gear into the bottom casing (**b**). **c, d.** Inserting the bit adapter piece into the bottom casing shell (**c**). The bit adapter stays in place inside the bottom casing shell if placed correctly (**d**) **e, f.** Inserting the Stage 1 (**e**) and Stage 2 (**f**) gear assemblies into their respective shells. **g.** Mating of the Stage 1 and Stage 2 gear assemblies by snapping their shells together. The latches on the side hold the two modules together. **h.** Mating the counter module to the rest of the Kepler driver. **i.** Fitting a 4 mm hex screwdriver bit into the bit adapter.

### Supplemental Note 2: Implant and saline bath assembly

Implant assembly can, in theory, be completed in as little as 4 hours. This is assuming that all parts have already been printed, washed, post-cured, and sanded, and that the probes have already been glued to the shuttle. The total assembly from printing to mounting can be completed over the course of 2 days otherwise. All implant pieces can be found in our GitHub repository (https://github.com/rjibanezalcala/EXPLORE).

#### Drilling

On the following implant components, drill a hole into the areas below (**Fig. S2.1a**) using the indicated drill bit size:

- **Skull interface**
  - 1.0 mm: The holes in the middle of each of the 3 recessed hexagonal areas.
- **Implant body**
  - 1.0 mm: The top bilateral sockets.
  - 1.0 mm: The bottom bilateral sockets
  - 1.0 mm: The bottom three screw holes
  - 1.4 mm: The top drive screw hole.
- **Probe shuttle**
  - 1.4 mm: The drive screw hole
- **Headstage interface**
  - 1.0 mm: The bilateral screw holes on the bottom
  - 1.0 mm: The single screw hole in the hex nut indent.
  - (optional) 7/64 in: The bilateral holes above the screw holes
- **Cap**
  - 1.0 mm: The single screw hole
- **Surgery stereotactic holder**
  - 7/64 in: The two holes on either side of the holder.
- **Shuttle stereotactic holder**
  - 1.4 mm: The drive screw hole

#### Threaded inserts and heat-setting

Heat-setting should happen as soon as possible between washing the 3D prints in IPA to remove excess uncured resin and the post-cure. This ensures malleability of the plastic and prevents cracking. A soldering iron set to about 200ºC seems to work well if the inserts are pushed in a slow, controlled manner.

- **Probe shuttle**
  - Heat-set an M1.4 threaded insert into the only socket on the shuttle from the top. The insert should be flush with the body of the shuttle (**Fig. S2.1b**).
- **Skull interface**
  - Heat-set or glue an M1 hex nut into each of the 3 recessed hexagonal areas (**Fig. S2.1c**). You may apply a thin layer of UV resin over the hex nuts to secure them in place if needed.
- **Implant body**
  - Heat-set an M1 threaded insert onto each of the 2 sockets located on the top side of the implant body (**Fig. S2.1d**). Take extra care to do so slowly to minimise cracking of the plastic. If cracking does happen, fill the area in with UV resin.
- **Heastage dock**
  - Heat-set one **M1.4 thread** into each of the 2 pre-drilled holes on the **headstage interface**, taking care to not push the insert all the way to the other side (**Fig. S2.1e**).

*IMPORTANT: Make sure that threaded inserts are well bonded to the plastic and do not turn when a screw is screwed into them!*

***(Figure S2.1 on next page)***

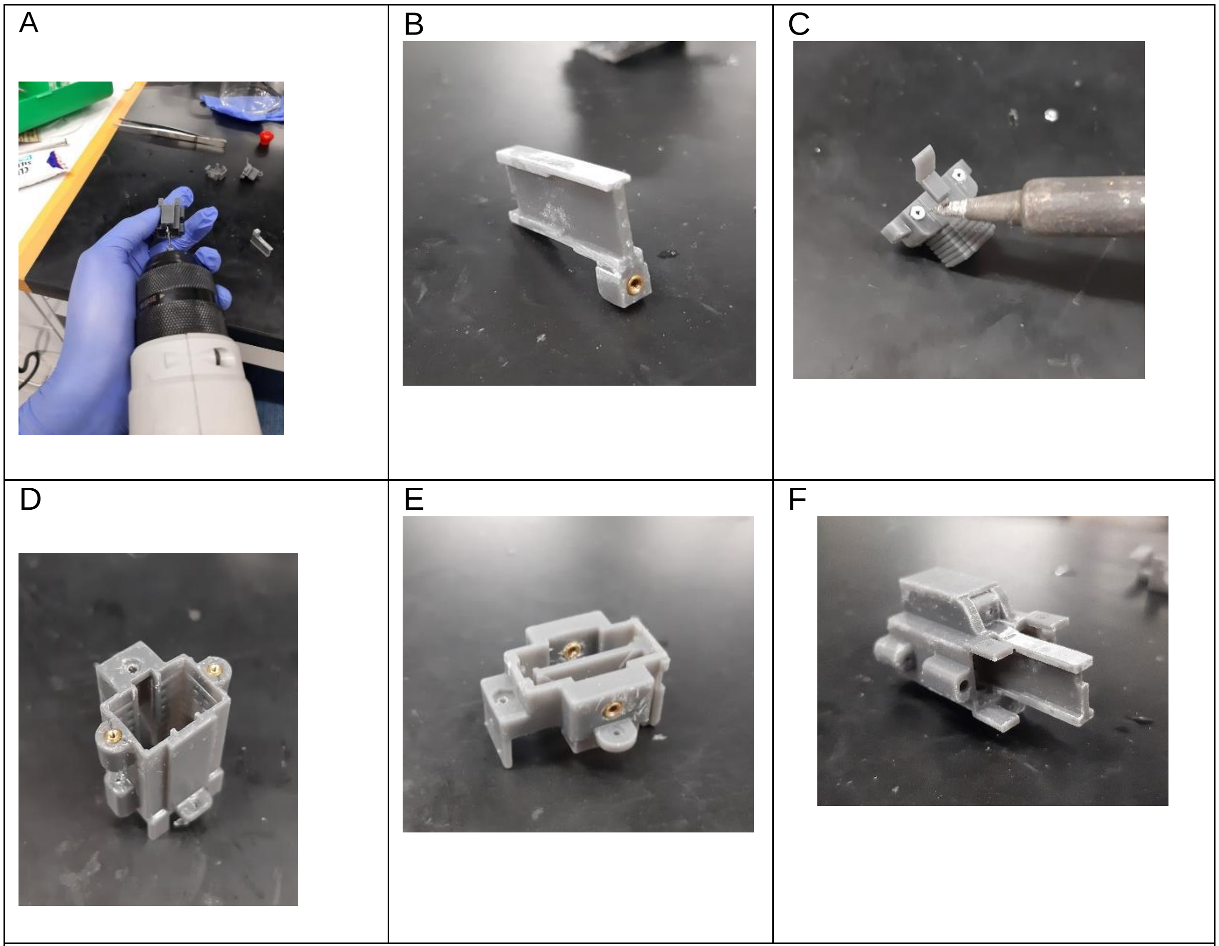

**Figure S2.1 | Drilling and threaded inserts. a.** Drilling of implant pieces to ensure that screws, inserts, and wires fit in their respective receptacles. **b.** Shuttle with M1.4 threaded insert. **c.** Heat-setting three M1 nuts onto the skull interface, two pictured, one on the back. **d.** Two M1 inserts on the top of the implant body. **e.** Two M1.4 threaded inserts on the headstage interface. **f.** Fitting the shuttle into the implant body. The shuttle may need to be sanded with sandpaper in order for it to fit, however some amount of resistance to movement is desirable.

#### Sanding

The only piece which may need some amount of sanding with sandpaper is the shuttle. Sand the edges of this piece if it does not fit into the implant body (**Fig. S2.1f**). However, make sure that the shuttle still moves inside the implant with some resistance, as this will help stabilise the probes to prevent undesired movement and drift.

#### The grounding circuit

1. Use the **1.6 mm drill bit** to drill two holes into the **copper sheet**. Cut two 4 mm x 4 mm sections around the holes and sand the edges down to create two small, round **copper plates** no more than 4 mm in diameter (**Fig. S2.2a, b**). You may

use a hammer to flatten the segments if needed. Sand one side the **copper plates** to remove any coating the metal may have. Set these aside.

2. Remove the black insulating plastic around two pin connectors using wire cutters or pliers. The pin should have a long prong on the flat side, and a shorter prong on the rounded side. Set the two bare metallic pin connectors aside for now.
3. Cut four 6.5 cm segments of the **PFA-coated wire**. Use **sandpaper** to scrape off about 1 cm of the wire's coating on each end. The wire cannot be soldered with the coating on. Set two segments aside.
4. Take two of the wire segments and carefully solder one end of each wire to the closed side of a socket connector (**Fig. S2.2c, d**). Be careful to not use too much solder as this may make fitting the socket connector into the implant difficult. *If the socket connector or wire do not hold the solder well, use flux!*
5. Use the soldering iron to tin both **copper plates** and solder one end of the remaining **wire** segments securely onto each of the plates (**Fig. S2.2e, f**). Take care to not solder over the hole you drilled. *If the copper or wire do not hold the solder well, use flux!* Finally, screw in a **bone screw** to test its fit, expand the holes using the drill if the screw does not go through. The screw should stay in place but be free to turn in place. Leave the screw in place for surgery.
6. Place a pin connector on each receptacle on the top of the **skull interface** rounded side first. Affix them using UV resin.
7. Solder the wire and copper plate assembly onto the pin connector from the bottom (**Fig. S2.2g**). Solder the wire parallel to the prong for a stronger bond. Use flux if you have difficulty soldering the two components together.
8. Thread each wire + socket connector assembly wire-first through the connector receptacles on either side of the bottom of the **implant body** and pull the wire through until the connector is inside the receptacle (**Fig. S2.2h**). If the pre-made channel is not wide enough, use the **1 mm drill bit** to open it.
9. With the help of **tweezers**, thread the wire through the pre-drilled holes above the receptacles so that the wire comes out through the top of the casing from the inside (**Fig. S2.2i**). Use **UV resin** to cover the hole and the wire. This will affix the **wire and socket connector** in place and protect the inside of the implant. Tape down the excess wire on the top to the side of the **implant body** to move it out of the way when loading the **probes** (**Fig. S2.2j**).

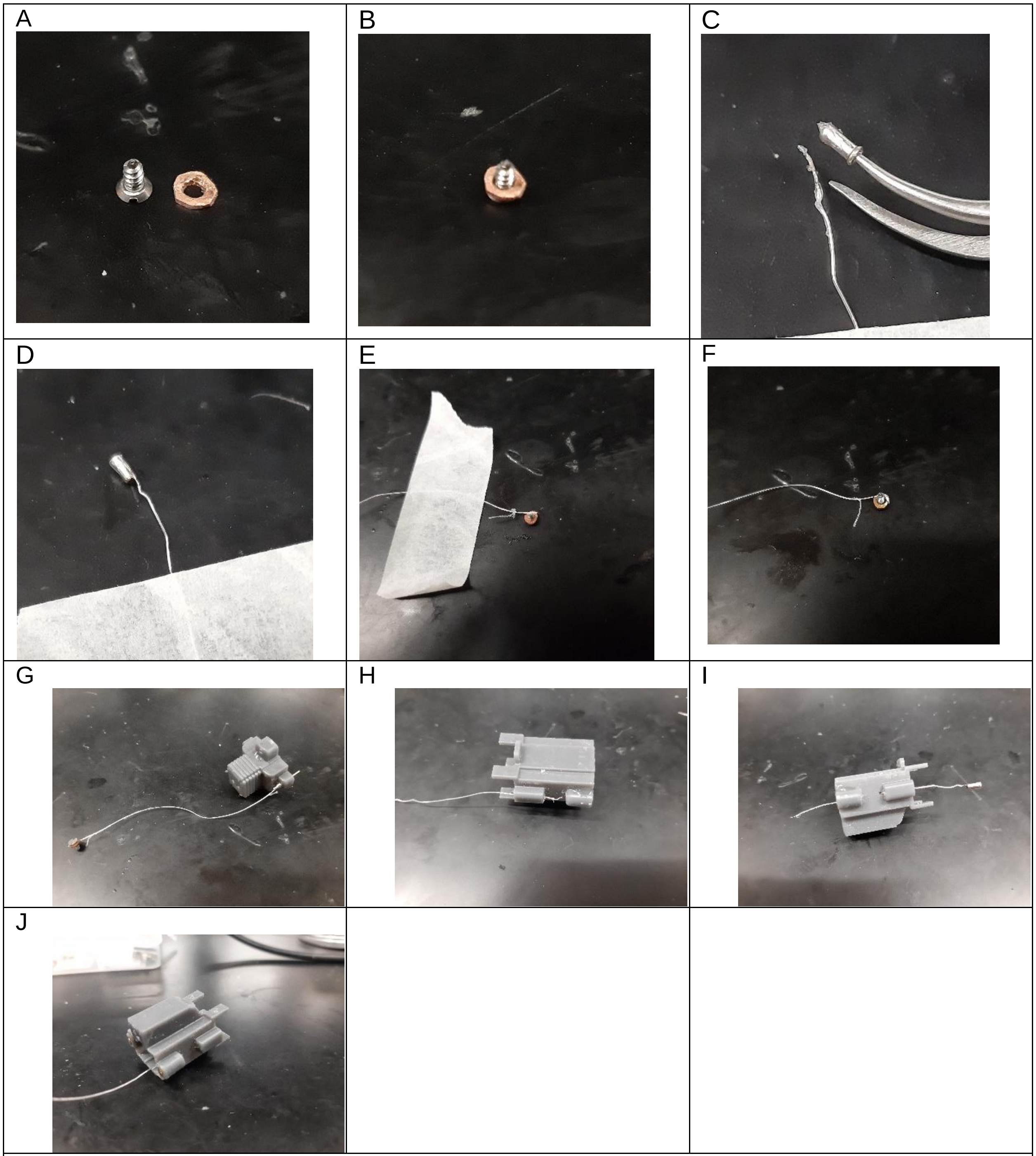

**Figure S2.2 | Grounding circuit components. a.** 4 mm diameter copper plate next to a skull screw. **b.** Test fitting the skull screw with the copper plate. The skull screw should fit through the plate and be able to rotate freely. **c, d.** Soldering wire to socket connector. The wire is soldered parallel to the side of the connector to increase strength of the joint. **e, f.** Soldering the wire to the copper plate, using tape to hold the wire down. **g.** Soldering the copper plate and wire to a pin connector inserted into the skull screw. **h.** Threading the socket connector and wire though the bottom pin receptacle on the implant body. **i.** Threading wire through the side pinhole on the implant body. **j.** Moving the wire to the side of the implant body, so it doesn't interfere with assembly of the implant.

#### Post-curing

Post-curing parameters will vary depending on your resin specifications. For Formlabs *Tough 2000* resin, we used a *Form Cure* at 60ºC for 60 minutes, as per manufacturer's recommendations. Please check your resin post-curing specifications.

#### Probe preparation

We use silicone glue to affix the probes onto the probe shuttle because it is strong and is water resistant, in addition to being flexible enough to be removed if the probes need to be removed from the shuttle for any reason. Silicone glue starts curing fairly quickly! Make sure to work quickly during this step as partially-cured glue will not hold as well.

*IMPORTANT: The probes will be exposed to damage at this stage. Make sure to take all possible precautions to avoid breaking the probe shanks. Wear gloves and use anti-electrostatic discharge equipment to prevent electrical damage to the probes.*

1. Apply **silicone glue** to the **probe shuttle** on one side. Carefully take one probe and press it against the glue while gently sliding it forward so that the probe catches onto the front stops on the shuttle. Carefully flip the shuttle over and repeat this with a second probe (**Fig. S2.3a**).
2. Place the shuttle and probes back into the box that the probes came in, held in between two of the foam pads. Set aside and <u>allow the glue to cure for 24 hours</u>.

#### Mounting the probes onto the implant

1. Mount the **stereotactic shuttle holder** onto one of the stereotactic arms then raise the DV as far up as possible. Place a **M1.4 x 4** screw into the screw hole.
2. Mount the **skull interface** onto the **holding bay**. Place the **holding bay** down beneath the stereotactic device arm. Secure the **holding bay** in place with tape or a **M6 x 15 screw** if appropriate (**Fig. S2.3b**).
3. Carefully take the **shuttle with the probes attached** out of the box and slide them into the holder.
4. Using a **screwdriver**, turn the **M1.4 x 4** screw <u>clockwise</u> until it catches and pulls the shuttle fully into the holder.

   *WARNING: The probes will be extremely vulnerable to damage at this point. Take care to not manoeuvre anything underneath the probes or you may risk breaking the probes.*

5. Carefully manoeuvre the shuttle around and into the skull interface; when lowering it, monitor the probe shank using the surgical microscope.
6. Lower the **implant body** onto the assembly carefully; first turn the casing slightly so that the probe's ribbon FPC connectors fit all the way through, then turn it again so that the drive screw receptacle lines up with the drive screw cover on the skull interface. Press the down until the screw holes line up with the **M1 nut** threads.
7. Screw in all three mating points between the **skull interface** and **shuttle casing** with **M1x4** screws, starting by the vertical one (**Fig. S2.3c**).
8. Short the Ground and Reference pads on the probes with wire, then solder one of the **ground wires** to the pads to complete the ground circuit; repeat on the other probe. Do not solder the two probes together, each probe should have its own ground.
9. Insert the drive screw into the drive screw receptacle on the **shuttle casing** and turn it clockwise to retract the probe. Fold the FPC ribbon and ground wire into the implant, letting the connector stick out from the top.
10. Lower the **headstage interface** onto the FPC connectors so that they thread through the slots. Have the connectors stick out about 5 mm from the top of the **interface** and drop the two **ribbon pinchers** into the headstage interface. Screw in a **M1.4x4 screw** on either side to pinch the connectors and hold them onto the middle spacer on the interface. Take care not to screw them in too much as the plastic may break.
11. When every piece of the implant is securely fastened in place, take the implant off the **holding block** and slightly lower the probe by turning the drive screw counterclockwise. Inspect the shanks under light to verify their integrity (**Fig. S2.3d, e**).
12. Place the **headstage interface cap** on if desired, however the cap must be removed prior to implantation in order to mount the implant onto the **surgery stereotactic holder** and to access the drive screw.
13. Keep the built implant on the **holding bay** until implantation.
14. In preparation for the surgery, mount the implant onto the **surgery stereotactic holder** (**Fig. S2.3f**) with the probes fully retracted for protection.

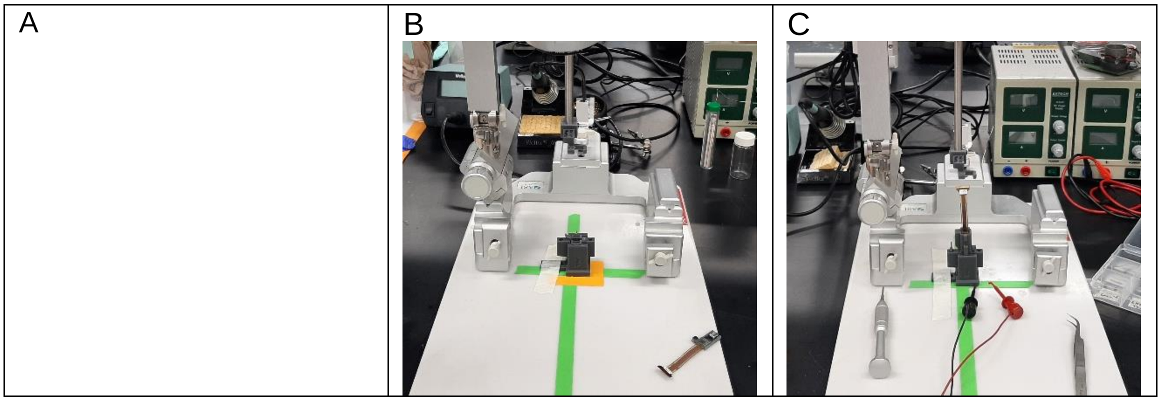

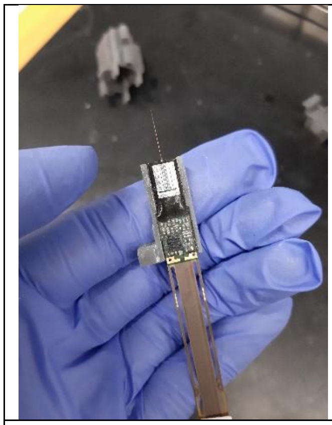

D

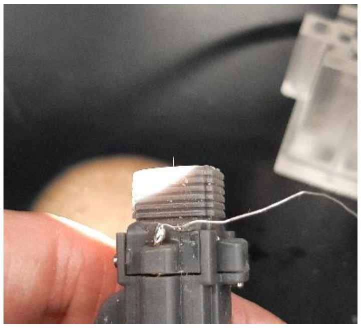

E

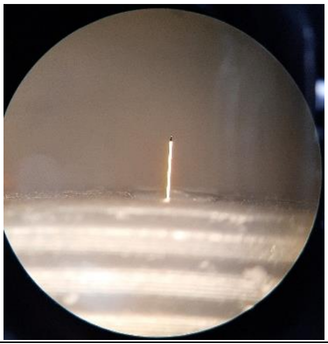

F

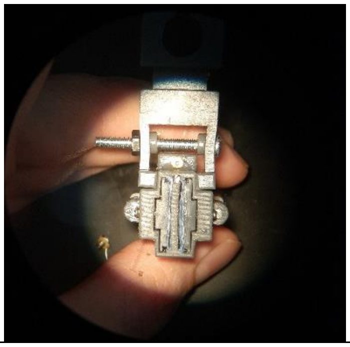

**Figure S2.3 | Probe mounting. a.** Glueing the probe to the shuttle piece with silicon glue, typically takes at least 12 hours to dry enough to continue. **b.** Preparing for the mounting procedure using a stereotaxic device. The shuttle and probe are placed in the shuttle holder and the skull interface is placed on the holding block below it. The shuttle and probe are then maneuvered into place while monitoring the probe's shank with a surgical microscope. **c.** After the probe has been mounted, the rest of the implant is assembled on the holding block to protect the probe shank. **d, e.** The probe is lowered slightly to verify the integrity of the shank under the surgical microscope. **f.** The implant is *firmly* mounted onto the implant holder in preparation for surgery.

#### Assembling the saline bath

Assembling the saline bath is as simple as cutting two copper plates (10 mm x 19.5 mm), running a wire through each tube from the front, then soldering the wires to the plates. The length of the wire can be 20 - 30 cm or more, if desired. We crimped one banana connector to each wire end to ensure compatibility with the A.M.P.I. Iso-flex Electrical stimulus isolator. We recommend affixing the copper plates with silicone sealant and letting it dry for at least 24 hours to ensure a watertight seal.

### Supplemental Note 3: Surgery protocol

#### Section 1: Surgical procedures

**Step 1: Surgical Prep**

1. Construct custom rat cage (**see 'Section 3: Custom extra-tall cage'**)
2. Sterilize tools before surgery
3. Sterilize implant with alcohol, then dip in saline
4. Follow PPE guidelines
5. Check and fill isoflurane and oxygen tank
6. Weight charcoal tanks, document weight:
7. Turn on heating element: heating pad
8. Weigh the animal

**Step 2: Anaesthesia / Shaving**

1. Administer Ketamine Cocktail (ketamine (100 mg/kg) – xylazine (10 mg/kg) solution) through an Intraperitoneal (IP) injection. Dosage: *0.1 ml/ 100 gm* or *0.05 ml/100 gm*
    a) Conduct a toe pinch to ensure that the animal is fully anesthetized
2. Write down the total amount administered including boosts. Boosts should be administered intramuscular and should be ≤ 0.01 ml.
3. Anaesthesia: induction: 2%, maintenance: 1-1.5%, oxygen flow: 2-2.5 L/min
4. Put PURALUBE ointment on eyes to lubricate and protect them. Do this continuously throughout the surgery.

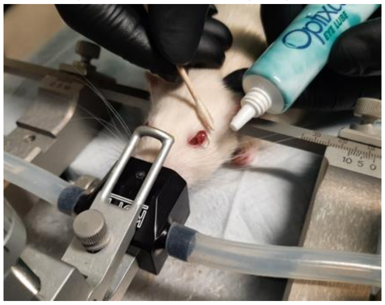

5. Shave scalp, from ears to near the nose *or* use Nair hair remover cream to remove hair from the desired area

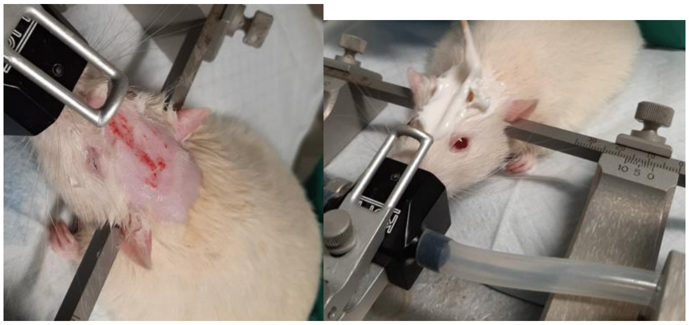

6. Secure animal in stereotaxic frame.

) Make sure the animal can say “yes” but not “no” ensuring that the skull does not move.

7. Insert the ear bars (<1mm difference) L=__________ R=__________
8. Clean surface with 70% alcohol, then iodine. 3X repeat
9. PURALUBE ointment on eyes AGAIN to prevent corneal drying
10. Use a thermometer to check the temperature of the rat before opening. Continue to check.

**Step 3: Begin surgery**

1. Inject 1ml of Bupivacaine subcutaneously around the incision site and massage it in

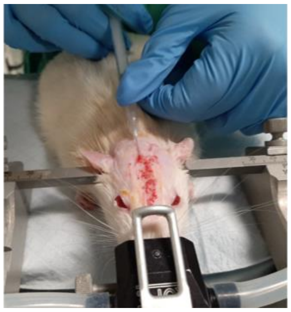

2. Expose cranium by making a midline anterior-posterior surgical incision, from the nasal bone to the end of skull with scalpel blade
3. Carefully push back and remove any connective tissue to fully expose the skull.

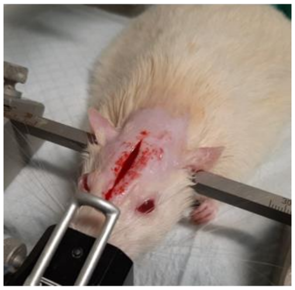

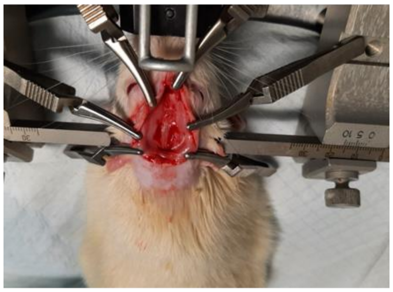

4. Remove any remaining connective tissue and with surgical bulldog clamps hold back the tissue

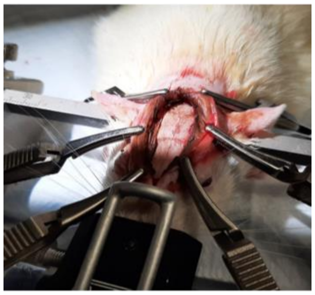

5. Use h2o2, styptic powder, or cold saline to clean and stop bleeding

**Step 4: Screws**

1. **FLAT SKULL**: Locate Bregma and Lambda and ensure < 0.5 mm difference between the two

| Bregma | Lambda |
| --- | --- |
| DV: | DV: |
| ML: | ML: |
| AP: | AP: |

2. Locate and mark the implant hole based on the coordinates below. Make a small indentation mark with a small ball-head 0.6mm drill bit once you locate where you will be implanting your probe.

| AP (-1.2mm from Bregma): | ML (+2.5mm from Bregma): |
| --- | --- |

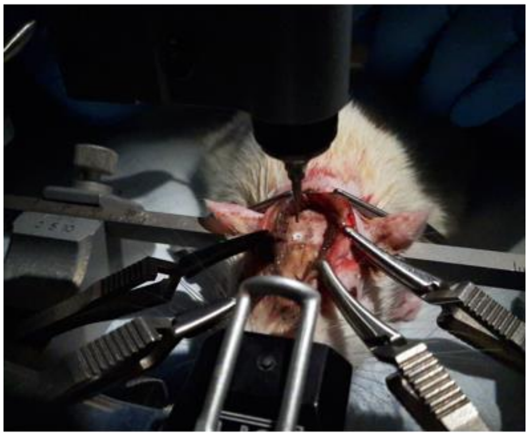

3. With a 1.5mm trephine micro drill bit go back to your target coordinates and drill a mark over your previous mark from Step 2.

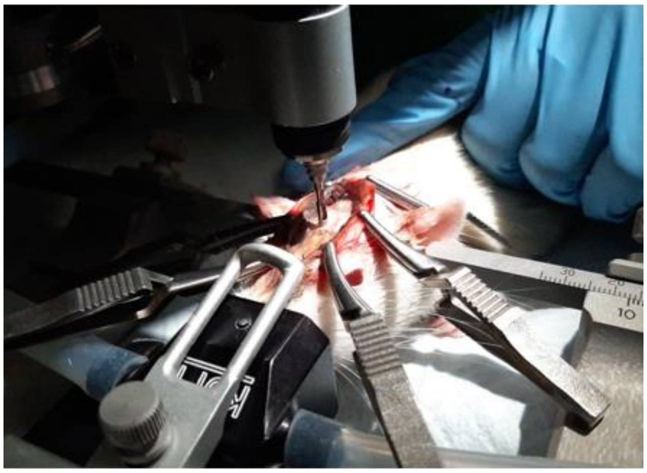

4. Temporarily lower the probe housing and place above the marked divot to ensure the screws are outside the area the housing will take up. You can mark the border around the probe housing with a surgical marker.

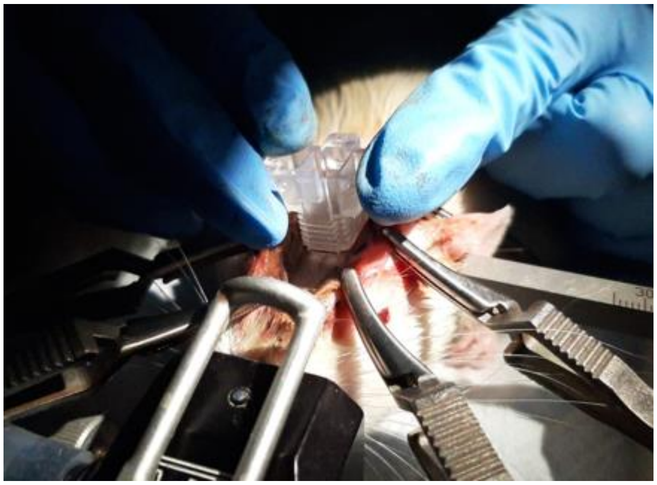

5. Plan where you will place your screws. NOTE: you will need 6-10 total screws. 1 screw will serve as the ground screw. This screw cannot be touching any other screws nor be near muscle or tissue as to reduce or avoid electromyographic interference during recordings.

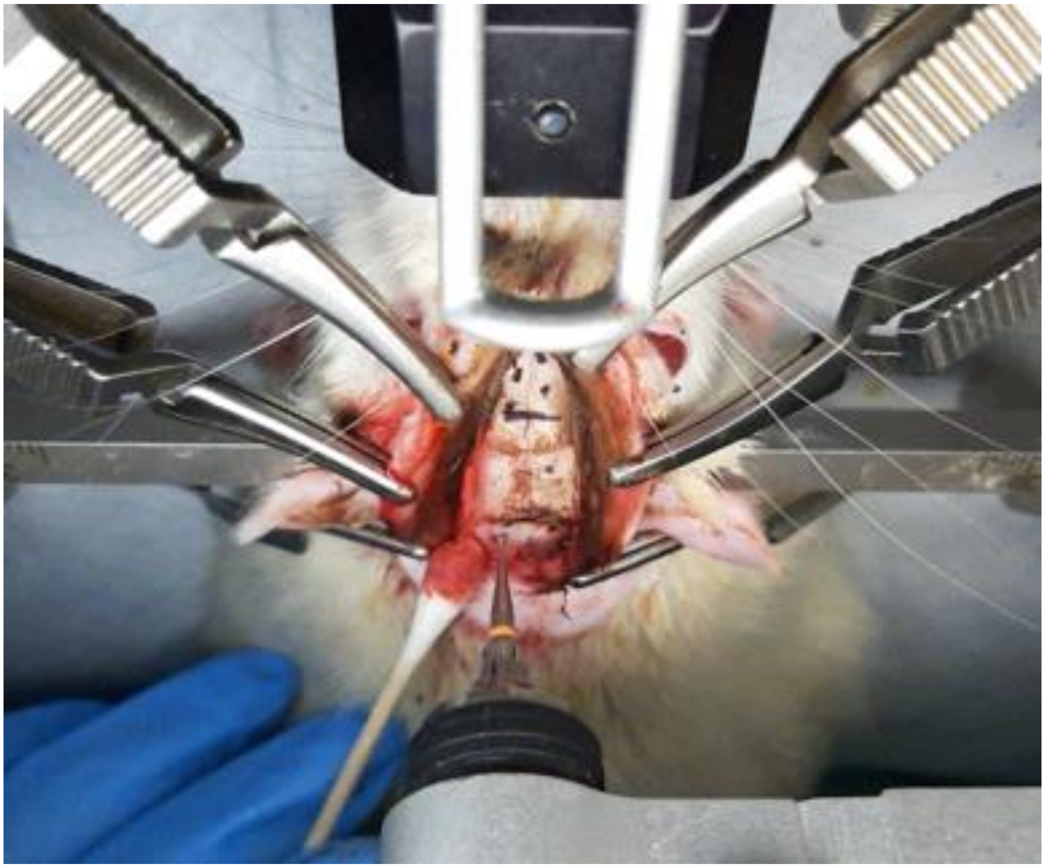

6. After you mark where you will be placing the screws, using a 0.6mm drill bit, make small holes that are deep enough for the screws to hold onto the bone but not deep enough to fully penetrate bone or dura mater. Once you have your hole, insert the screws using a small screwdriver and some pressure. Make sure to not screw in fully (as long as they are tightened and secured).

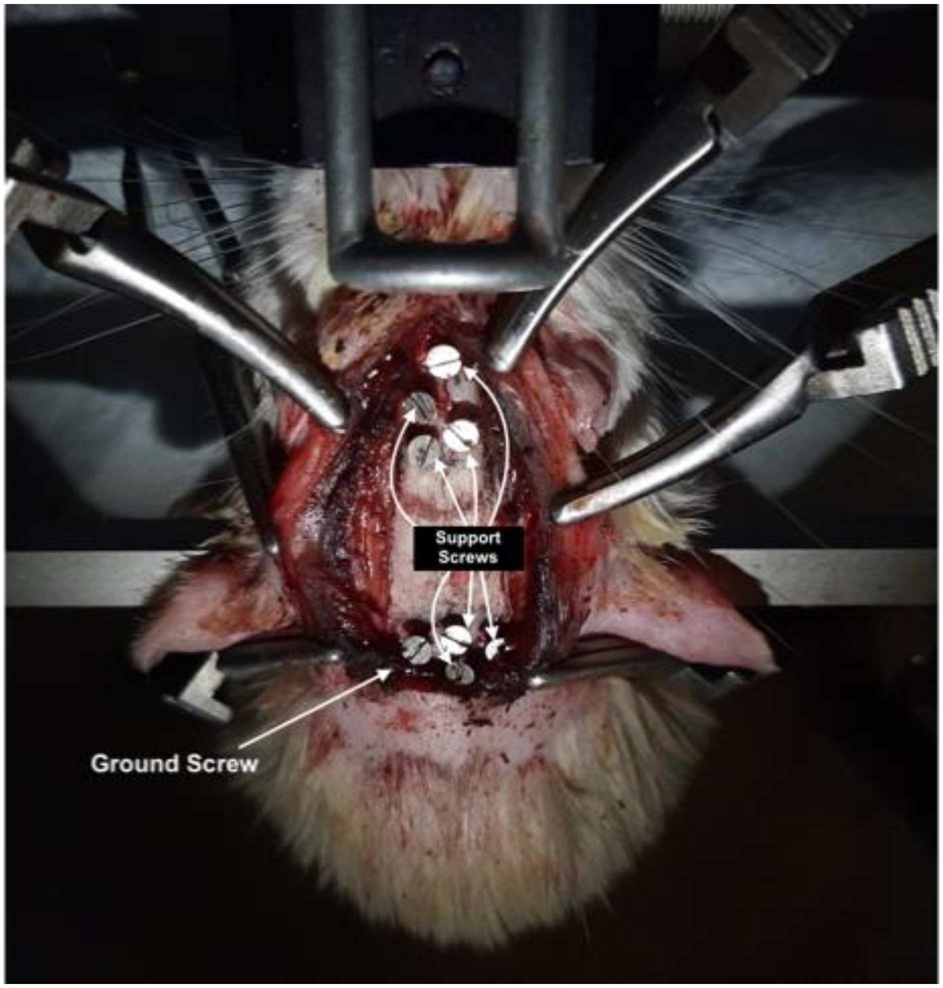

**Step 5: Probe implant**

1. Based on the Bregma, locate your coordinates

ML bregma ______ +2.5 =_______// AP ______ -1.2 = ________

   a) Locate Bregma, then move to coordinates.
   b) Mark the skull, then lift drill bit to ensure the smaller hole is in the middle of the marked craniotomy hole
2. With the 1.5mm trephine drill bit, carefully drill until the bone is translucent and barely attached on one side.
3. With small surgical tweezers carefully remove the bone. Save the piece to measure later.
4. Carefully remove any remaining dura mater using precision tweezers under microscope

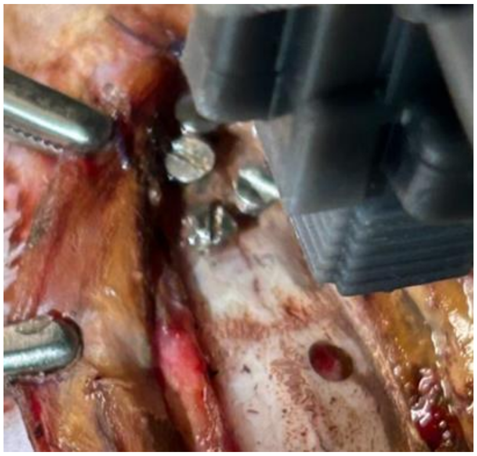

5. Inside of the craniotomy drop around 1-2 drops of 4 µg/mL dexamethasone is dropped into the top of the trephined hole, allowing for temporary softening of the brain, and reducing risks of inflammation

**Step 6: Probe**

1. Secure implant on the stereotactic arm and on the specialized implant holder
2. Locate your ground screw and screw the copper ground pad onto the skull. Careful to not break the wire connecting the ground pad to the implant

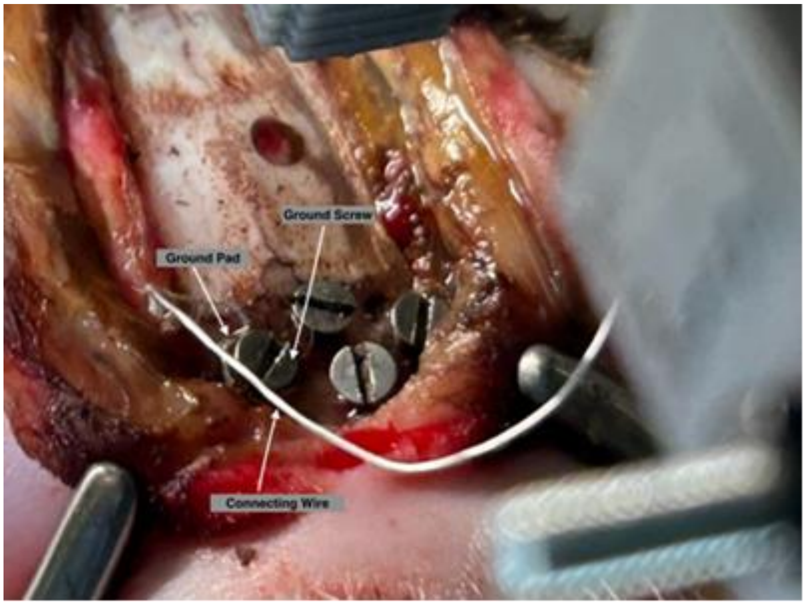

3. With the screwdriver make ~17 turns to bring out the probe approximately 5 mm
4. Locate bregma, then move to target coordinates

| AP (-1.2mm from Bregma): | ML (+2.5mm from Bregma): |
|---|---|

5. Measure the skull thickness by measuring the bone extracted from the trephine hole in **Step 5** with callipers. Take note of it.
    a) Although risky, this distance can also be calculated by subtracting the DV coordinate where the skull is located from the DV coordinate from where the surface of the brain is located. To do this, first expose the probe shanks to some arbitrary amount (f.e. 8 mm).
    b) Touch the probe shanks to a portion of the skull near the craniotomy while monitoring them with a surgical microscope. Take note of the DV coordinate on the stereotaxic device.
    c) Place the implant over the hole and zero at the Dura mater coordinate. Lower the implant until the shanks touch the brain. Take note of this DV.
    d) Subtract the DV from where the shanks touched the skull from the DV where the shanks touched the brain inside the craniotomy. This will be the skull thickness.
    e) Fully retract the probes again.
6. Place the implant over the hole and zero at the Dura mater coordinate
7. To ensure that the tip of the probe is implanted at a KNOWN depth, first expose the probe shank X mm by turning the drive screw <u>counter-clockwise</u> Y times (see formulae below or consult the [surgical log excel sheet calculator]).
    a. $X = (Skull\ thickness + padding) + DVinit$ ,
    Where $DV_{init}$ is the desired <u>initial</u> implantation depth or *DV* (can be anywhere between 1.4 and 2.0 mm, depending on the brain region which is being targeted), and *padding* is the thickness of the bottom wall of the implant's skull interface. Typically, *padding = 0.5 mm*.
    b. $Y = X / (Drive\ screw\ pitch\ in\ mm)$.
    Drive screw pitch will likely be 0.3 mm if the materials in **Supplemental Note 4** are used.

| DV(-3.5 from Dura) |
| --- |

*WARNING: The probe shanks will be exposed to damage at this point. If you need to manoeuvre under the implant, do so with extreme care!*

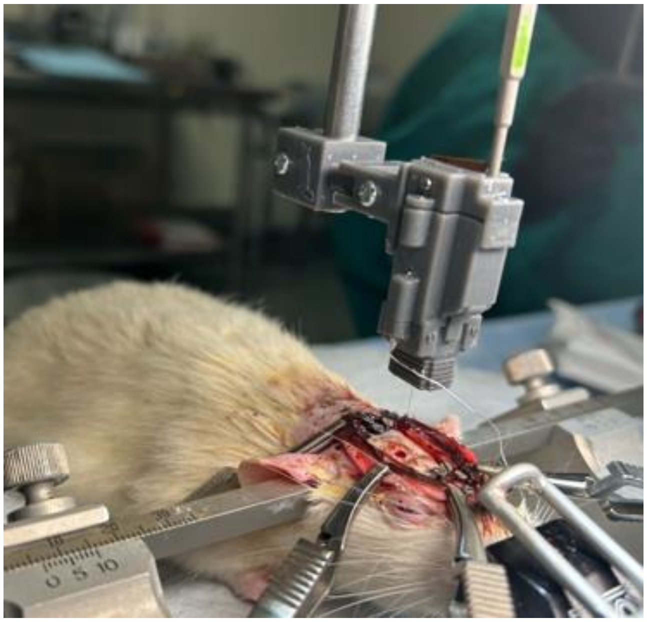

8. Add more dexamethasone and with a point swab or rolled up sterile paper towel bit, remove any excess liquid or blood
9. Make sure there is no dried blood or remaining dura and add more dexamethasone.
10. Using a microscope, make sure you can see both the tip of the probe and the surface of the brain.
11. Slowly lower the probe into the brain, but before you fully implant, stop at the top of the brain tissue and “dance” or “bounce” on it with the probe
    a) You are making sure that when you are dancing the probe is not bending. IF it does bend add more dexamethasone to irrigate the brain tissue.
    b) Once you can see the fragile probe is not bending, slowly lower it until the implant housing lays flat on the skull.
    c) Go back up, check the probe is still intact. If it is, repeat the “dancing” and irrigation steps and fully lower.

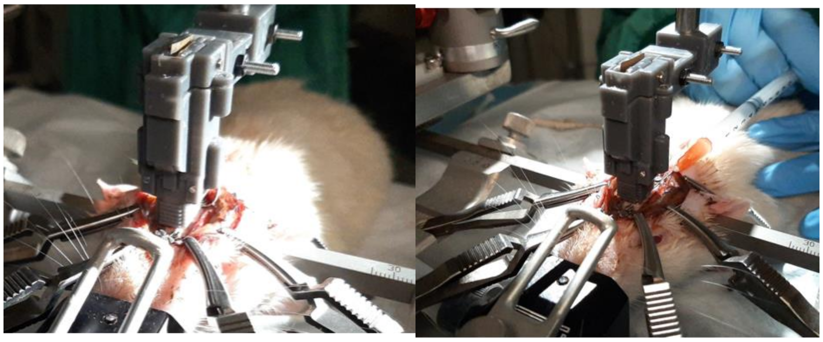

12. Once the implant housing is flat on the skull again, using a syringe with a 25” needle carefully put liquified sterile Vaseline into the probe area on top of the skull and within the trephine hole to prevent the buildup of blood or other bodily fluids or tissues onto the probe

13. Wipe the excess around the exposed skull area with a cotton swab
14. Use 1-3 sticks of silver nitrate (activate it using saline) to put around the muscle tissue and the skull to limit re-growth.

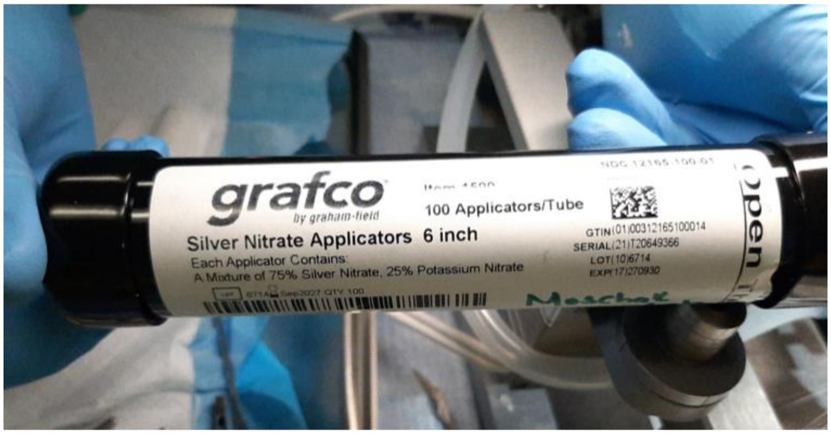

**Step 7: Cement Metabond**

1. Mix liquid-Metabond (5 drops blue liquid + 2 drops of Gold catalyst) with solid-Metabond (2 scoops of metabond powder)
2. Make sure the skull is clean and mostly dry and proceed to fully cover the skull with a thin layer of liquid-metabond. Let it dry.

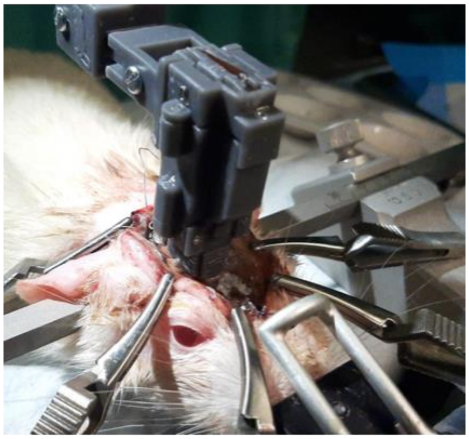

AVOID GETTING NEAR THE HOLES OR UNDER THE IMPLANT

3. Use the metabond mixture to build a layer on top of the skull
4. Let the cement fully dry (While waiting, cut their nails)

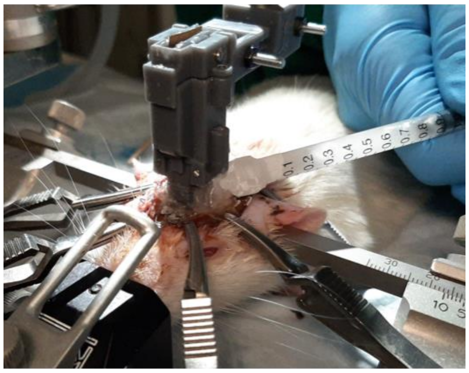

5. Dental Cement: Add a layer of cement above the metabond and layer by layer build a headcap around the implant, making sure that your layers dry before adding more cement.
6. Make sure the entire region is covered in cement (screws, any exposed skull, and the bottom ⅓- ⅔ of the implant housing). Be sure no hair is caught in the cement, and that no cement falls into the probe area.
7. When the cement hardens, loosen the screw and nut that hold the implant onto the holder and slide it off the implant by raising the stereotactic arm.

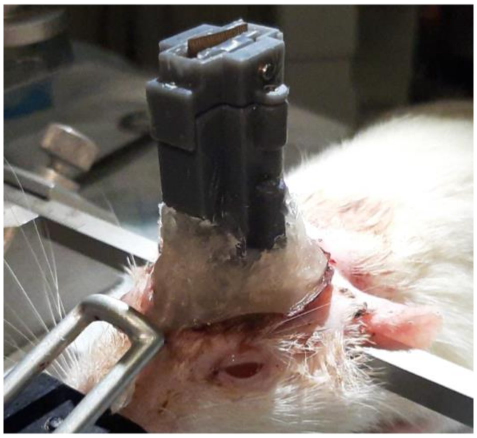 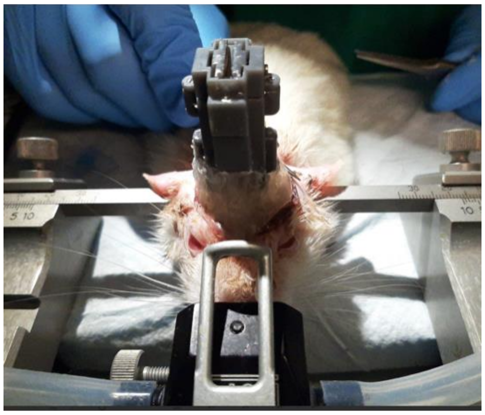

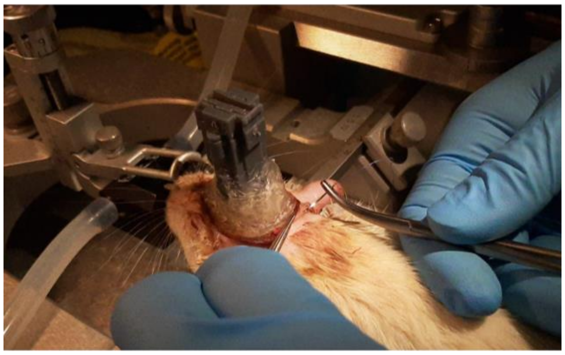

8. Once your cement layers have fully dried, carefully remove the implant holder
9. If there is excess skin do 2-3 sutures in the back over the cement headcap that you build using 5-0 silk sutures.
10. Check temperature = ____________ F

**Step 8: Closing up**

1. Lower the anaesthesia levels Oxygen= _________, Iso =_________
2. Rinse area with antibiotic lavage and chlorhexidine solution

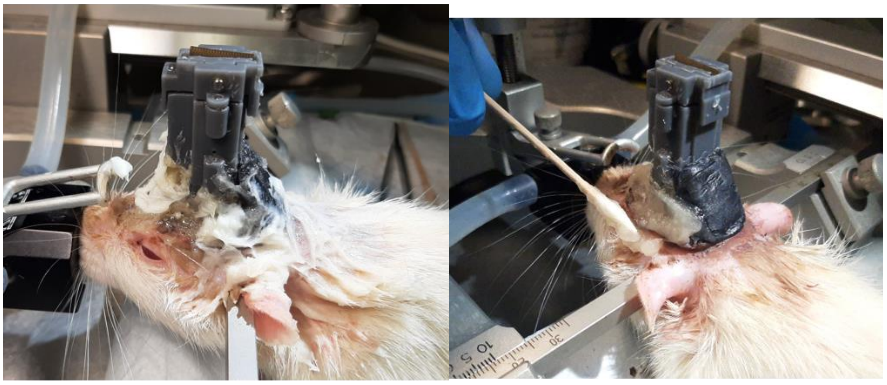

3. Add a mixture of triple antibiotic (neomycin, polymyxin B, and bacitracin) and chlorohexidine creams around the area and under the nails.

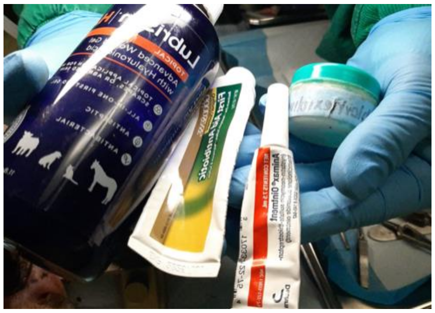

**Step 9: Immediate post-op care and probe adjustment**

- Inject Antibiotic and Meloxicam in 10ml of Ringer solution.
  - Inject the solution warmed (5ml on each side, **see Table S3.1 for dosing**)
  - Inject Rx for 3 additional days post-op and one extra day of antibiotic only in 10ml of Ringer (**see 'Section 4: Post-op care guidelines'** for additional first-week post-op care).

| | | | |
|---|---|---|---|
| Enrofloxacin (Antibiotic) | **Antibiotic**<br>Dose<br>5mg = x mg<br>1 kg kg | **Antibiotic**<br>Concentration<br>22.7 mg = mg<br>1 ml x ml | **Antibiotic**<br>Final volume: |
| Meloxicam (NSAID) | **Meloxicam**<br>Dose<br>1mg = x mg<br>1 kg kg | **Meloxicam**<br>Concentration<br>5 mg = mg<br>1 ml x ml | **Meloxicam**<br>Final volume: |

**Table S3.1 | Antibiotic and NSAID injection dosing.** Enrofloxacin and Meloxicam dosing chart to mix in with 10 ml Ringer.

- Using the *Kepler screwdriver*, lower the probe 100 - 200 µm (see surgical log excel sheet calculator).

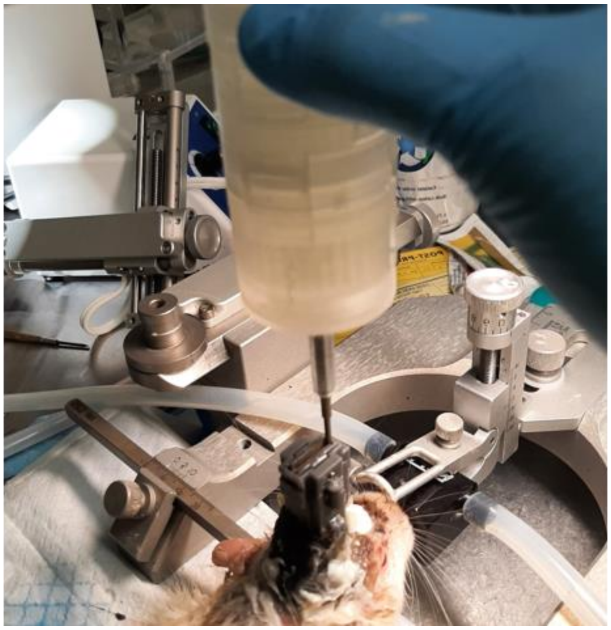

- Apply the mixture of creams (triple antibiotic: neomycin, polymyxin B, and bacitracin; and chlorohexidine creams) to promote healing and lower the risk of infection.
- Remove from anaesthesia. Time: __________
- Clean any blood from animal using saline
- Place in a new cage with surgical bedding, half on top of a heating pad
- Wait for the righting reflex, annotate the time the animal wakes up.
- Fill out post-op yellow card

#### Section 2: Potential complications and mitigation strategies for Neuropixels implant surgery

Damage to the Neuropixels probe during surgery can halt the procedure and compromise data integrity. To mitigate this, it is crucial to have a second implant prepared and ready for immediate use to avoid delays in cases where the probe shank fully breaks off. Probe damage can be minimized by ensuring it is fully retracted during steps where the probe is not in use, such as when screwing in the ground pad and ground screw. Retracting the probe into the implant during these steps protects it from accidental stress or impact. Proper handling and careful attention to each step of the surgery significantly reduces the risk of probe breakage.

If the connecting wire breaks while securing the ground pad, ensure soldering equipment, including a soldering iron, solder, and flux are available to repair the connection. Should this occur, the implant must be removed; this step must always be performed with the probe fully retracted to avoid further damage and before inserting the probe into the craniotomy hole. To further prevent the wire from breaking, be sure the wire is appropriately sized—around 50 mm in length—to allow manoeuvrability while minimizing the risk of tangling or breakage. The wire should be soldered parallel to the bottom of the ground pin on the skull connector for best stability. Adjust the implant's positioning using the stereotactic device to maintain alignment and stability during this process.

To prevent the implant from detaching post-surgery, ensure all connective tissue is meticulously removed during the preparation phase. Use silver nitrate sticks during surgery to prevent tissue re-growth underneath and around the implant. Additionally, applying silver nitrate around the cement 1–2 days post-op can further secure the implant if deemed necessary. This comprehensive approach ensures a stable and long-lasting implant, reducing the likelihood of complications.

Neuropixels implant surgeries present specific challenges that require careful preparation and real-time monitoring to ensure animal welfare and procedural success. Another critical complication is respiratory distress, including the cessation of breathing during surgery. To mitigate this risk, it is essential to maintain accurate ketamine dosing and to regulate isoflurane (iso) anaesthesia carefully. Isoflurane levels should be kept steady and low, not exceeding 2%, with oxygen flow consistently set at 0.5 L/min higher than the iso level. Continuous monitoring of respiratory rate is imperative, with adjustments to iso and oxygen levels made accordingly. Should an animal exhibit signs of waking at 2% iso, a 0.1 ml intramuscular (IM) boost of ketamine can be administered, followed by a reduction in iso once sedation stabilizes.

In the event of apnoea, immediate action is required. The animal should be removed from the nose cone, and a cardiac massage should be initiated while providing oxygen support. If breathing does not resume, cardiopulmonary resuscitation (CPR) should be started, including chest compressions and manual resuscitation with an oxygen pump, ensuring visible lung expansion. If these measures are ineffective, administer atipamezole hydrochloride (5.0 mg/mL stock solution) intramuscularly (IM) in the leg, avoiding the vein in the medial region. The dose is determined by body weight and can be estimated using a mental shortcut: extract the hundreds digit of the weight in grams, multiply by two, and shift the decimal two places left (e.g., a 200 g animal: 2 × 2 = 4 → 0.04 mL). Alternatively, a precise calculation follows the equation Dose (mL) = 0.000184 × weight in grams, yielding 0.0368 mL for a 200 g animal, rounded to 0.04 mL. Both methods produce comparable results, with minor adjustments made as needed. If that does not work, as a final measure, humane euthanasia should be performed to minimize suffering.

**Potential surgery complications, summary ( ! )**

1. ( ! ) Rat stops breathing during surgery.
    a. Immediately remove the rat from the isoflurane and begin CPR with assisted hand pump. Place the nosecone of the CPR pump on the rat's mouth and nose ensuring a tight seal. Press the hand pump to administer air. Ensure the rat’s lungs can be seen inflating and deflating. Sometimes the air will enter the stomach – this means the rat is angled oddly. Ensure the rat is flat on their ventral side.
2. ( ! ) Rat starts waking up during surgery.
    a. Immediately cease any cutting/drilling. Check that the isoflurane line is connected properly, with adequate flow and dose being administered. If the rat wakes up fully, disconnect them from the bite bar before they damage their teeth. If the rat is barely waking up, hold the rat down and turn up the isoflurane briefly to administer a higher dose. Ideally, you should always keep an eye on both the rat’s physiological signs of anaesthesia AND the isoflurane/oxygen levels to prevent this.
3. ( ! ) Rat is bleeding during surgery.
    a. This is expected, but depending on the level and location of bleeding it could be cause for concern.
        i. Bleeding at the skin
            1. Apply slight pressure using gauze and cold saline
            2. If bleeding continues, use a small amount of Kwik Stop Styptic Powder
        ii. Bleeding at the skull
            1. Apply slight pressure using gauze and cold saline
            2. If bleeding continues, use a small amount of Kwik Stop Styptic Powder. Note styptic powder can only be used before any craniotomies have been made.
        iii. Bleeding at the trephine hole
            1. With a point swab or rolled up sterile paper towel bit, remove any excess liquid or blood
        iv. Bleeding at the edges of the implant
            1. Wet a point swab with cold saline and with light pressure clean around the edges
4. ( ! ) The implant bent.
    a. Inspect the shank visually using a microscope. If no damage is visible and time allows, connect the implant to the system and test for damage before resuming implantation.
    b. If the implant is broken, use the backup (*you should have a backup always*).

#### Section 3: Custom extra-tall cage

Due to the size of the implant and risk of impact with traditional vivarium cages, we custom built an extra-tall cage for Neuropixels rats using the lower halves of two of our traditional cages. The steps below describe how these cages were built:

1. Stack the two cage halves open-side-down and *temporarily* tape them together with duct tape to make handling easier.
2. Test-fit the resulting box onto the vivarium rack and mark where the vivarium rack mating ports will need to be with a marker. The cage half with the markings will be the top half of the cage.
3. Begin by drilling a pilot hole using any drill bit smaller than 7/8 inch (about 2.22 cm) to guide the cut. Then, use a Dremel™ 545 Diamond Coated Cut-Off Wheel (which has a 7/8-inch diameter) to expand the hole to its final size. Ensure all burrs and melted acrylic plastic are removed for a clean finish.
4. The final hole will match the widest part (7/8 inch) of the green Tecniplast Green Line GR900 SealSafe Plus airflow mating ports. Insert the mating ports into the openings at the marked positions (**Fig. 3.1a, top, green plastic and rubber grommets**), ensuring proper alignment with the vivarium rack (**Fig. S3.1a, bottom**).
5. To secure the mating ports, apply hot glue between the outer edge of the mating port (7/8 inch) and the inner edge of the newly cut hole, forming a bond in the space between them. Focus on keeping the glue on the outside of the cage to prevent the rat from accessing it while ensuring a strong seal. Once the glue has fully set, set the cage aside.
6. To give the animal access to water, opt for water bottles with a bent waterspout or bend the spouts yourself. The bend angle should be greater than 90º, but less than 180º to allow water to flow through (**Fig. S3.1b**).
    a. You may also omit bending the spouts; however, a bend can help with positioning the water bottles on the side of the modified cage in a way they don't protrude out as much.
    b. Place an extra rubber stopper along the length of each waterspout, 1-3 cm from the tip, to prevent the rat from getting stuck under the protruding metal tip, as this can be fatal. Set the bottles aside once prepared.
7. Using a drill/Dremel tool, make holes in the front side of the lower cage half (opposite side to where the mating ports are) to allow the water bottles to be inserted from the front of the cage (**Fig. S3.1c**). *Unless you have a reason for not doing so, water bottles need to be placed in the front of the cage for the cage to fit in the vivarium rack.*
    a. If the cage allows, you may insert the bottles' waterspouts through an existing opening instead (**Fig. S3.1d, arrow**).
8. Remove the tape from Step 1 and stack the two cage halves as they were in the previous steps. Tape the back edge (under the mating ports) of the two cage halves together to form a hinge (**Fig. S3.1e**).

⚠ **Ensure that the sticky side of the tape is not accessible from the inside of the cage as this may pose a health risk to the animal.**

9. Add tape to secure the water bottles from the outside of the cage (**as shown in Fig. S3.1c**), ensuring the rubber stoppers from **step 4b** are present to stop the spout from sliding into the cage too much. Only 1-3 cm of the spout should be inside the cage, adjust the rubber stopper to achieve this.

   ⚠ **Excessive protrusion is a risk for headcap arrestment and death of the rat.** *Excessive protrusion of waterspout is shown in* ***Fig. S3.1d****. Ensure the waterspout is receded enough to prevent the implant from becoming lodged between it and the cage wall – this is a potential failure point. We use rubber stoppers along the waterspout to prevent it from reaching deep into the cage.*

10. Tape the post-op and rat information card holders to the front of the cage.
11. □**Clean the cage** with soap and warm water to remove any acrylic dust from drilling.
12. Add fresh, sterile low-dust surgical bedding to the inside of the cage. Fill the front of the cage with veterinarian-approved rat chow pellets. Add a cup of diet gel (wet food) topped with sugar pellets for easy nutrition while the rat heals.

***(Figure S3.1 on next page)***

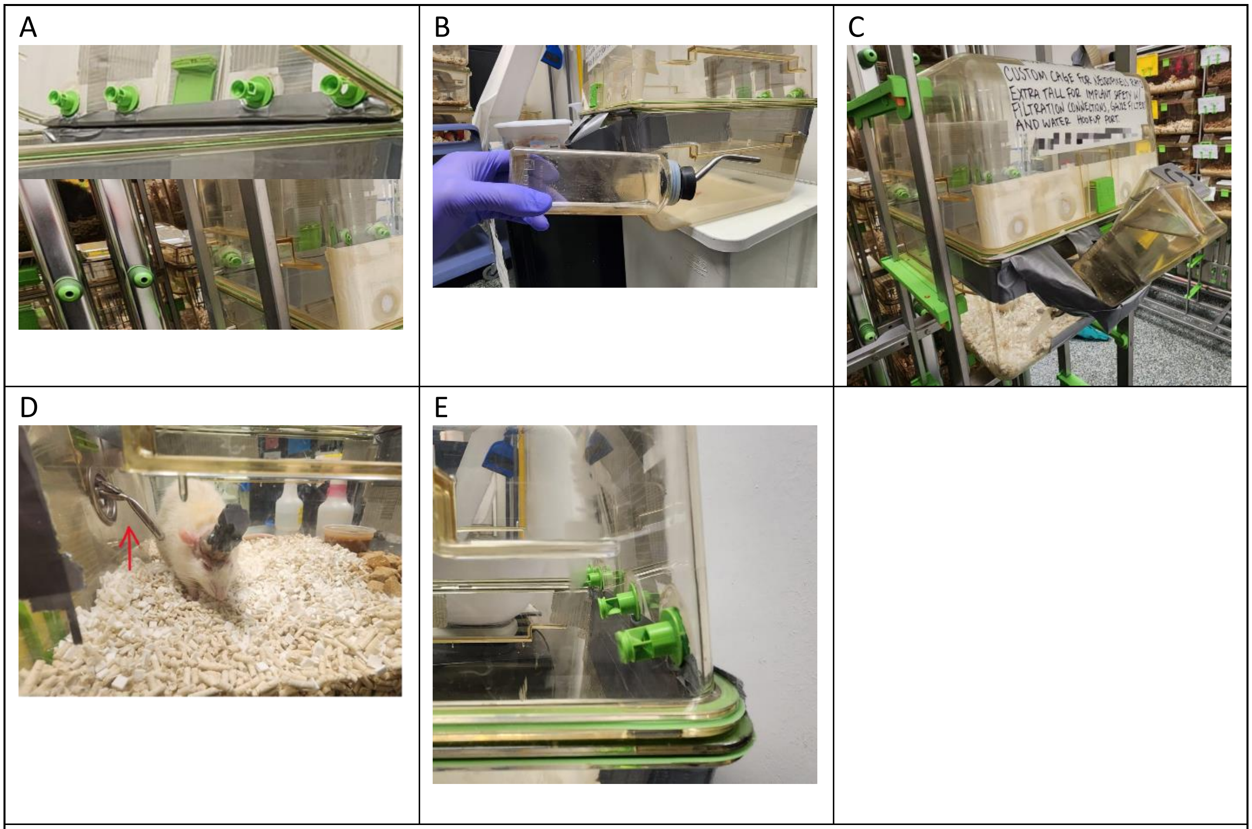

**Figure S3.1 | Custom home cage. a.** Vivarium rack mating ports on custom cage. **b.** Bending the water bottle spout to allow it to sit flusher with the outside cage wall. **c.** Water bottles are positioned on the front of the cage to allow the custom cage to fit on the rack. **d.** Water bottle spout introduced through a pre-existing entry hole on the cage. The figure shows too much of the spout inside the cage; ideally this length should be reduced to 1 – 3 cm by adding a rubber stopper along the length of the spout on the outside of the cage. **e.** Duct tape hinge on the back of the cage. The sticky side of the tape is covered from the inside. Mating ports also pictured.

### Section 4: Post-op care guidelines

##### 4.1 First-week post-op care

As a general rule of thumb, cage bedding should be changed every 1 to 2 weeks depending on cleanliness. Due to the custom nature of the cage, it cannot be autoclaved without destruction, thus it must be manually deep cleaned with soap and warm water every few weeks.

⚠ **For the list items below, it is recommended to anaesthetize the rat with vaporized isoflurane prior to post-op care due to the risk of movement and to minimize pain.**

- Routinely inject gently warmed 10mL ringer with antibiotic Enrofloxacin and NSAID meloxicam (**Table S3.1**) every day for 3 days after the surgery date. Bolus is spread across two subcutaneous injections (5mL, 5mL) in the flank/back of the rat to accommodate for the 10mLs.
- Every day for a week, add a mixture of triple antibiotic (neomycin, polymyxin B, and bacitracin) and chlorohexidine creams around the implantation area and under the nails. (**Optional: add lidocaine cream for gentle numbing (at all stages of cream ideally – but not too often, lidocaine has its issues namely toxicity with excessive use.**).
    - (The rats nails should be trimmed during surgery to prevent excessive damage from grooming at the edges of the implant)
    - Lubrisyn (hyaluronic acid) may be added to the implantation area once the tissue has mostly scarred over to further promote healing.

##### 4.2 Margin Care – 2-3 weeks after surgery

Margin care refers to caring for the margins of the implant's dental cement where the skin meets. Margin care is important to maintaining an infection-free site and prolonging the life of the implant and rat.

- Enrofloxacin (Enro) wash
- Benzocaine infusion
- Tweezer removal of gunk
- Enro wash 2
- Air Gun Drying
- Enro wash + silver nitrate stick (cauterization of open wounds; prevent bleeding, crusting, hair regrowth, skin regrowth)
- Saline wash to stop silver nitrate reaction
- Air Gun Drying 2
- Liquid Antibiotics + hyaluronic acid
- Air Gun Drying 3
- Optional: wait a few weeks and redo the procedure above before continuing, or:
    - Refill cement edges (only if necessary!)

### Supplemental Note 4: Materials and sourcing

| | Part | Quantity | Supplier | Supplier ID |
|---|---|---|---|---|
| **Fastenings** | M1 x 4 screw | 2 pack (pk) | McMaster Carr | 91430A151 |
| | M1.4 x 12 screw (0.3 mm pitch) | 1 pk. | | 91800A711 |
| | M1.4 x 4 screw | 2 pk. | | 91430A156 |
| | M6 x 15 screw (optional, if using aluminium breadboard) | 1 | | 90128A335 |
| | M1 threaded insert | 1 pk. | | 92120A110 |
| | M1.4 threaded insert | 1 pk. | | 92120A140 |
| | M1 nut | 3 pk. | | 90591A311 |
| | M2.6 x 25 screw | 1 pk. | | 90353A120 |
| | M2.6 nut | 1 pk. | | 90592A008 |
| **Hardware** | Drill bit 1.0 mm | 1 | McMaster Carr | 2958A25 |

| | | | | |
|---|---|---|---|---|
| | Drill bit 1.4 mm | 1 | | [2958A34]() |
| | Drill bit 7/64 in | 1 | | [2901A507]() |
| | Slotted screwdriver bit 2mm, with 4mm hex shank | 1 | | [5750A375]() |
| | Slotted screwdriver bit 1.5 mm, with 4mm hex shank | 2 | | [5750A373]() |
| | Drill | 1 | Grainger | |
| | Soldering iron | 1 | | [799RP8]() |
| | Solder | ? | | [19YP69]() |
| | Lubricant oil | 1 | | [41MW02]() |
| | Silicone glue | 1 | | [22N770]() |
| | Sandpaper | | | |
| | UV resin | | | |
| | Copper or copper | | | |

| | | | | |
|---|---|---|---|---|
| | alloy sheet | | | |
| | Plastic tip tweezers | 1 | DigiKey | 2014-2ACFR.SA.1.ITU-ND |
| | Aluminium breadboard (optional) | 1 | Thorlabs | MB3060/M |
| | Screwdriver | 1 | StarTech | CTKRPR (kit including slotted screwdriver bit) |
| **Electronics** | Pin connector male (x2 if 2 probes) | 30 | DigiKey | 350-10-101-00-006000 |
| | Socket connector female (x2 if 2 probes) | 30 | | 2-5331272-3 |
| | Silver or Stainless Steel PFA coated wire | 1 | A-M Systems | 786500<br>791600 |
| **Implant printable components** | Shuttle case | 1 | This paper | https://github.com/rjibanezalcala/EXPLORE/tree/main/IMPLANTS |
| | Skull interface | 1 | | |

| | | | | |
|---|---|---|---|---|
| | Probe shuttle | 1 | | |
| | Headstage interface | 1 | | |
| | Ribbon pincher | 2 | | |
| | Headstage interface cap | 1 | | |
| | Implant stereotactic holder | 1 | | |
| | Implant building aides | 1 | | |
| | Skull interface holding bay | 1 | | |
| | Shuttle elevator | 1 | | |
| **“Kepler” screwdriver printable components** | Bottom casing | 1 | This paper | https://github.com/rjibanezalcala/EXPLORE/tree/main/KEPLER |
| | Bottom casing shell | 1 | | |
| | Knob (small or large) | 1 | | |
| | Bit adapter | 1 | | |

| | | |
|---|---|---|
| | Middle indicator | 1 |
| | Top casing | 1 |
| | Top casing shell | 1 |
| | Carrier | 2 |
| | Carrier spacer | 2 |
| | Planet gear | 6 |
| | Planet gear spacer | 6 |
| | Sun gear shaft | 1 |
| | Sun gear socket | 1 |
| | Sun spacer | 2 |
| | Ring gear | 2 |
| | Counter ring gear (optional) | 1 |
| | Counter carrier (optional) | 1 |

| | | |
|---|---|---|
| | Counter carrier spacer (opt.) | 1 |
| | Counter planet gear (opt.) | 3 |
| | Counter planet gear spacer (opt.) | 3 |
| | Counter sun gear (opt.) | 1 |
| | Counter sun gear spacer (opt.) | 1 |
| | Counter knob (opt.) | 1 |
| | Counter adapter sun gear (opt.) *Replaces one regular sun gear* | 1 |
| | Counter face (opt.) | 1 |

| | | | | |
|---|---|---|---|---|
| | Counter shell (opt.) | 1 | | |
| | Top casing shell for counter module (opt.) *Replaces regular top shell* | 1 | | |
| **Other 3D printables** | Stereotactic implant holder | 1 | This paper | https://github.com/rjibanezalcala/EXPLORE |
| | Stereotactic shuttle holder | 1 | | |
| | Implant “holding block” | 1 | | |
| | Saline bath (optional) | 1 | | |
| **3D printer and accessories** | Tough 2000 photopolymer resin | | Formlabs | RS-CFG-TO20-01 |
| | Clear V4 photopolymer resin | | | RS-CFG-GPCL-04 |

| | | | | |
|---|---|---|---|---|
| | Resin tank V2.1 | | | RT-F3-02-01 |
| | Build platform | | | BP-F3-01 |
| | Form3 SLA 3D printer, wash, and cure | | | PKG-F3-SVC-COMPLETE |
| **Software** | Blender *Needed to make edits to implant* | | Blender | Blender 3.4.0 |
| | Preform *For printing using Formlabs 3D printers* | | Formlabs | Preform |
| | Spike GLX *If using PXIe acquisition system* | | Github | Spike GLX |
| | Open Ephys GUI *If using PXIe acquisition* | | Open Ephys | Open Ephys GUI |

| | | | | |
|---|---|---|---|---|
| | *system, alternative to SpikeGLX* | | | |
| | Bonsai *If using ONIX acquisition system* | | Bonsai RX | Bonsai |
| **Recording equipment** | Neuropixels 1.0 probe | | Neuropixels | PRB_1_4_0480_1 |
| | 1.0 headstage and cable *If using PXIe acquisition system (see below)* | | | HS_1000, CBL_1000 |
| | PXIe card *If using PXIe acquisition system (see below)* | | | PXIE_1000 |
| | PXIe-1082 Chassis | | National Instruments | https://www.ni.com/en-us/support/model.pxie-1082.html |

| | | | | |
|---|---|---|---|---|
| | ONIX PCIe Acquisition System | | Open Ephys | OEPS-9006 |
| **Other hardware** | A.M.P.I. *Iso-flex* Electrical stimulus isolator | | A.M.P.I.<br><br>Microprobes | Iso-flex<br><br>USA seller |
| | A.M.P.I. *Master-9* pulse stimulator | | A.M.P.I.<br><br>Microprobes | Master-9<br><br>USA seller |
| **Surgical supplies** | Metabond Kit | | Parkell | https://www.parkell.com/C-B-Metabond-Quick-Adhesive-Cement-System |

**Suppliers**

- McMaster Carr, USA (MMC)
- Digikey, USA (DK)
- Mouser Electronics, USA (ME)
- A-M Systems, USA (AMS)
- Open Ephys, Portugal (OE)
- IMEC, USA
- Grainger, USA
- Thorlabs, USA
- Formlabs, USA (FL)
- Neuropixels, USA
- Microprobes, USA
- AMPI, Israel

**Repositories**

- GitHub repository with 3D print files: https://github.com/rjibanezalcala/EXPLORE
- GitHub repository with signal processing algorithms: https://github.com/lddavila/cluster_neuronspikes/tree/main/Porting%20Open%20Ephys

Data repository: TBD
